# Origin and evolution of grapevine genomes

**DOI:** 10.64898/2026.04.20.719760

**Authors:** Xiangfeng Wang, Jie Sun, Jingxuan Wang, Xuenan Zhang, Shaoying Chen, Jingyun Jin, Xinhe Zhang, Falak Sher Khan, Ke Wang, Junjie Mei, Wei Zheng, Linling Guo, Haisheng Sun, Chonghuai Liu, Naomi Abe-Kanoh, Wenxiu Ye, Li Guo

**Author notes:** **Corresponding authors:** Li Guo. Wenxiu Ye., Naomi Abe-Kanoh.

## Abstract

Grapevines (*Vitis*) belonging to grape family (Vitaceae) are symbolic fruit crops pivotal to human civilization. The evolutionary history of grapevines divergent from other Vitaceae plants remains mysterious, requiring a family-wide whole-genome phylogenomic analysis. Here, we conduct chromosome-level phylogenomics to investigate the origin and evolution of grapevines using 29 genome assemblies of five genera *Vitis*, *Parthenocissus, Ampelopsis, Tetrastigma,* and *Cissus,* 27 of which are newly released in this study. Phylogenomic and macrosynteny analysis unanimously support *Ampelopsis* as a sister lineage to *Parthenocissus,* placing both closer to *Vitis*, with introgression and incomplete lineage sorting contributing to these relationships. Ancestral genome reconstruction delineates the major chromosome rearrangement events in Vitaceae karyotype evolution, highlighting the conserved karyotype in *Vitis* and the extensive karyotypic reorganization in *Tetrastigma* and *Cissus*. Pan-3D genome analysis highlights the contributions of structural variants (SVs) to the variation of A/B compartments and topologically associated domains (TADs), revealing a strong purifying selection of SVs at TAD boundaries. We further demonstrate that Helitron transposons drive the expansion and expression regulation of NLR immune-receptor genes in *Vitis*. Importantly, we discovered an NLR gene *VbRpv35* from wild grapevine *V. bellula* resistant to downy mildew (DM), whose heterologous expression in *V. vinifera* confers enhanced DM resistance. Taken together, we provide phylogenomic insight into the origin and evolution of grapevines and valuable resources for grapevine improvement and understanding angiosperm evolution.

## Introduction

Vitaceae, the grape family, harbors perennial plants including 16 genera and nearly 950 species^1^. Cultivated grapevine (*V. vinifera*) is the best-studied species in Vitaceae and one of the most economically important fruit crops with a long domestication history dating back to 11,000 years ago in Western Asia^2^. Despite their pivotal roles in human civilization, the evolutionary history of grapevines and Vitaceae has been elusive and intensively debated. Early studies based on plastid sequences have classified the family into five major tribes: Ampelopsideae, Cisseae, Cayratieae, Parthenocisseae and Viteae^1,3,4^, with Ampelopsideae proposed as the earliest diverging clade. A few studies have constructed phylogenetic trees based on selected low-copy genes^4^ and transcriptome data^5,6^. While these studies offer valuable insights, the full complexity of Vitaceae evolution remains uncaptured mainly owing to these datasets may introduce biases compared to whole-genome data. Furthermore, potential gene flows and hybridization among clades have posed severe challenges in an accurate resolution of species relationships and thus the elucidation of their evolutionary histories^7^. Above all, phylogenomic analysis of Vitaceae family has been impeded by the limited availability of reference genomes across all five tribes. Majority of released Vitaceae genomes come from *Vitis* genus, particularly *V. vinifera*^8–14^. So far, chromosome-scale genome assemblies of key evolutionary nodes such as Ampelopsideae and Parthenocisseae remain unavailable, hindering a comprehensive phylogenomic analysis for understanding Vitaceae macroevolution.

Chromosomal rearrangements following whole-genome duplication (WGD) profoundly influence evolutionary trajectories^15,16^ in plants. In several well-studied families, such as Malvaceae^17^ and Brassicaceae^18^, comparative analyses of chromosome-scale assemblies have revealed the ancestral chromosome configurations and lineage-specific rearrangements including fusions, inversions, and translocations that underlie family diversification, and have demonstrated that chromosomal structural data can resolve phylogenies of ancient hybrid or polyploid genomes. In *V. vinifera*, the history of chromosomal rearrangements has been investigated through ancestral eudicot karyotypes (AEK) and ancestral core eudicot karyotype (ACEK) analysis^19^. Notably, Vitaceae plants exhibit significant karyotype variations, where grapevines (*Vitis*) typically have 19 or 20 haploid chromosomes, while the number of chromosomes in *Ampelopsis* spp. and *Parthenocissus* spp. ranges from 20 to 40. Furthermore, the tropical genera of Vitaceae display even greater karyotype variation, with 12 or 24 haploid chromosomes in *Cissus* spp., 20 to 60 in *Cayratia* spp., and 11 to 26 in *Tetrastigma* spp.^20^. Recent studies show that *T. hemsleyanum*^21^ and *C. rotundifolia*^22,23^ exhibit substantial karyotype differences compared to *V. vinifera*. Chromosomal rearrangements have generated karyotype variation in Vitaceae, but the processes underlying these rearrangements remain largely uncharacterized. With a relatively simple genome duplication history (lacking post-γ WGD events)^21–24^, it also provides an ideal reference for investigating plant WGDs and chromosomal rearrangements through phylogenomics-based ancestral chromosome reconstruction from chromosome-scale assemblies. Chromosome-level genomes are only reported for Cayratieae, Cisseae, and Viteae to date, missingAmpelopsideae and Parthenocisseae phylogenetically distinct and morphologically diverse tribes^25^. This gap limits reconstructing a complete evolutionary framework for Vitaceae, resolve karyotype evolution, and identifying lineage-specific genomic innovations. Chromosome-scale genomes for these two tribes are therefore essential to provide a comprehensive reference set for family-wide comparative genomics.

Genome evolution is driven by genomic variations which can significantly influence phenotypic traits for species adaptation, such as disease resistance, as reported in numerous studies^26–29^. Wild grapevines typically harboring a richer repertoire of resistance genes are more disease-resistant than the cultivated grapevines^8,9,30,31^. However, the evolutionary mechanisms behind resistance gene gain and loss remain largely unknown in grapevines. In plants, nucleotide-binding leucine-rich repeat (NLR) genes encode intracellular immune-receptors that recognize pathogen effectors to trigger host immune responses^32^. The number of NLR genes varies greatly among species, even among closely related species, due to rapid gene gain and loss^33–35^. This variation reflects species adaptation to pathogens and affected by different selection pressures^32^. To date, pan-NLR analyses were conducted in various species^36^, including Solanaceae^37^, Arabidopsis^38^, rice^39^, grapevine^8,30^, melon^40^, potato^37^, and wheat^41^, providing insights into the rapid changes of NLR repertoire (NLRome) and composition in plants. Comprehensive genome-wide analysis of NLRs relies on high-quality genome assemblies. However, many plant families still lack a sufficient number of representative genomes. As a result, current pan-NLRome studies are mainly conducted within single species or limited genera^38–40,42^. Pan-genome studies involving NLRs have largely emphasized gene number variation and NBARC sequence comparisons, while other aspects of their evolutionary dynamics remain less explored. At present, in-depth NLR analyses remain lacking across families such as Vitaceae. Moreover, copy number variation in NLRs may contribute to the variable disease resistance, yet the mechanisms underlying such numerical differences are still poorly understood.

Helitrons represent a distinct class of transposable elements (TEs) widespread across eukaryotic genomes. Unlike many other DNA transposons, Helitrons transpose via a rolling-circle mechanism, often retaining the original copy-a process characterized as “peel-and-paste”^43,44^. A notable functional feature of Helitrons is their propensity to capture and mobilize host genomic fragments^45–47^. This ability to acquire genomic material facilitates segmental duplications, structural rearrangements, exon shuffling, the formation of chimeric transcripts, and the dissemination of regulatory elements. Notably, when the termination signal is impaired, such as those lacking a functional hairpin structure at the 3’ end, Helitrons may exhibit enhanced capture activity^44^. In maize, Helitron activity is particularly pronounced constituting approximately 4% of the genome^48^. A substantial proportion of Helitrons carry typically one to three gene fragments, although some contain fragments from up to twelve distinct genes^47–49^. Their preferential localization in gene-rich regions further underscores their impact on genomic architecture and evolution^50^. Helitrons are found to capture and duplicate NLR genes in Solanaceae, contributing to lineage-specific expansion of NLR repertoires^51^. Therefore, Helitrons are key drivers of genomic innovation and diversity of eukaryotic genomes, and the mechanism underlying this process demands a better understanding from large-scale evolutionary genomic analyses.

Here, we generated chromosome-level genome assemblies for 27 Vitaceae species, including the first reference genome for two tribes, Parthenocisseae (*Parthenocissus tricuspidata*) and Ampelopsideae (*Ampelopsis glandulosa*). We constructed the first phylogenetic tree of Vitaceae based on nuclear genomic data from five tribes, revealing the evolutionary relationships within the family. Comparative analysis of SV frequencies at TAD boundaries and within TADs suggested a purifying selection of SVs at TAD boundaries. Additionally, correlation analysis showed a strong linkage between NLRs and Helitrons in Vitaceae. Furthermore, we presented evidence that Helitrons drove the proliferation of NLRs within the Vitaceae family. Finally, we identified a new NLR *VbRpv35* from wild grape *V. bellula* resistant to DM, and showed that its heterologous expression in *V. vinifera* conferred enhanced DM resistance. These findings provide novel and comprehensive insights into the evolutionary history of Vitaceae and serve as a valuable genomic resource for accelerating advanced breeding and cultivation in viticulture.

## Results

### Chromosome-level assembly and annotation of the 27 Vitaceae genomes

To assemble a representative set of Vitaceae genomes for phylogenomic investigation of grapevine evolution, we generated PacBio high-fidelity (HiFi) sequencing (mean coverage: 79×) and high-throughput chromatin conformation capture sequencing (Hi-C) sequencing reads (mean coverage: 144×) for 26 Vitaceae species. This dataset includes 24 *Vitis* species, one Boston Ivy (*P. tricuspidata*), and one Creeper (*A. glandulosa*), representing 26 new accessions and 9 new species (**Supplementary Table 1**). The HiFi reads were assembled with hifiasm^52^ followed by de-duplication using Purge-dups^53^, and contaminated sequences from microbes and plastids were removed to yield initial contigs. The resulting draft assemblies consisted of 37 to 217 contigs, with contig N50 ranging from 12.83 to 30.27 Mb (mean: 22.57 Mb) (**Supplementary Fig. 1 and Supplementary Table 2**). The draft assemblies were further anchored onto chromosomes using the Hi-C reads (**Supplementary Table 1 and Supplementary Fig. 2-3**) with Juicer^54^ and 3d-DNA pipeline^55^, followed by manual check and correction for errors in Juicebox^56^, yielding the chromosome-level assemblies with an average anchor rate of 98.09% (**Supplementary Table 2**). Additionally, sequencing reads of *V. vinifera* cultivar Chardonnay were obtained from public dataset^6^, including 156.9 Gb HiFi and 91.7 Gb Hi-C reads, and 58.9 Gb Oxford Nanopore (ONT) ultra-long reads (N50 > 100 kb). The *V. vinifera* genome assembly was conducted using both ONT and HiFi reads, followed by the de-duplication, contamination removal, and polishing using long reads, obtaining a gap-free assembly. The 27 genome assemblies all contained 19 chromosomes in 24 *Euvitis* species except for *V. rotundifolia* (Muscadine), *P. tricuspidata*, and *A. glandulosa* having 20 chromosomes, consistent with previous karyotyping reports for Vitaceae^20^. Notably, the genomes of *P. tricuspidata* and *A. glandulosa* represented the first chromosome-level assemblies released for *Ampelopsis* and *Parthenocissus*, expanding genomic resources for the grape family. Hi-C interaction maps showed no obvious misplaced contigs in these genome assemblies, indicating their high structural accuracy. In addition, validations using BUSCO^57^, merqury^58^ and LTR_retriever^59^ yielded scores of 97.5% to 98.8% (mean: 98%), quality values of 62 to 71 (mean: 66), and LTR Assembly Index (LAI) values of 13 to 21 (mean: 18) (**Fig. 1b and Supplementary Table 2 and Fig. 4**), indicating the high quality and accuracy of the assemblies. For genome annotation, we employed *ab initio* predictions combined with homologous proteins and transcriptomic evidence, identifying 30,221 to 40,280 protein-coding genes with a mean BUSCO score of 94.65% (**Supplementary Table 3**), reflecting the high completeness of these genome assemblies.

**Fig. 1.**
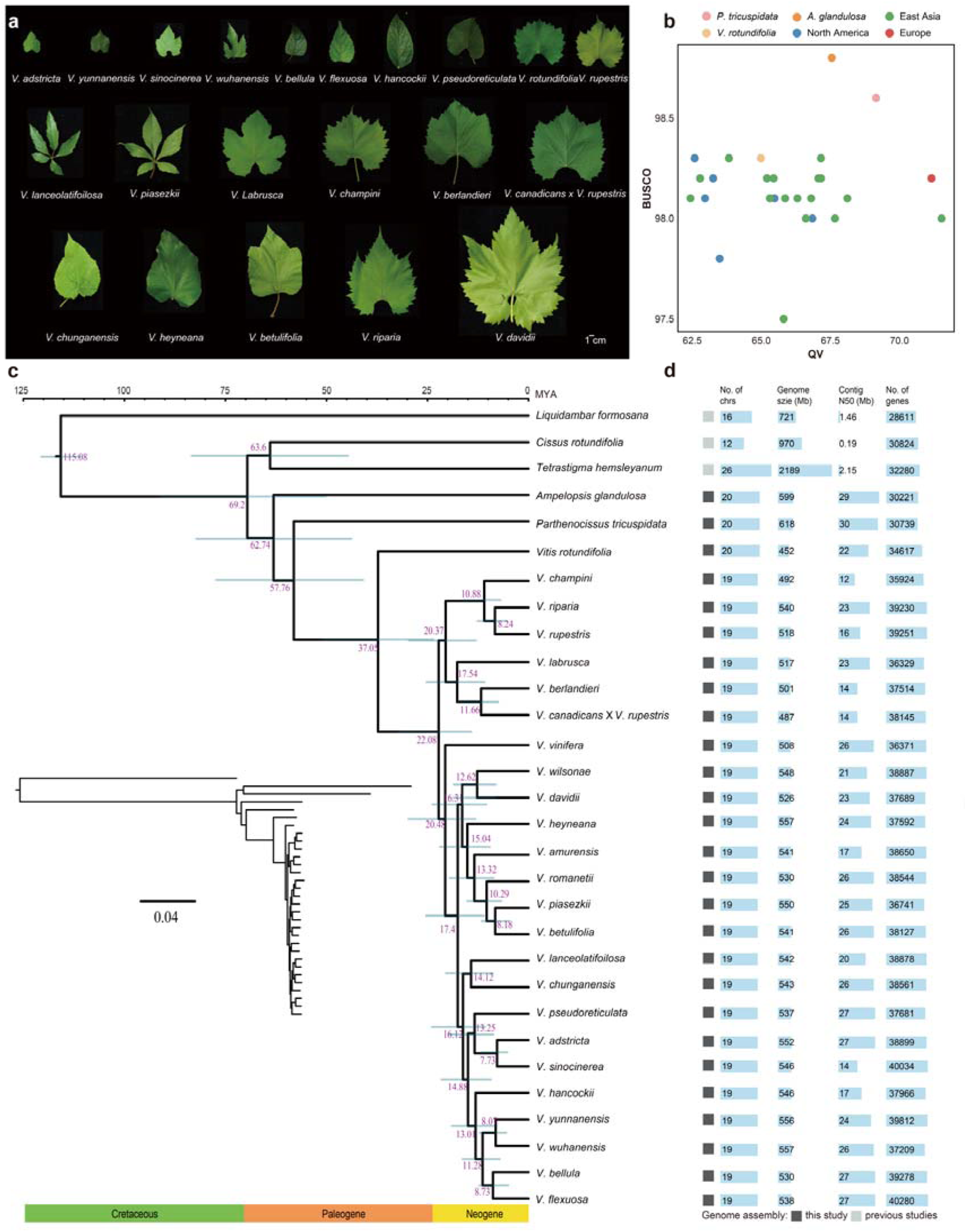
Phylogenomic analysis of Vitaceae plants spanning ∼70 million years of evolution. **a,** Leaf morphologies of *Vitis* accessions used in this study. **b,** BUSCO and QV metrics across the Vitaceae genomes. Colors represent Viticeae (excluding *Vitis*) and *Vitis* accessions categorized by different geographical origins. **c,** Maximum-likelihood tree of the concatenated 2,365 SCOs from the 30 species. The phylogenetic timescale (million years ago, MYA) is shown at the top, with node divergence points labeled in purple and 95% confidence intervals represented by blue horizontal bars. **d,** Assembly metrics of the genomes in (**c**).

### Vitaceae phylogenomics reveals gene flow and incomplete lineage sorting

To investigate the evolutionary history of Vitaceae, we constructed a family-level phylogenomic tree based on 2,365 single-copy orthologs (SCOs) from 30 species, including 27 newly assembled genomes, two previously published Vitaceae genomes (*C. rotundifolia*^22^ and *T. hemsleyanum*^21^), using *Liquidambar formosana*^60^ as the outgroup. Divergence time estimation using MCMCTREE, with fossil calibrations informed by TimeTree^61^, placed the split of Vitaceae into two major clades at ∼69.2 million years ago (Mya) (**Fig. 1c**). At *∼*63.6 Mya, one clades split into *C. rotundifolia* and *T. hemsleyanum*, while the other clade diverged into *A. glandulosa* at *∼*62.7 Mya and subsequently split into *P. tricuspidata* and *Vitis* at ∼57.8 Mya. Gene family contraction and expansion analyses across different clades were performed using CAFE^62^. *T. hemsleyanum* exhibits a pronounced expansion of gene families, suggesting a potential for functional innovation associated with its increased genome size compared with other genomes (**Supplementary Fig. 5a**). KEGG analysis revealed that all Vitaceae species show contraction of gene families related to primary metabolism, reflecting an evolutionary strategy to maintain essential biological functions while enhancing metabolic efficiency and adaptability by reducing redundancy. Meanwhile, different species have independently evolved with expanded related gene families in specific secondary metabolism such as terpenoids, flavonoids, and glucosinolates, likely contributing to defense mechanisms and ecological adaptation strategies (**Supplementary Fig. 5b-c; 6**).

Previous studies report that *A. glandulosa* is the earliest diverging lineage in Vitaceae based on plastid genes^3,4^. To resolve this discrepancy regarding *A. glandulosa* phylogeny, we constructed 2,365 species trees using each SCO alignment and obtained a coalescent-based phylogenetic tree with ASTRAL^63^. As a result, 49.17% of the phylogenetic trees placed *A. glandulosa* close to *P. tricuspidata*, while 41.18% supported a distant phylogenetic relationship (**Supplementary Fig. 7**). Additionally, 55.71% of trees supported that *A. glandulosa* forms neighboring lineages with *P. tricuspidata* and *V. vinifera*, while 39.54% supported *P. tricuspidata* and *V. vinifera* form neighboring lineages with *C. rotundifolia* and *T. hemsleyanum* (**Supplementary Fig. 7**). Therefore, phylogenomic using both concatenated alignement and coalescent approach supported the more likely placement of *Ampelopsis* close to *Parthenocissus* and *Vitis*.

The minority but substantial amount of trees supporting early divergence of *Ampelopsis* suggested potential gene introgressions and/or incomplete lineage sorting (ILS) of ancestral polymorphisms in Vitaceae evolution. To test this hypothesis, we first performed ABBA-BABA analysis and estimated Patterson’s *D*-statistics and *f*-branch scores to detect gene flows. Patterson’s *D*-statistics detected the excess of shared derived alleles between *Vitis* and *A. glandulosa,* between *Vitis* and *T. hemsleyanum*, as well as between *A. glandulosa* and *T. hemsleyanum*, supporting the presence of gene flows (**Fig. 2a, Supplementary Table 4**). In addition, the *f*-branch score (*f*_b_ = 0.2132) further indicated extensive gene flow between *Vitis* and *A. glandulosa* (**Fig. 2b**). Since frequently shared alleles among non-sister lineages are probably caused by interspecific hybridization^64^, we calculated Quantifying Introgression via Branch Lengths (QuIBL) to differentiate introgression from ILS^64,65^. The calculation indicated a significant fit of the tree topology to the ’introgression+ILS’ model (**Supplementary Table 5**), supporting the presence of both ILS and introgression between *A. glandulosa* and *C. rotundifolia* as well as between *T. hemsleyanum*. Furthermore, whole-genome alignment between *A. glandulosa* and *P. tricuspidata* showed a strong collinearity without any large-scale chromosomal rearrangements, consistent with their gene-based synteny (**Fig. 2c**). The genome synteny between *A. glandulosa* and *P. tricuspidata* was significantly higher than those of between *A. glandulosa* and *C. rotundifolia* or *T. hemsleyanum* (**Fig. 2c**), supporting the phylogenetic position of *A. glandulosa* which closer to *P. tricuspidata* than to *C. rotundifolia* or *T. hemsleyanum*. The 1:1 inter-chromosomal collinearity between *C. rotundifolia*, *T. hemsleyanum*, *P. tricuspidata*, and *A. glandulosa* with *V. vinifera* suggests that these five species, and more broadly the Vitaceae family, have not undergone any post-γ WGD event (**Fig. 2c**). Collectively, the construction of the Vitaceae phylogenetic tree demonstrated a comprehensive and complex evolutionary history of Vitaceae, with gene introgression and ILS being among the contributing factors.

**Fig. 2.**
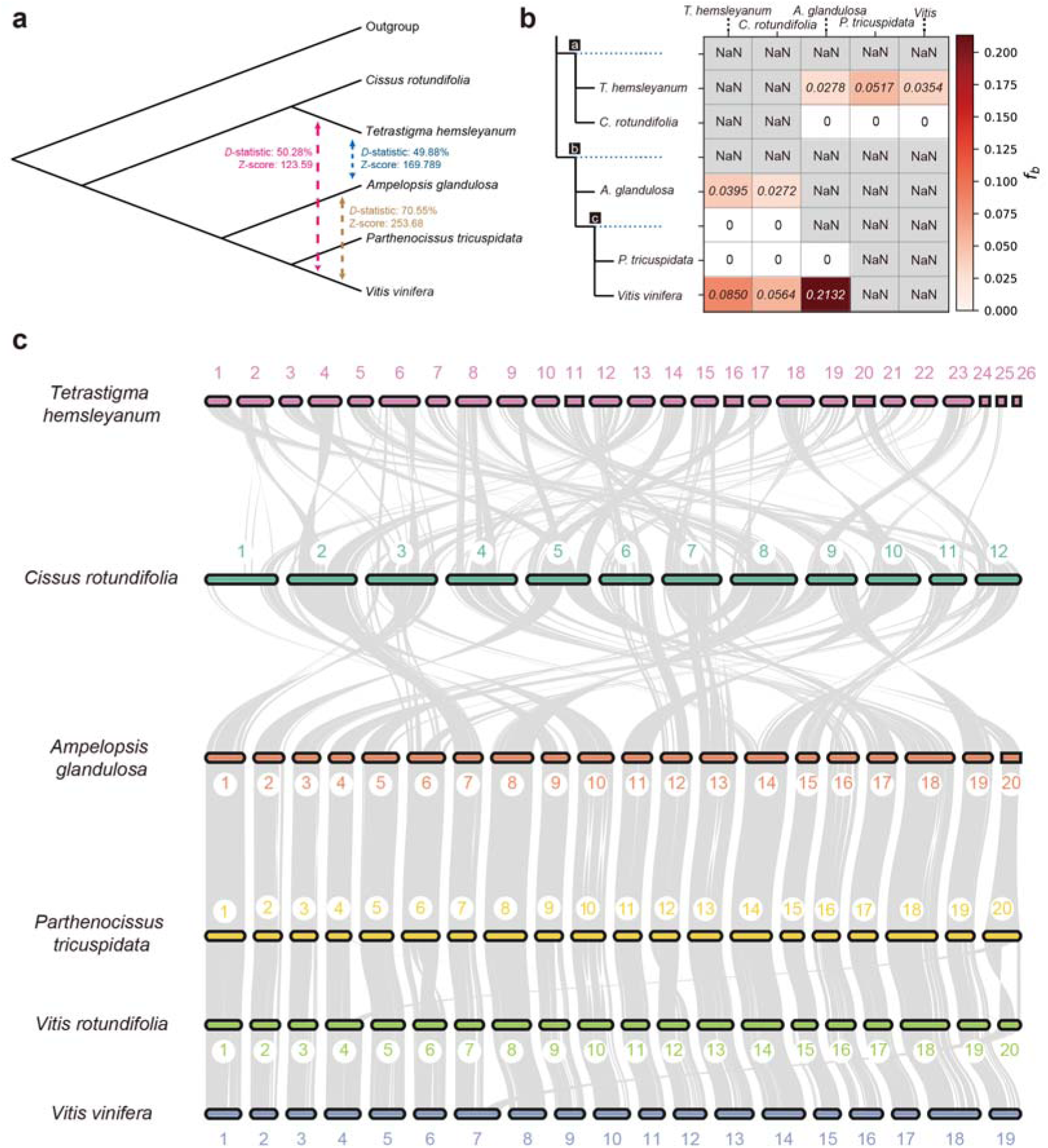
Genome evolution of major lineages in Vitaceae. **a**, ABBA-BABA analysis among the Vitaceae exhibiting the *D-*statistics and Z-score of gene flow. **b,** D-statistics analysis of gene flow among Vitaceae species branches and internal tree nodes (a, b, c on branches). *f*_b_ indicates the f-branch score, and NaN represents ’Not a Number’. **c**, Genomic collinearity analysis among six Vitaceae species. The numbers represent chromosome numbers, and the gray lines indicate collinear blocks.

### Reconstruction of Vitaceae ancestral genomes

Chromosomal rearrangements play a crucial role in karyotype diversification and species divergence. Diverse karyotypes (n=19, 20, 26 etc.) in Vitaceae highlight a series of complex chromosome rearrangements occurred during speciation^20^. Although many studies reported ancestral karyotypes in plants^17,66–69^, ancestral Vitaceae genomes and the chromosome rearrangement shaping the diverse Vitaceae karyotypes have not been reconstructed. Here, we reconstructed the ancestral genomes of the Vitaceae from the chromosome-level genomes of representative Vitaceae species, along with shared chromosomal synteny blocks (CLSBs) identified via the WGDi pipeline^67^ (**Fig. 3a**). We selected six accessions representing distinct karyotypes across the five major tribes of Vitaceae. Among them, *V. vinifera* serves as the representative of the subgenus Euvitis (**Supplementary Table 1**), whose members share a conserved karyotype^8^. Because the karyotypes of *C. rotundifolia* and *T. hemsleyanum* differ substantially from those of other species (**Fig. 2c**), we did not attempt to infer ancestral Vitaceae karyotypes (AVK) directly from a set of representative extant species. Instead, we inferred ancestral karyotypes at internal nodes in a bottom-up manner, based on the karyotypic structures of extant genomes. Using this approach, we successively reconstructed the ancestral Viteae karyotype (V, *n* = 20), the ancestral Viteae + Parthenocissae karyotype (VP, *n* = 20), the ancestral Viteae-Parthenocissae-Ampelopsideae karyotype (VPA, *n* = 20), and AVK (*n* = 21) (**Supplementary Fig. 10-14**). Chromosomal rearrangement events were then identified along the evolutionary trajectory from ACEK through AVK to the extant species (**Fig. 3; Supplementary Fig. 15-19**). The reduction from 21 protochromosomes in AVK to 19 chromosomes in Euvitis likely involved two end–end joining (EEJ) events, AVK/14_21/EEJ and V/7_20/EEJ, resulting in stepwise decreases in protochromosome number (**Fig. 3**).

**Fig. 3.**
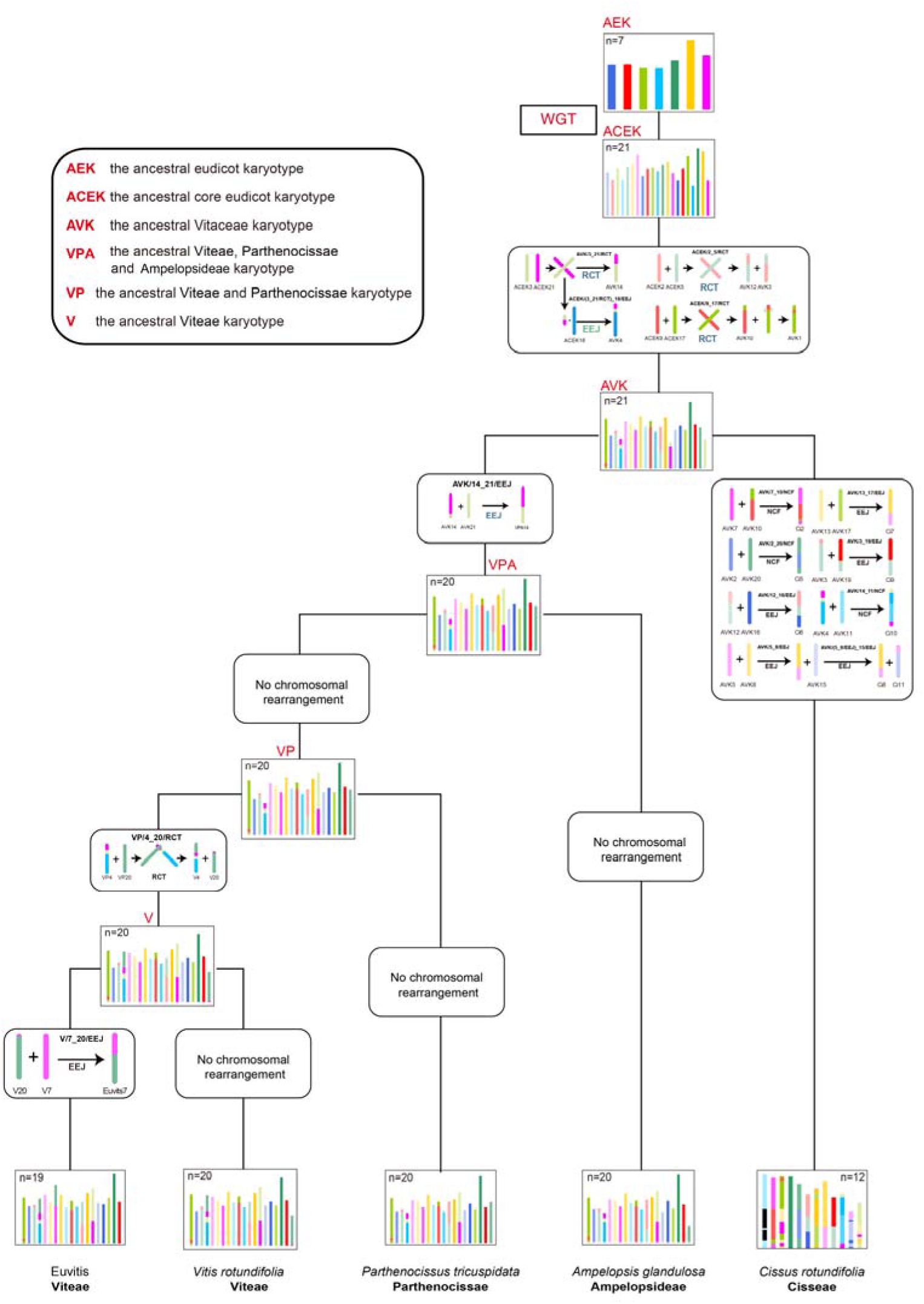
Vitaceae ancestral genome reconstruction reveals karyotype evolutionary history and mechanism. Schematic diagram illustrates the karyotype evolution of five representative tribes of Vitaceae, inferred from chromosomal rearrangements. Colored blocks on progeny chromosomes correspond to homologous chromosomes in the ACEK, depicting chromosomal changes during the evolution from ACEK to extant species. WGT represents the whole-genome duplication event common to core eudicots. "n" represents the number of chromosomes. Nested Chromosome Fusion(NCF), Reciprocal Chromosome Translocation (RCT), End–End Joining(EEJ).

Interestingly, although *A. glandulosa* and *P. tricuspidata* diverged approximately 20–25 million years earlier than *Vitis* (**Fig. 1c**), *A. glandulosa* and *P. tricuspidata* exhibited identical karyotypes with only a Reciprocal Chromosome Translocation (RCT, VP/4_20/RCT) event distinguishing the two species from *Vitis* spp. (**Fig. 3; Supplementary Fig. 12b**). In contrast, while diverged earlier than *Vitis* at approximately 35 mya (**Fig. 2c)**, *T. hemsleyanum* and *C. rotundifolia* showed pronounced karyotypic differences compared to *Vitis* (**Fig. 3 and Supplementary Fig. 17 and 18**). Moreover, apart from rearrangements inherited from the ACEK-to-AVK transition, no additional chromosomal rearrangement events were shared between *T. hemsleyanum* and *C. rotundifolia* (**Fig. 3 and Supplementary Fig 20**).

Because shared chromosomal fusion events can provide robust evidence for phylogenetic relationships^18^, we systematically compared *A. glandulosa* with other species to clarify its phylogenetic position within Vitaceae (**Supplementary Table 6**). This analysis revealed one shared EEJ event (AVK/14_21/EEJ) among *A. glandulosa*, *P. tricuspidata*, and *Vitis* (**Supplementary Fig. 20**). By contrast, no additional shared rearrangements were detected between *A. glandulosa* and either *T. hemsleyanum* or *C. rotundifolia* beyond those occurring during the ACEK-to-AVK transition (**Fig. 3**). Collectively, these results not only reconstructed the ancestral Vitaceae karyotype and the trajectory of karyotype evolution from AVK to extant lineages, but also were consistent with the phylogenomic tree inferred from single-copy orthologs, all supporting a closer phylogenetic relationship of *A. glandulosa* with *P. tricuspidata* and *Vitis* than with *T. hemsleyanum* and *C. rotundifolia*.

### Landscape of transposable elements and structural variants across Vitaceae

TEs are widespread repetitive sequences that substantially contribute to plant genome evolution, such as long terminal repeats retrotransposons (LTRs), typically dominating repetitive sequences of plant genomes^70–72^. To investigate the association between TEs and Vitaceae evolution, we annotated the TEs of the 29 Vitaceae genomes using RepeatModeler^73^ and RepeatMasker^74^ pipelines. TEs accounted for 42.2 to 55.4% of their genomes, with LTRs standing out as the most abundant TEs (**Supplementary Table 7**), showing unique composition preferences among species (**Fig. 4a**). However, certain types of TEs showed species-specific patterns. For example, *V. rotundifolia* exhibited a significantly higher proportion of hAT elements compared to others, while LINE ratio was considerably lower in *A. glandulosa*. We further classified intact LTRs into 16 subfamilies using TEsorter^75^, most of which were similarly distributed across the five tribes (Viteae, Ampelopsideae, Parthenocisseae, Cisseae and Cayratieae) (**Supplementary Fig. 21a**). However, significant differences in specific LTR abundances were observed among tribes, with *Ale* being the most abundant in Viteae, while *SIRE*, *Tekay*, and *Angela* dominated in *A. glandulosa*, *T. hemsleyanum*, and *C. rotundifolia*, respectively (**Supplementary Fig. 21a**), reflecting lineage-specific dynamics of LTR retrotransposon expansion within the Vitaceae. Pan-TE analysis across *Vitis* species of different geographic distributions^76^ revealed a weak correlation (Pearson’s correlation) between genome sizes and TE sizes. However, a positive correlation between LTR length with genome sizes was observed with an outlier in *Ale* and *Athila* varying within and among subfamilies and clades (**Fig. 4b and Supplementary Fig. 21b, 22**). These findings underscore a strong association between genome sizes and specific LTR subfamilies, with different roles played by distinct families likely contributing to the speciation of these species.

**Fig. 4.**
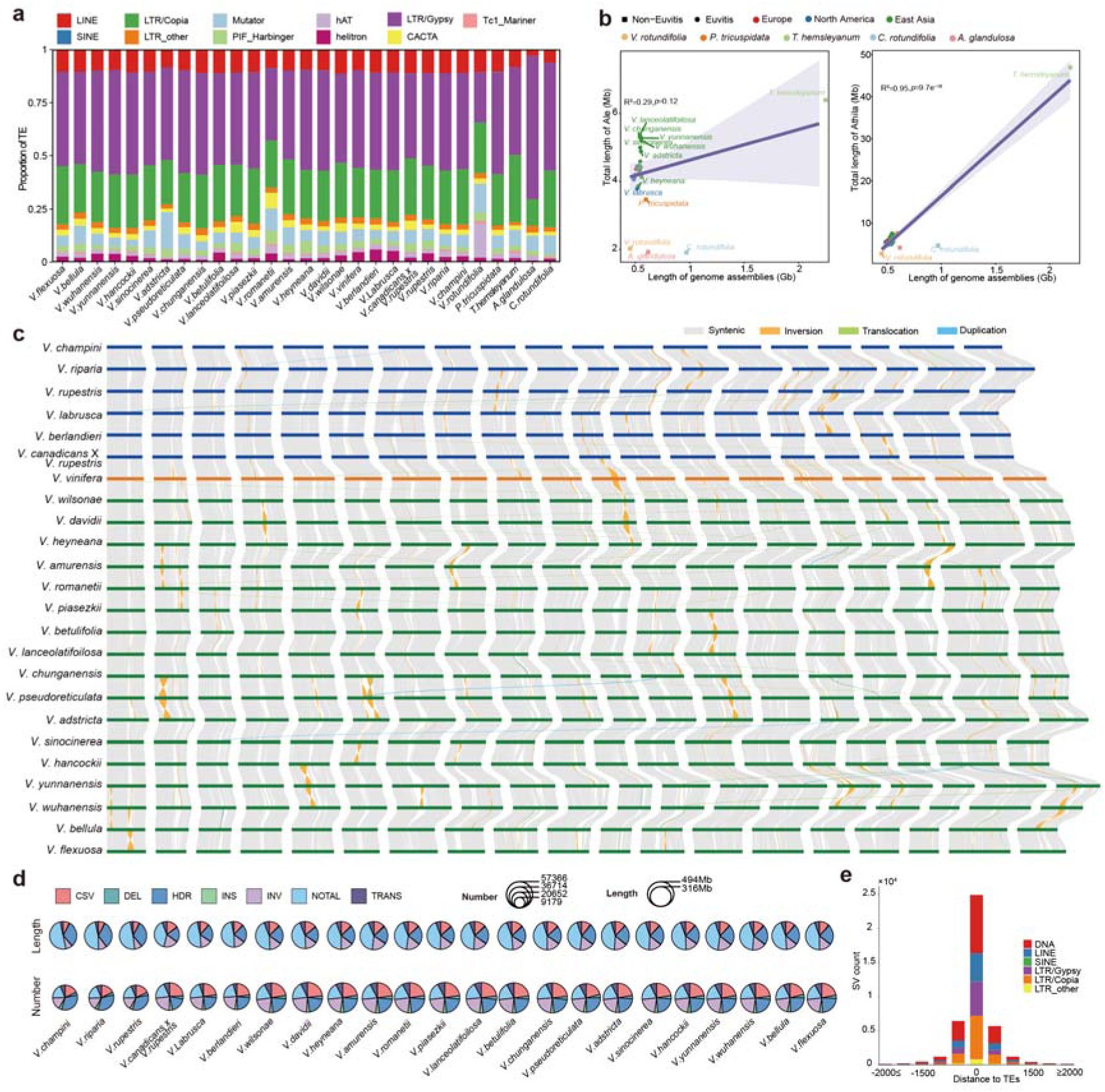
Landscape of transposable elements (TEs) and structural variants (SVs) across the Vitaceae. **a**, TE compositions of the Vitaceae genomes. Different colors represent different types of TEs. **b,** Correlation of LTR/Ale and LTR/Athila with the genome sizes. R² and *p* represent the goodness of fits and significances for the regressions, respectively. **c,** Whole-genome syntenies across the 24 Euvitis genomes illustrating syntenic regions and three SV types (Inversion, Translocation and Duplication). **d,** The SV profiles of the number and length across different SV types (CSV: Complex structural variants, DEL: Deletion, HDR: Highly divergent regions, INS: Insertion, INV: Inversion, NOTAL: Not aligned region, TRANS: Translocation). **e**, Distribution of SVs relative to TE distance. Bar colors represent the types of TEs proximal to SVs.

TE insertion is a major contributor to genome variants among species and individuals, likely coupled with functional consequences in evolution^77–80^. Using the assembly of *V. vinifera* as a reference genome, we systemically identified large-scale genomic variants in grapevines by genomic alignment of the 24 *Vitis* species followed by structural variation (SVs) detection using SyRI (**Fig. 4c**). This has detected a total of 330,292 simple SVs including 41,722 insertions, 43,441 deletions, 245,129 inversions, 84,650 translocations and 259,869 complex SVs. Interestingly, grapevines of Asian origins contained an overall higher number of SVs than those of North American origins, although the total length of variants in different types was comparable between the two groups based on comparisons with the *V. vinifera* reference genome (**Fig. 4d**). Our analysis revealed that chromosome arms exhibited high synteny across all genomes without large-scale chromosomal rearrangements. Large insertion/deletion polymorphisms were absent from chromosome arms, with the vast majority of SVs being smaller than 100 kb, and the largest reaching approximately 770 kb (**Supplementary Fig. 23**). While the number and size of SVs showed little variation among species, extensive 275,156 high-density recombination (HDR) regions were detected when compared to the *V. vinifera* reference genome. We also detected 227,279 NOTAL (unaligned sequences) events, reflecting the sequence divergence among *Vitis* species. We found that a subset of SVs were located in close proximity to retrotransposons and DNA transposons (**Fig. 4e**), suggesting that some of these SVs may have originated through TE transposition, a phenomenon also observed in other species^81^. Although DNA-type TEs accounted for only ∼10% of the total genome size in each *Vitis* species (**Fig. 4a and Supplementary Table 7**), their high copy number made them one of the primary drivers of SV formation (**Fig. 4e**). A number of SVs were located within gene regions, predominantly in introns, while very few affected exons (**Supplementary Fig. 23**).

### Pan-3D landscape shows significant 3D structure variations among *Vitis spp*

Eukaryotic genomes fold into three-dimensional (3D) structures such as A/B compartments and TADs, that organize the spatial genome into distinct regulatory territories^82^. However, it should be noted that plant genomes may form compartment-like and TAD-like structures through different mechanisms compared to animals^83^. To understand the diversity and functional implications of the 3D genomic structure of grapevines during evolution, we conducted a pan-3D genome analysis of 25 *Vitis* genomes using mapping Hi-C reads to each assembly. First, pan-A/B compartment analysis revealed that a relatively small proportion of the genomic bins (24.3%; Conserve A: 14.1%; Conserve B: 10.1%) exhibited conservation (**Fig. 5a,b**). By contrast, 67.3% of the compartments displayed variability, indicating extensive compartment switching owing to species divergence. The 3D genome structure and its evolution profoundly influence the functional divergence of genes^84,85^. The variable A/B compartments were enriched with genes associated with secondary metabolite biosynthesis, such as flavonoid production and immune responses (**Supplementary Fig. 23a**), reflecting their involvement in the adaptation of grapevines to environmental pressures. In contrast, the conserved A/B compartments were enriched with genes associated with basic biological functions as expected (**Supplementary Fig. 23b, 25**). Second, we identified TADs in 25 grapevine genomes and examined the distribution of TAD boundaries across different types of A/B compartments. We observed that genomic compartments undergoing A/B switch harbored a greater number of TAD boundaries than those without switching, suggesting large-scale compartments often having larger TADs (**Fig. 5c**). Moreover, A compartments had much more TAD boundaries than B compartments **(Fig. 5c),** a pattern also noted in the soybean 3D-genome studies which showed that compartmentalization may be a key factor driving TAD formation^84,85^. Third, we performed pan-TAD analysis and identified core, dispensable and private TADs by comparing TAD boundary positions within genome-wide synteny blocks. Unlike soybeans, the majority of TAD boundaries were dispensable across *Vitis* genomes (**Fig. 5a**), whereas conserved core boundaries were rare (11), mostly located within variable A/B compartments. This was likely caused by the species divergence in our genus-level analysis compared to the soybean study using cultivars. Despite variable A/B compartments dominating grapevine 3D genomes, dispensable boundaries were more frequently located within conserved A/B compartments, while unique boundaries were more likely to occur in conserved or dispensable B compartments (**Fig. 5d**). This implies that the evolutionary trajectories of A/B compartments and TADs appear to be independent of unknown mechanisms.

**Fig. 5.**
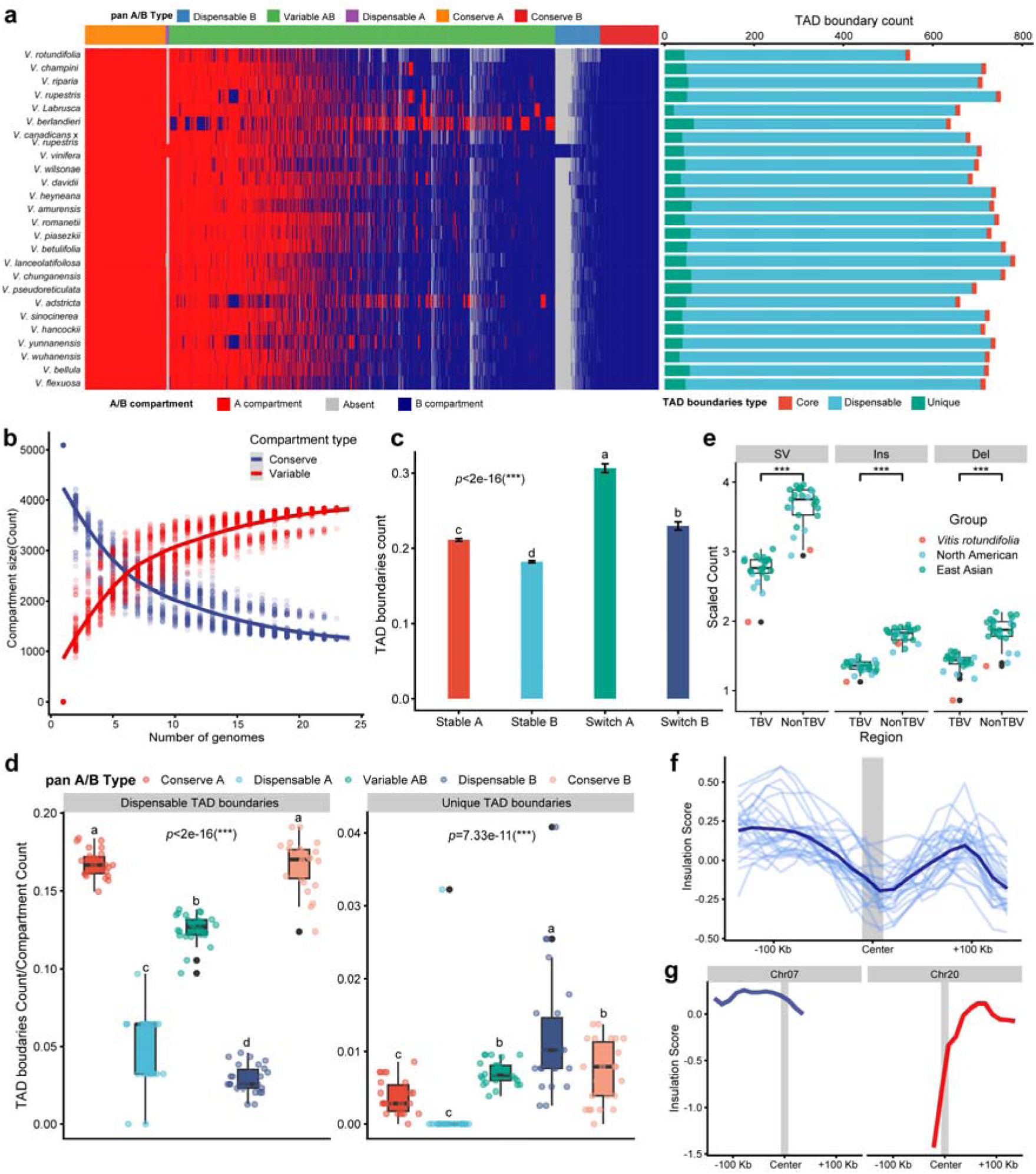
Pan-3D genomic landscape and evolution of grapevines. **a,** Pan-A/B compartment and TAD boundaries landscape of the 25 *Vitis* accessions. Absent represents bins with the 0 value of E1. E1 represents the first eigenvector from PCA of the normalized Hi-C interaction matrix, generally distinguishing A/B chromatin compartments. **b,** Profiling of compartment types across the 25 *Vitis* accessions. **c,** Distribution of TAD boundary in different A/B compartments. Letters indicate distinct significance levels. **d,** Distributions of Dispensable and Unique TAD boundaries count ratio in different pan A/B compartment types. The color of the dots represents different pan A/B compartment types. **e,** Average of SV number on TBV and non-TBV regions with their flank regions (±50kb) in each species. Different colors points represent the different group of species. **f,** Insulation score landscape spanning the flanks (±100 kb) of Chr7 fusion site across diverse *Euvitis* species. The navy line represents the mean insulation score, and cornflower blue lines represent individual species. The gray shaded region represents the fusion interval. **g,** Insulation score landscapes of the flanks(±100 kb) of orthologous pre-fusion regions on Chr7 and Chr20 of *Vitis rotundifolia*. The gray shaded region represents the fusion interval.

Although SVs can impact genome function via affecting 3D genome organization^84,85^, pan-genome level analysis of such effects remains limited but would offer mechanistic insight into the 3D genome evolution. Leveraging the high-quality *Vitis* genomes, we first identified 17,603 genome-wide TAD boundary variations (TBVs) and classified them into two types based on two major SV types causing the variations: insertions (Ins) and deletions (Del). Most of these TBVs were inside syntenic regions and few were found in unaligned regions given genomic differences. We observed that TAD boundaries with variations contained significantly fewer insertions and deletions than those without variations (**Fig. 5e**). This discrepancy is likely caused by an unbalanced SV distribution (**Supplementary Fig. 26a-b**), suggesting that mechanisms beyond SVs are involved in TBV formation. To understand how SVs located within TAD boundaries regions affect them, we examined their enrichment patterns at TBV regions. We found no preferential enrichment for either insertions or deletions across the different types of TBVs (**Supplementary Fig. 26c**). This suggests that both SV types may contribute similarly to TAD boundaries variations, a pattern consistent with reports in soybeans^62^. Our pan-3D and SV analysis in grapevines highlighted the positive roles of genome sequence variations in 3D organization of grape genomes, with implications in the adaptation of grapevines. Furthermore, at the fusion site on Chr7 (derived from ancestral Chr7 and Chr20), we observed a distinct valley in the insulation score^86^(**Fig 5f**). The insulation score quantifies the aggregate of chromatin interactions occurring across genomic intervals, and local minima indicate regions of high insulation that correspond to TAD boundaries^86^. A similar pattern was observed at the end of Chr7 and the start of Chr20 in *V. rotundifolia* (**Fig 5g**), indicating that TAD boundary was well maintained without visible fusions.

### Pan-NLRome analysis reveals chromosome-preference for NLR gene evolution

Novel traits in species’ evolution are often manifested at the genetic level. Cultivated grapevines suffer from devastating diseases such as downy mildew and grey molds due to domestication and breeding, and resistant cultivars are highly desired for viticulture. NLR encode immune-receptors playing key roles in plant immunity as R proteins. Although NLR genes in grapevines are reported by several studies^8,30^, the NLRome and their evolutionary trajectories in Vitaceae remain poorly understood. We combined NLR-Annotator^87^ and Resistify^88^ to comprehensively annotate 454 to 862 NLR genes across 29 Viticeae genomes (**Fig. 6a and Supplementary Table 8**), with *V. vinifera* harboring the fewest genes but relatively more abundant than many eudicot plants^33^. Interestingly, despite the relatively large genome size (∼1 Gb), *C. rotundifolia* had an ordinary number of NLR genes (555), suggesting that NLR gene expansion is independent of genome expansion in Vitaceae (**Supplementary Table 8**). We constructed a phylogenetic tree using NB-ARC domain sequences across the Vitaceae family (**Supplementary Fig. 27a**). The results revealed lineage-specific differences: *A. glandulosa* contains a relatively lower proportion of CNLs but a higher proportion of TNLs compared with other species, whereas *P. tricuspidata* harbors a comparatively lower proportion of TNLs (**Supplementary Fig. 27b**). These patterns further underscore the rapid and dynamic evolution of NLR genes within Vitaceae.

**Fig. 6.**
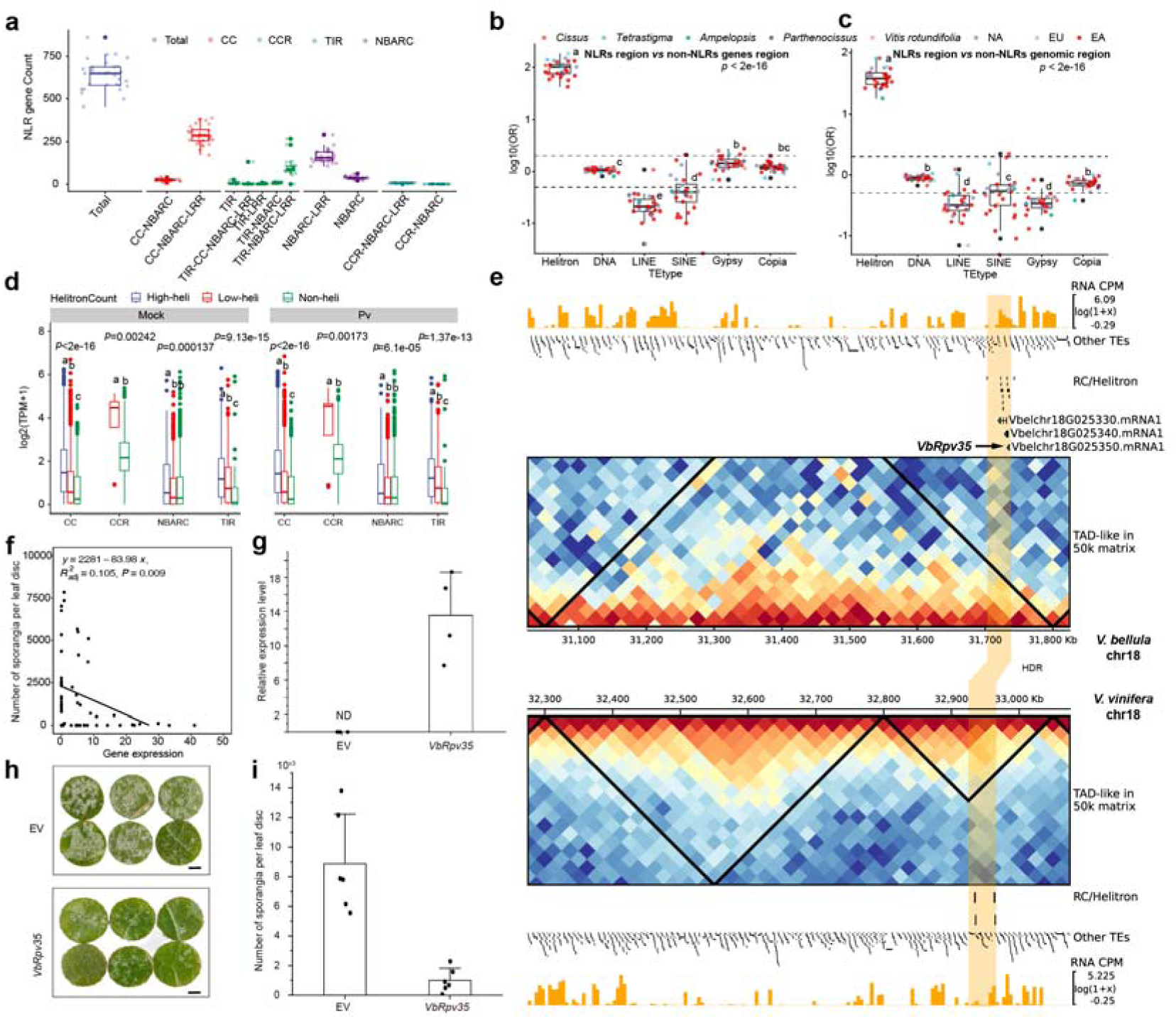
Helitron-mediated evolution and regulation of grapevine NLR resistance genes. **a**, Distributions of NLR gene types across 27 Vitaceae accessions. (TIR: Contains a Toll/interleukin-1 receptor (TIR) domain, CC: Contains a Coiled-coil (CC) domain without TIR domain, CCR: Contains an RPW8 (Resistance to Powdery Mildew 8)-like CC domain, NBARC: Lacks an N-terminal domain). **b-c.** Enrichment of different TE types in the 2-kb flanking regions of NLR genes, compared with non-NLR gene regions (**b**) and other genomic regions (**c**). Statistical significance was evaluated using odds ratio (OR) analyses. **d**, The impact of Helitrons frequency within the 2 kb upstream and downstream regions of NLR genes on the expression levels of different NLRs in 25*Vitis* samples (24 h post-infection (Pv) and Mock). The boxplot is generated using the expression levels of a single NLR gene from 25 different species. (n: CC=2166/10970/1636, CCR=NA/16/292, NBARC=514/4738/4030, TIR=2698/1806/114, High-heli/Low-heli/Non-heli). **e,** Comparison of NLR gene landscapes between downy mildew-resistant (*V. bellula*) and downy mildew-susceptible (*V. vinifera*) species. HDR: Highly Diverged Region. The triangle formed by two black lines and the baseline represents TAD-like regions. **f,** The correlation between the expression levels of *VbRpv35* homologs and sporangia count measured 24 hours after inoculation with *P. viticola*. **g,** The transcriptional level of the *VbRpv35* in grape leaves following transient transformation was verified using the empty vector (EV) as a control. The PCR validation was repeated three times independently with similar results. ND, not detected. **h and i,** Phenotypic analysis of grape leaf discs transiently transformed with either the empty vector (EV) or *VbRpv35* construct. Following a 2-day expression, the transformed leaf discs were inoculated with a *P.viticola* suspension (1 × 10^5^ sporangia/mL). Symptom imaging (**h**) and sporangia quantification (**i**) were conducted 7 d post-inoculation (dpi). Error bars represent standard error (SE) of the mean from three independent biological replicates.

Distribution of *Vitis* NLR genes is previously found biased towards specific chromosomes, with chr18 and chr19 significantly enriched for NLR genes^30^, similarly reported in *O. sativa* and *Leersia* ^89^. We thus traced how NLR genes may have evolved in the Vitaceae family preceding the emergence of chromosomal preferences. Based on synteny analysis using ACEK ancestral genome as a reference, we mapped the chromosomes of *V. vinifera, P. tricuspidata, A. glandulosa, T. hemsleyanum*, and *C. rotundifolia* against ACEK to retrieve their ancestral chromosomal segments (**Fig. 3, Supplementary Table 10**), and the number of NLR genes per segment (**Supplementary Fig. 28a**). Across all six Vitaceae species, NLRs were consistently enriched on only one chromosome per trio of ACEK-derived homoeologous chromosomes, with the same ACEK chromosome showing enrichment in all species (**Supplementary Fig. 28a**). Given that extensive chromosomal rearrangements occurred during the transition from ACEK to the derived AVK karyotype, this conserved pattern across independently rearranged genomes supported the inference that NLR enrichment preferences were already established in ACEK prior to AVK formation. Surprisingly, our analysis of NLR genes in *Citrullus lanatus, Solanum lycopersicum* and *Malus fusca* failed to show the similar NLR distribution pattern for these eudicots (**Supplementary Fig. 28b**).

### Helitrons are the putative major drivers of NLR evolution in Vitaceae

TEs are widely recognized for driving genome expansion and gene duplications^90^. We wonder whether any specific types of TEs orchestrated NLR evolution, and thus analyzed the distribution of 6 major TE types, including retrotransposons (*Copia*, *Gypsy*, LINE, and SINE) and DNA transposons (DNA transposons and Helitrons) near NLR and non-NLR genes. No enrichment of TEs was observed near genes, except for Helitrons, which were highly enriched in NLR genes but not for non-NLR genes across the Vitaceae (**Fig. 6 and Supplementary Fig. 29-33**). Strikingly, approximately 40% of the Helitrons were found near NLR gene regions (within 2 kb upstream and downstream regions), with 75% of NLR regions flanked by the Helitrons (**Supplementary Fig. 34**), showing a close association between NLR genes and the Helitrons. Given the ability of Helitrons to replicate adjacent sequences and insert them into new genomic locations via a rolling-circle mechanism^91^, Helitrons may have driven NLR gene expansion in the Viticeae. We observed that Helitrons elements can capture the coding sequences (CDS) of NLR genes in six representative species within Vitaceae (**Supplementary Fig. 35**). Moreover, statistical analyses demonstrated that Helitrons were significantly enriched within 2 kb upstream and downstream of NLR genes, whereas other types of transposable elements showed no such enrichment (**Fig. 6b-c**). These findings suggest that Helitrons transposition may represent a driving force underlying NLR expansion and diversification.

Helitrons are known to regulate gene expression in maize^91,92^, which prompted us to investigate their impact on grapevine NLR gene expression with and without infection. To this end, we performed RNA-seq of leaves from 25 *Vitis* species inoculated with and without *Plasmopara viticola* causing downy mildew at 0 and 24 h post-infection. We observed a positive correlation between Helitrons presence and NLR gene expression (**Fig. 6d**). This relationship was independent of *P. viticola* infection and NLR gene types (**Fig. 6d**), suggesting that Helitrons insertions may play a universal role in upregulating all types of NLR gene expression. Interestingly, we observed that other types of TEs had minor effects on only a few types of NLR gene expression (**Supplementary Fig. 36**). These results highlight the specific role of Helitrons in both NLR gene expansion and expression of Vitaceae plants.

Finally, through evolutionary genomic and multi-omic analysis, we identified an NLR gene from downy-mildew resistant grapevines but absent in susceptible ones, which could be deployed in improving grapevine disease resistance. The NLR gene *Vbelchr18G025350* (*VbRpv35*) from *V. bellula*, commonly known as beautiful grape, showed high expression and up-regulation upon *P. viticola* infection (**Supplementary Table 9**). This gene was located within an NLR gene cluster with proximity to a high density of Helitrons (**Fig. 6e**). Although a syntenic region was present in cultivated grape *V. vinifera*, it lacked the NLR genes and exhibited significantly fewer Helitrons (**Fig. 6e**). Genome alignment showed that this region was highly variable among different species with abundant TE insertions including Helitrons, where TAD boundary changes and the presence or absence of NLRs were observed (**Fig. 6e, Supplementary Fig. 38**). Interestingly, TAD boundary changed independent of other structural variations, likely due to the presence or absence of NLR genes (**Fig. 6e, Supplementary Fig. 38**). Furthermore, homologs of *VbRpv35* were identified in multiple *Vitis* species, including *V. rotundifolia* (*Vrotchr18G021550*). Strikingly, the expression levels of these homologs 24 hours post-infection across the *Vitis* accessions were negatively correlated with the number of *P. viticola* spores on leaf surfaces (**Fig. 6f**). Furthermore, transient expression of *VbRpv35* in leaves of *V. vinifera* significantly reduced disease symptom 7 d post-inoculation (**Fig. 6g-i**). These results together suggest that *VbRpv35* confers resistance against *P. viticola*.

## Discussion

The rise of grapevines (*Vitis*) and their divergence from relatives in the grape family has been intriguing evolutionary biologists. The exact time of their origin remains unknown, although there is paleobotanic evidence discovering 60 to 66 million-year-old fossils of grape seeds in Colombia and India^93^. Genomic technologies offer an exciting opportunity to dissect the evolutionary history of grapevines with genome sequences becoming available for *Vitis* species^8–14^, *Tetrastigma*^21^, and *C. rotundifolia*^22,23^. With the advancement of high-throughput sequencing technologies, we generated genome sequences from 27 Vitaceae species, including 26 new accessions and 7 new species in our study. These newly assembled genomes, especially the first chromosome-level reference genomes for two tribes, Ampelopsideae and Parthenocisseae, which correspond to newly sequenced species relative to previous studies^25^, provide valuable and unprecedented insights into the evolutionary relationships in Vitaceae. The phylogenomic tree using single-copy nuclear orthologs represents the first genome-based inference of evolutionary relationships among major Vitaceae clades, which showed that *A. glandulosa* diverged earlier than *P. tricuspidata* but later than *C. rotundifolia* and *T. hemsleyanum,* consistent with recent nuclear genome–based studies^25^. The functional differences among significantly expanded gene families also reflect species-specific evolutionary strategies for adaption to diverse environmental conditions^1,3,4^(**Supplementary Fig. 5 and 6**).

Previous Viticeae phylogenetic studies were mainly using specific nuclear or plastid genes^3,6,94–96^ with discrepancies between nuclear and plastid phylogenies. These studies have focused on several low-copy nuclear genes^4^ and transcript sequences^5,6^. While these approaches have provided valuable insights, the selective use of genes tends to introduce bias, potentially affecting the accuracy of phylogenetic inferences. Moreover, a systematic analysis of gene flow among species from the five tribes has not yet been conducted. We demonstrated that gene flow and incomplete lineage sorting (ILS) are critical factors likely contributing to such phylogenetic conflicts, as similarly shown in a butterfly evolution study^65^ (**Fig. 7a**). Additionally, Hi-C data analysis showed that both *A. glandulosa* and *P. tricuspidata* have *n* = 20 chromosomes (**Supplementary Fig. 3**), providing fresh chromosome-level genomic insights for these species and facilitating the investigation of karyotype evolution in Vitaceae. In Brassicaceae, shared chromosomal fusion events have been used to infer phylogenetic relationships^18^. Based on AVK reconstructed here, we identified a shared EEJ event (AVK/14_21/EEJ) among *A. glandulosa*, *P. tricuspidata*, and *Vitis* (**Fig. 3),** which is absent in *C. rotundifolia* and *T. hemsleyanum*. Together, these observations present strong evidence supporting a closer relationship between *A. glandulosa* and *P. tricuspidata* with *Vitis* than with the *C. rotundifolia* or *T. hemsleyanum*. The discovery and identification of Vitaceae fossils have shown that *Ampelopsis*, *Parthenocissus*, and *Vitis* have an oval chalaza, whereas *Cissus*, *Tetrastigma*, and *Cayratia* possess a long or linear chalaza^97^, consistent with our phylogenomic based taxonomy^93,97^. Therefore, compared to traditional gene-based phylogenies, this study, via integration of karyotype analysis, whole-genome comparative analysis, and phylogenomic based on whole-genome data, provides a more robust approach for inferring phylogenomic relationships in lineages with complex evolutionary histories.

**Fig. 7.**
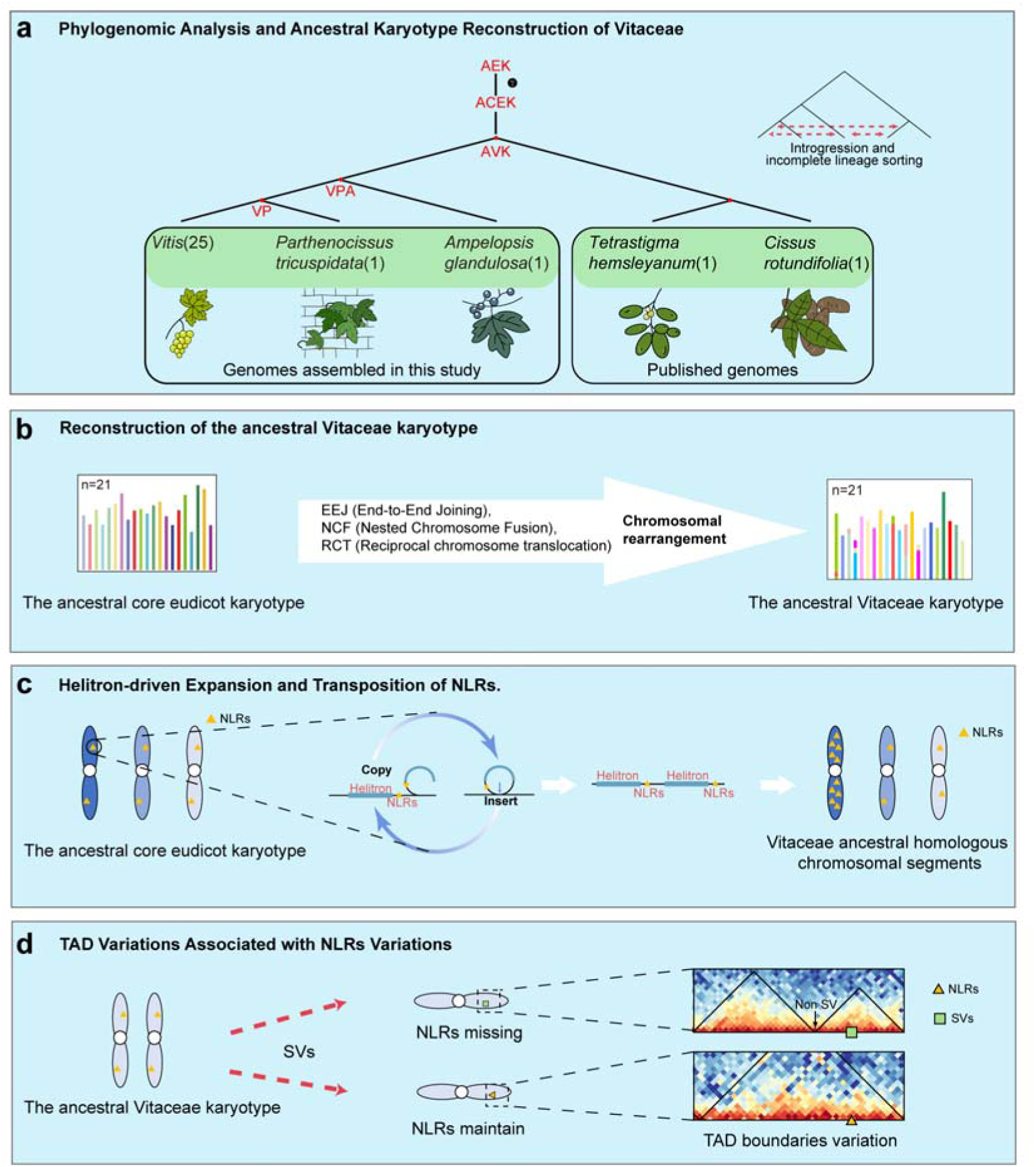
Schematic diagrams depicting the major genomic evolutionary events in Vitaceae. **a**, Phylogenomic analysis and ancestral karyotype reconstruction of Vitaceae. Phylogenomic analysis has revealed significant introgression and incomplete lineage sorting among species within the Vitaceae. γ represents the ancient whole-genome triplication (WGT) event shared by core eudicots. **b**, Reconstruction of the ancestral Vitaceae karyotype. **c**, Helitron-driven expansion and transposition of NLRs. Helitrons replication captures adjacent NLR genes and transposes them to new locations within the genome, exhibiting biased enrichment on ancestral homologous chromosomes. **d**, TAD variations associated with NLR variations. In the absence of structural variations (SVs) at TAD boundaries, intra-TAD structural variations can still contribute to TAD reorganization.

Karyotype variations drive species diversification and speciation. Reconstructing the evolution of Vitaceae karyotypes facilitates our understanding of the origin and evolution of grapevine genomes. So far, family-centric ancestral karyotype reconstructions are reported for only a few plant families^17,66–69^, but little is known about how grape family evolves at the karyotype perspective. The high-quality chromosome-level genome assemblies enabled our reconstruction of Vitaceae ancestral karyotypes (**Fig. 7b**), which provide insights into the rise and evolution of grapevines. In addition, we observed substantial karyotypic divergence between AVK and *T. hemsleyanum* (**Supplementary Fig. 18**). This divergence is not limited to RCTs, and involves extensive intrachromosomal SVs following rearrangement events. Notably, the formation of a single extant chromosome is often coupled with multiple successive rearrangements, with individual events incorporating segments derived from three or more AVK protochromosomes. These complex patterns suggest that additional intermediate karyotypes between AVK and *T. hemsleyanum* likely existed. Future high-quality genome assemblies from broader Vitaceae lineages will help fully resolve the karyotypic evolutionary trajectory from AVK to *T. hemsleyanum*.

Cultivated grapevines are more susceptible to diseases than their wild relatives due to continuous domestication and breeding. It remains uncharacterized how the karyotype evolution may have affected the repertoire of disease resistance genes in grapevines. Our pan-NLRome analysis of Vitaceae genomes revealed a biased distribution of NLR genes on one of the ACEK chromosomes (**Fig. 7c**). Several factors can influence gene distribution in plant genomes, such as natural selection and chromosomal rearrangements, which act genome-wide and unlikely favor certain chromosomes. The observed enrichment of Helitrons around NLRs thus prompted us to hypothesize that the action of Helitrons may contribute to the expansion of NLRs (**Fig. 6b-c**), particularly on one of three ACEK chromosomes. Although we observed Helitrons spanning NLR coding sequences, providing strong evidence for Helitron-mediated gene amplification, tracing their origins remains challenging because the rolling-circle replication mechanism of Helitrons does not leave deletions or duplications at the original locus^91^. Together with previous reports in Solanaceae, these observations suggest that Helitron-mediated capture and duplication of NLR genes may represent a conserved evolutionary mechanism in plants, a hypothesis that will require broader phylogenetic sampling to be rigorously tested.

Notably, Vitaceae is one of the few plant families without experiencing a unique WGD event^24^. Also lacking a unique WGD, watermelon had comparable NLR genes among homologous ancestral chromosomes (**Supplementary Fig. 28b**). Thus, we postulate that Helitrons activities influenced the distribution of NLR genes in eudicots following the shared γ event, which may have been adjusted by subsequent species-specific genome duplications in certain lineages, leading to the varied NLR gene distribution on ancestral chromosomes across different species. Although Helitrons have been implicated in the expansion of NLR genes, Helitrons copy numbers were comparable among ACEK-homologous chromosomal segments across the six species (**Supplementary Fig. 39**). This discordance suggests that Helitrons may have facilitated NLR expansion but were subsequently reduced through selective removal. The passive elimination of Helitrons may also affect nearby NLR retention^91^. For instance, for the region harboring *VbRpv35*, both *V. bellula* and *V. rotundifolia* possessed NLRs coupled with a high density of Helitrons (**Fig. 6e and Supplementary Fig. 38**), whereas in *V. vinifera*, all NLRs and the majority of Helitrons within this region have been lost, suggesting that the removal of Helitrons in *V. vinifera* may have concomitantly eliminated adjacent NLRs.

SVs regulate gene expression via disrupting cis-regulatory elements or even 3D genome organizations by alteration of TAD boundaries^84,85,98,99^. For example, LTR induced SVs at TAD boundaries in *Gossypium*^84,99^ and *G. max*^85^, leading to boundary changes. In this study, we performed the first genus-wide pan-3D genome evolution analysis for *Vitis*, observing a remarkable variation of spatial genome features such as TADs and A/B compartments, which are related to SV distributions. We observed that the number of SVs near TBVs was significantly lower than that in non-TBV regions. Further analysis revealed heterogeneous SV distributions across TBVs, with some variable boundaries exhibiting very few SVs and others showing relatively high SV density (**Supplementary Fig. 26**a), suggesting purifying selection that minimizes the detrimental effects of SVs on TAD boundary variations (**Fig. 5f**). It also indicates that *Vitis* TBVs are not necessarily driven by SVs at the boundaries. We propose that sequence variations within TADs, such as gene gains or losses, may also lead to changes in TAD structure. This is consistent with the fundamental role of TADs, where structural adjustments may occur to accommodate functional changes within the TAD (**Fig. 7d**). Chromosomal rearrangements in general disrupt 3D genomic organization and merge previously separated TADs which can facilitate novel enhancer-promoter interactions, thereby reshaping the local regulatory landscape^100,101^. However, our study showed that the chromosomal fusion in grapes did not account for the formation of a merged TAD. Rather, a distinct TAD boundary was maintained at the fusion site to effectively insulate the two flanking domains, although the insulation strength appeared slightly lower than that of the pre-fusion state (**Fig. 5f, g**). Notably, the insulation intensity at the end of *V. rotundifolia* Chr07 was substantially weaker than that the start of Chr20, hinting a higher degree of structural plasticity in this genomic region. We hypothesize that such disparities in subtelomeric insulation may be attributed to factors such as telomere attrition^102,103^, which could render this specific terminus more susceptible to inter-chromosomal interactions and subsequent fusion events, ultimately leading to the formation of Chr7. While useful for studying chromatin interaction landscapes, Hi-C data alone are insufficient to characterize the evolution of epigenetic regulatory signatures (*e.g.* enhancers) at finer scale. It necessitates a future investigation on the pan-cistrome of *Vitis* using technologies such as ATAC-seq and Hi-ChIP to understand the cis-regulatory element evolution in grapevine genomes behind trait evolution.

In summary, our genome assembly and analysis of 27 Vitaceae species provide fresh phylogenomic insight into the origin and evolution of grapevines and present valuable resources for future grapevine improvement and angiosperm evolutionary studies.

## Methods

### Preparation of plant materials

The Vitaceae accessions used in this study were grown under controlled conditions at two locations in China: the Zhengzhou Fruit Research Institute, Chinese Academy of Sciences (Zhengzhou, Henan, China), and the Institute of Advanced Agricultural Sciences, Peking University (Weifang, Shandong, China). All accessions were managed using standard viticultural techniques, which included routine tasks such as irrigation, fertilization, pruning, and disease prevention.

### Extraction of DNA and RNA

High molecular weight (HMW) genomic DNA was isolated from young grapevine leaves using the CTAB (cetyltrimethylammonium bromide) method. DNA quality was assessed using a Qubit® 2.0 Fluorometer (Life Technologies, CA, USA) and a pulse field gel electrophoresis apparatus (BioRad, CA, USA), following the manufacturer’s protocol. Total RNA was isolated from multiple tissues using Trizol RNA extraction reagent (Thermo Fisher). The extracted RNA was assessed using the RNA Nano 6000 Assay Kit of the Bioanalyzer 2100 system (Agilent Technologies, CA, USA). RNA samples with a RIN (RNA integrity number) value > 6.0 were processed for downstream library construction for RNA sequencing.

### Genome sequencing

To generate PacBio HiFi data, 15 µg of purified HMW DNA were used to construct a standard PacBio SMRTbell library (N50=15 kb) using PacBio SMRT Express Template Prep Kit 2.0 (Pacific Biosciences, Menlo Park, USA). Circular consensus sequencing was performed using a PacBio Sequel IIe instrument at Biomarker Technologies Corporation (Qingdao, China) and Berry Genomic (Beijing, China). For NGS paired-end sequencing, AMPure PB magnetic beads (Pacific Biosciences, Menlo Park, USA) were used to concentrate 15 µg of genomic DNA and applied for library construction according to the TruSeq DNA Sample Preparation Guide (Illumina, San Diego, USA). 150 bp paired-end (PE150) reads were generated using a HiSeq X Ten sequencing platform at Biomarker Technologies Corporation (Qingdao, China). For Hi-C sequencing, young leaves were cross-linked with 40 ml of 2% formaldehyde solution and applied for library construction following a standard Hi-C protocol. The Hi-C sequencing library underwent a test sequencing run to evaluate valid interaction read pairs using HiC-Pro (v3.1.0), followed by high-coverage sequencing with an Illumina NovaSeq 6000 platform.

### Transcriptome sequencing and analysis

To prepare an RNA sequencing library, a total amount of 1-3 μg RNA per sample was applied for library construction using VAHTS Universal V6 RNA-seq Library Prep Kit for Illumina® (Vazyme, Nanjing, China) following the manufacturer’s instructions. The RNA concentration of the library was initially quantified using Qubit® RNA Assay Kit on a Qubit® 3.0 (Thermo Fisher Scientific, Waltham, USA), diluting the RNA to 1 ng/µl. An Agilent Bioanalyzer 2100 system (Agilent Technologies, CA, USA) was employed to measure the insert size. After confirming that the insert size matches the expected size, a Bio-RAD CFX 96 quantitative PCR system (Bio-Rad, Hercules, USA) was used to precisely measure the effective concentration of the library (library effective concentration > 10 nm). A HiSeq X Ten sequencing platform at Biomarker Technologies Corporation (Qingdao, China) was used to execute the paired-end sequencing program (PE150) to produce 150 bp paired-end reads. Raw RNA-seq reads were further quality-filtered and trimmed using fastp^104^ (version 0.23.2) with default parameters. To conduct the quantification of gene expressions, reference indices were constructed using Kallisto (Bray et al., 2016) (parameters: index) with coding sequence (CDS) data from reference genomes. Transcript abundances were estimated using Kallisto^105^ with (parameters: quant), generating gene expression statistic in transcripts per million (TPM).

### Genome assembly

To assemble the T2T genome of *V. vinifera*, the HiFi (308×) and the ONT (117×) reads were employed to generate an initial assembly of two haplotypes using hifiasm^52^ (v0.19.8) (parameters:--hom-cov --ul). To remove plastid sequences from the initial assemblies, contigs were aligned to the *V. vinifera* mitochondrial and chloroplast genomes (GenBank accessions FM179380.1 and DQ424856.1, respectively) using minimap2^106^ (v2.24) (parameters: -x asm5). Contigs exhibiting at least 50% sequence identity with a plastid genome were removed from the final assemblies. Other contaminant sequences (e.g., microbial DNA) were further removed by aligning the contigs to all RefSeq bacterial genomes from NCBI using BLAST^107^ (v2.13.0+) (parameters: -task megablast). The filtered initial assemblies of two haplotypes were then scaffolded using the Hi-C reads by Juicer^54^ and 3d-DNA pipeline^55^. Hi-C interaction matrices were generated from the alignments using Juicer, and the assemblies were further scaffolded with 3D-DNA. Based on signals from Hi-C interaction heatmaps, manual adjustments were applied to the scaffolded pseudochromosomes using Juicebox^56^ (v1.11.08), generating chromosome-level assemblies. To close gaps on the chromosome-level assemblies, the ONT reads were first applied for filtering with Filtlong (v0.2.1) (https://github.com/rrwick/Filtlong) (parameters --min_length 80000 --min_mean_q 9), and consistency correction was performed for the filtered reads using NECAT^108^ (version 0.0.1). Next, the corrected ONT and HiFi reads were mapped to the chromosome-level assemblies, and sequences for gap filling were selected based on the following criteria: (1) the contig must span the entire gap region; (2) sequences flanking the gap (within 100 kb) must exhibit >90% sequence identity with the contig and have an alignment length exceeding 100 kb. Subsequently, the filled positions were examined in IGV^109^ for correctness validation. The genome assemblies of the other 26 Vitaceae accessions were generated from the HiFi reads using the aforementioned approaches without the gap-filling using ONT and HiFi reads.

### Quality assessment of assembled genomes

The generated genome assemblies were vigorously validated by following independent approaches: (1) LAI assessments were performed using LTR_FINDER_parallel^110^(v1.2 parameters: -harvest_out) and LTR_retriever ^59^ (v2.9.0 parameters: -inharvest) ; (2) QVs were assessed using Merqury ^58^ (v1.3 parameters: k = 21 counts); (3) BUSCO^57^.

### Centromere analysis

The processed CENH3 ChIP–seq data were obtained from the NCBI database under accession number PRJNA1021789. Reads were processed with fastp to remove adapter sequences and low-quality bases (Phred score < 20), then aligned to the respective genome assembly using Bowtie2 with default parameters, reporting up to ten valid alignments per read pair. Read pairs with MAPQ < 30 were discarded using samtools, and duplicate reads were removed with Picard MarkDuplicates (--REMOVE_DUPLICATES true --VALIDATION_STRINGENCY LENIENT). For reads mapping to multiple locations with varying alignment scores, the best alignment was selected, while alignments with ≥2 mismatches or with only one read in a pair were discarded. Coverage values were calculated as counts per million mapped reads using bamCoverage from deepTools^111^ v3.5.1. Broad peaks were called with MACS2^112^ v2.2.7.1, and reproducible peaks across replicates were identified using IDR^113^ v2.0.4.2.

### The annotation of Transposable Elements

To construct the de novo repeat library, we utilized RepeatModeler^73^. Repetitive elements were annotated by running RepeatMasker^74^ v.4.1.2, utilizing the Repbase database (v.20181026) for comprehensive annotation. Intact LTR (Long Terminal Repeat) elements were identified using a combination of LTR_Finder^110^ (v.1.2), LTRharvest^114^ (v.1.6.2), and LTR_retriever^115^ (v.2.9.0) pipelines. The LTR sequences from RepeatMasker and intact LTR elements obtained from LTR_retriever were subsequently processed through TEsorter^75^ (v.1.3) to classify the subfamilies of LTR retrotransposons (LTR-RTs).

### Genome annotation

To identify the genome assembly genes, we performed gene model prediction using an integrated pipeline incorporating the following data sources: ab initio predictions, homologous protein comparisons, and RNA-seq evidence. In the ab initio prediction phase, we trained the GeneMark-ET model using BRAKER2^116^ (v2.1.6) and further refined it by training the semi-HMM model SNAP via MAKER ^117^ (v3.01.03). The homologous protein sequences with non-redundant, human-curated were retrieved from the UniProt Swiss-Prot database (https://www.uniprot.org/downloads), ensuring their uniqueness. We further refined the homologous protein set using CD-HIT^118^ (v4.8.1) with default parameters to reduce potential redundancy. Transcriptome evidence was integrated into genome annotation, combining leaf transcriptome data with multi-tissue transcriptomic datasets from NCBI. Subsequently, each transcriptome dataset was utilized for gene annotation corresponding to the accessions of the respective population.

### Phylogenetic analysis

To construct the concatenated SCOs tree, we used the concatenated multiple sequence alignments of 2,365 SCOs generated by Orthofinder^119^ (parameters: -M msa). Poorly aligned regions were removed using TrimAl^120^ (v1.4.12) with parameters "-gt 0.6 -cons 60", yielding the trimmed supermatrix. Maximum likelihood phylogenetic trees were constructed using RAxML^121^ (v8.2.12) with the GAMMAJTT model. Divergence times were estimated using the Bayesian method implemented in the MCMCTREE program of PAML v4.9^122^. Calibration priors were defined based on previously published divergence time ranges obtained from TimeTree (www.timetree.org). Two uniform calibration constraints were applied: 33.2-59.1 Mya for the divergence between *P. tricuspidata* and *C. rotundifolia*^123–129^, and 2.5-10.18 Mya for the split between *V. rotundifolia* and the North American clade^123,130,131^. CodeML^122^ was used to evaluate amino acid substitution models. CAFE5^62^ was used to infer gene gain and loss rates for each genome. Orthogroups generated by OrthoFinder were treated as distinct gene families and provided as input for the CAFE5 analysis. The identified genes were further subjected to Gene Ontology (GO) and Kyoto Encyclopedia of Genes and Genomes (KEGG) enrichment analyses, with a significance threshold of p < 0.05.

For the coalescent-based phylogenetic analysis, each single-copy ortholog (SCO) sequence set was aligned using MUSCLE ^132^(v5.3) and trimmed with TrimAl (v1.4.12, - automated1) to remove ambiguously aligned regions. Maximum-likelihood (ML) gene trees were inferred using IQ-TREE^133^(v2.2.3) with the best-fit substitution model selected by ModelFinder (-m MFP) and branch support assessed with 1,000 ultrafast bootstrap replicates (-bb 1000). Bootstrap values were extracted from the nodes, and gene families with a median bootstrap support greater than 80 were retained for downstream analysis. The filtered ML gene trees were subsequently analyzed with ASTRAL^63^ (v5.7.1) to construct the coalescent-based consensus species tree.

### Gene flow analysis

Using BWA^134^ (v 0.7.18), the NGS reads from *Vitis vinifera*, *A. glandulosa*, *P. tricuspidata*, *C. rotundifolia*, *T. hemsleyanum* and *Liquidambar formosana* were aligned to the *A. glandulosa* genome with default parameters. Then, samtools^135^ (v.1.9) was used to sort the aligned reads, and Picard (v.2.21.6) was used to remove redundant reads. Variant detection and filtering were performed using GATK^136^ (v.4.1.2.0), with the filtering parameters set to ’QDL<L2.0, QUALL< 30.0, SORL>L3.0, FSL>L60.0, MQL<L40.0, MQRankSumL<-12.5, ReadPosRankSumL<L-8.0’. Patterson’s *D* and the *f*_b_ admixture ratios for all potential combinations were calculated using Dtrios within Dsuite^137^ (v.0.5 r47), utilizing SNP from GATK. To differentiate gene flow from incomplete lineage sorting (ILS), we employed the tree-based method QuIBL (https://github.com/miriammiyagi/QuIBL). For the QuIBL analysis, we prepared the input files by aligning 2,365 single-copy orthologous genes in fasta format. The BIC test, with a strict cutoff of ΔBICL>L10, was then applied to distinguish between the ILS-only model and the introgressionL+LILS model. The statistical significance of Patterson’s *D* values was assessed using the block-jackknife procedure implemented in Dsuite, which generates associated *Z*-scores for each tested combination of species.

### Synteny analysis

Synteny analysis was performed using JCVI^138^ (v1.1.19) by identifying syntenic blocks through an all-against-all BLAST search, followed by chaining hits with a distance cutoff of 20 genes.

### Reconstruction of ancestral chromosome

To assess synteny and homologous gene relationships across the genomes, we employed the WGDI^67^ toolkit (v0.74). Intra- and inter-genomic synteny blocks were identified using the -d parameter to generate homologous gene dot plots, following established pipelines for Brassicaceae study^18^. We reconstructed ancestral karyotypes at internal nodes using a bottom-up approach based on the chromosomal structures of extant species. Ancestral Karyotypes at key internal nodes—V, VP, VPA, and AVK—were inferred (**Data Availability**). For instance, a comparison between Euvitis and *V. rotundifolia* (Vr) revealed high conservation, with the exception of chromosome 7(**Supplementary Fig. 11a**). By integrating outgroup comparisons with *P. tricuspidata* (Pt) and *A. glandulosa* (Ag), we observed that Vr7 maintains synteny with Pt7 and Ag7(**Supplementary Fig. 11b**). This suggests that the karyotypic configurations of Vr7 and Vr20 represent an ancestral state predating the divergence of Euvitis. Consequently, the Vr genome was utilized as the representative model for the ancestral *Vitis* karyotype (V).

### Karyotypic Evolution analysis

We analyzed the chromosomes of the sampled Vitaceae species in comparison to the 21 inferred AVK protochromosomes and mapped the distribution of these ancestral chromosomal segments across the current genomes using the WGDI toolkit with the "-km" parameter. To reconstruct the chromosomal evolution of the Vitaceae family, we compared the karyotypes of phylogenetically proximal lineages (e.g., AVK-VPA and V-Euvits). Based on the features of three primary rearrangement events (RCT, NCF and EEJ), we inferred the specific rearrangement history within the Vitaceae clades (**Supplementary Fig. 15-19**).

### Structural variant detection and analysis

To detect SVs, we utilized Syri^139^ (v1.6.3) to analyze the genome assembly alignments against the *V.vinifera* reference, generated using nucmer^140^ (v 4.0.1) with parameters "-ax asm5 -eqx". This approach allowed for the identification of SVs, including insertions, deletions, inversions, translocations, and HDR. The detected SVs located in intergenic, upstream, downstream, intronic, and exonic regions were then annotated using ANNOVAR^141^ for further analysis.

### Hi-C data processing

HiC-Pro^142^ suite v.3.1.0 was used to perform valid mapping of Hi-C reads. Briefly, the Hi-C reads from the Vitaceae accessions were mapped to both the assembled genomes and the common reference genomes using Bowtie2^143^. During mapping, reads were aligned end-to-end, with those spanning ligation junctions trimmed at the 3’ end and realigned. Aligned fragment mates were paired into a single paired-end BAM file, while invalid Hi-C reads (dangling-end, same-fragment, self-circled, and self-ligation) were excluded. After duplicate removal, valid pairs generated raw Hi-C matrices at 10 kb to 1 Mb resolutions, followed by normalization using the iterative correction and eigenvector decomposition (ICE) method^144^. For contact domain analysis, valid pairs were converted into .hic format via Juicer tool^54^ (parameters: pre command). For compartment signal calculation and TAD-like or loop identification, they were converted into .cool format via HiCExplorer^145^ v3.7. (parameters: hicConvertFormat). Finally, Hi-C contact maps were visualized in .h5 format.

### Characterization of pan-3D genome architecture

The E1 values from the eigenvector decomposition of Hi-C contact maps determined A/B compartment status. We used cooltools^146^ (v0.7.0) (parameters: cooltools call-compartments) to obtain E1, E2, and E3 values from a 1-Mb resolution Hi-C contact matrix. Since E2 and E3 can also reflect A/B compartments, we manually examined E1–E3 tracks against gene density and plaid patterns to finalize the E1 list. Eigenvalues were assigned as A or B based on correlation with gene expression (CPM). Compartments were classified as Switch A/B or Stable A/B, depending on upstream bin consistency. To transfer A/B compartments from the target to the reference genome, we used minimap2, transanno^147^ v0.4., and CrossMap^147^ v0.7.0. The weighted average balanced E1 defined A/B compartments. Reference genome bins were categorized as Conserved A/B (A/B in all samples), Dispensable A/B (A/B or absence (E1=0 or NA) in all samples, no variation allowed), and Variable A/B. TAD boundaries were identified using a 50 kb resolution ICE-normalized matrix (parameters: --fdr 0.05) in HiCExplorer ^145^, from which insulation scores were also derived. Transanno and CrossMap projected TAD boundaries from the target to the reference genome, classifying them as Core (present in all samples), Dispensable (present in multiple samples), or Unique (present in one sample). TAD boundary variation (TBV) was determined by comparison to the reference genome, with TBV_Ins present in the target but absent in the reference, and TBV_Del absent in the target. SV enrichment in TBV regions was assessed by defining SV-Enriched TBV, where the number of SVs in a TBV region (b) exceeded those upstream (a) or downstream (c), satisfying (b > a and b ≥ c) or (b > c and b ≥ a).

### Identification of NLR genes

To identify NLR genes from the genomes, we used NLR-Annotator^87^ (v2.1b) to detect genomic fragments containing putative nucleotide-binding domains and leucine-rich repeat sequences, integrating the results with gene annotations. Additionally, Resistify ^88^(v0.5.2) was employed to identify and classify NLR genes based on protein sequences. The results from both tools were then integrated to provide the final identification and classification of NLR genes.

### Identification of disease-resistance genes

To identify disease-resistance genes, we used a stepwise filtering process. First, we screened for NLR genes located within variable TAD regions in disease-resistant species. Then, transcriptome data were used to further filter candidates, retaining only those genes whose expression showed significant changes upon *P. viticola* infection. *VbRpv35* is one of them.

To examine the association between *VbRpv35* expression levels and plant phenotypes using population-scale data, we identified homologs of *VbRpv35* in other *Vitis* accessions. The identification of NLR homologous genes involved two main steps. First, the protein sequences of *V. vinifera* and *V. bellula* were aligned to other genomes using Miniprot^148^ to identify homologous genes. Second, the protein sequence of *VbRpv3* was compared against protein sequences from other genomes using blastp, with a homology threshold set at e^-20^, to verify its homologous relationships. We then analyzed the correlation between the number of sporangia after *P. viticola* inoculation and the expression levels of *VbRpv3* homologs, using phenotypic and transcriptomic data obtained from previously published studies^8^.

### Construction of expression vector and transient expression in grape leaf

A 3657 bp DNA fragment of *VbRpv35* was synthesized and cloned into the plant expression vector pCR-35S containing the 35S promoter. Subsequently, the recombinant expression vector and the empty vector were transformed into *Agrobacterium tumefaciens* strain GV3101, respectively. The transformed strains were inoculated onto an LB solid medium supplemented with 25 µg/mL rifampicin and 50 µg/mL kanamycin and cultured at 28°C for 48 h. Single colonies containing the target plasmid or the empty vector were picked and inoculated into 5 mL of LB liquid medium with the same antibiotics, followed by pre-culture at 28°C for 8 h. The bacterial suspension was then transferred to 200 mL of fresh LB medium at a 1:40 ratio and cultured overnight at 28°C with shaking. Bacterial cells were collected by centrifugation at 4,000 rpm for 10 min and resuspended in infiltration buffer (0.5 M MES, 1 M MgCl_2_, 150 mM acetosyringone) to an OD_600_ of approximately 0.6.

Leaf discs (8 mm in diameter) were prepared from *V. vinifera* cv. Chardonnay leaves were placed on moist filter paper for vacuum infiltration for 2 minutes. The leaf discs were first cultured in the dark at 28°C for 24 h, then transferred to a growth chamber set at 18 ± 2°C with relative humidity of 80 ± 10% in a growth chamber under a 12 h white light (60 μmol·m^-2^·s^-1^)/12 h dark regimefor an additional 24 h of acclimation.

To verify the transcriptional level of *VbRpv35*, cDNA was synthesized from 500 ng of total RNA using SPARKscript II RT Plus Kit (with gDNA Eraser, Sparkjade). Quantitative PCR was performedusing 2xSYBR Green qPCR Mix (With ROX) (Sparkjade) and CFX Opus 96 (Bio-rad). Primers used in PCR amplification are as follows:5’-GCTCATTCGAGATGGTGTTTGG-3’; reverse: 5’-TCCACCTTGGTATCTTGTCCTTATT-3’ for *VbRpv35*; and 5’-CTTGCATCC CTCAGCACCTT-3’ and 5’-TCCTGTGGACAATGGATGGA-3’ for *VvACT1* as a reference (**Supplementary Table 11**). The CDS sequence of VbRpv35 is provided in Supplementary Table 11.

### Downy mildew infection assay

A 10 mL suspension of *P. viticola* at a concentration of 1×10^5^ sporangia/mL was prepared and incubated at 19°C for 1 hour to facilitate the release of zoospores. Grape leaf discs transfected with either the empty vector or the target vector were floated on the surface of the spore suspension with the abaxial side facing downward. The samples were then transferred to a growth chamber and maintained at 18 ± 2°C with relative humidity of 80 ± 10% in a growth chamber under a 12 h white light (μmol·m^-2^·s^-1^)/12 h dark regime for 12 h. Phenotypic observations and sporangia quantification were performed 7 d post-inoculation.

## Acknowledgments

We thank the Bioinformatics Platform at Peking University Institute of Advanced Agricultural Sciences (PKU-IAAS) for providing high-performance computing resources. Both L.G. and W.Y. are supported by the Key R&D Program of Shandong Province, China (2024CXPT031 and ZR202211070163), Taishan Scholars Program of Shandong Province, and Weifang Key Laboratory of Grapevine Improvement and Utilization, China. L.G. is also supported by the Natural Science Foundation for Distinguished Young Scholars of Shandong Province, China (ZR2023JQ010).

## Author Contributions

L.G., W.Y. and N.A.-K. conceived and supervised the project. J.M., W.Z., L.G., H.S. and C.L. curated and prepared the grapevine samples. X.W., J.S. K.W. and J.W. conducted genome assembly and annotation. X.W. and X.Z. performed ancestral karyotype analysis. S.C. performed variant calling. X.Z. performed the phenotyping and molecular experiments. L.G., W.Y. and N.A.-K. interpreted results. X.W., J.W., J.S., and S.C. prepared the figures and tables, L.G., X.W., J.S., J.J., F.S.K. and W.Y., wrote the manuscript. All authors read and approve the manuscript.

## Supplementary Figures and Tables

**Supplementary Fig. 1 |** The evaluation of genome assembly quality. The genome size and contig N50 of 27 Vitaceae species. The colors represent Viticeae species with *Vitis* accessions were categorized by different geographical origins.

**Supplementary Fig. 2 |** Hi-C heatmap for 12 samples. a-l: *Vitis romanetii, Vitis heyneana, Vitis yunnanensis, Vitis wuhanensis, Vitis amurensis, Vitis chunganensis, Vitis betulifolia, Vitis sinocinerea, Vitis rotundifolia, Vitis adstricta, Vitis labrusca and Vitis canadicans x Vitis rupestris*

**Supplementary Fig. 3 |** Hi-C heatmap for 15 samples. a-o: *Vitis rupestris, Vitis riparia, Vitis berlandieri, Vitis champini, Vitis wilsonae, Vitis lanceolatifoilosa, Vitis flexuosa, Vitis davidii, Vitis piasezkii, Vitis hancockii, Vitis bellula, Vitis pseudoreticulata, Vitis vinifera, Parthenocissus tricuspidata and Ampelopsis glandulosa*

**Supplementary Fig. 4 |** Coverage Profiles of HiFi and ONT Reads Mapped to *Vitis vinifera*.

**Supplementary Fig. 5 | a.** Expansion and contraction of gene families. The bar chart shows the number of gene families that have undergone expansion or contraction in each species. Significantly enriched KEGG pathways for expanded and contracted gene families in *T. hemaleyanum* (**b**) and *C. rotundifolia* (**c**).

**Supplementary Fig. 6 |** Significantly enriched KEGG pathways for expanded (top) and contracted (bottom) gene families in (**a**)*P. tricuspidata,* (**b**) *A. glandulosa* and (**c**) *Vitis*.

**Supplementary Fig. 7 |** Coalescence analyses of 2,365 concatenated single-copy genes. Summary of the proportion of gene tree topologies using single-copy genes.

**Supplementary Fig. 8 |** Dot plot of sequence alignment between *P. tricuspidata* and *V. vinifera*.

**Supplementary Fig. 9 |** Dot plot of sequence alignment between *A. glandulosa* and *V. vinifera*.

**Supplementary Fig. 10 | Chromosomal Synteny Analysis**: Dotplots of *V. vinifera* (a), *V. rotundifolia* (b), *P. tricuspidata* (c)*, A. glandulosa* (d), *T. hemsleyanum* (e) and *C. rotundifolia* (f) compared to ACEK.

**Supplementary Fig. 11 |** Construction of ancestral Vitis karyotype(V). **a**. Genome-wide synteny comparison between *V. rotundifolia* (Vr) and *V. vinifer*a (Euvitis). Chromosome 7 of *V. vinifera* (Euvitis) was formed by the EEJ fusion (V/7_20/EEJ) of Vr7 and Vr20. **b**. Chromosome 7 is intact in *P. tricuspidata* and *A. glandulosa*, indicating that the fusion of Vr7 and Vr20 occurred after their divergence. **c**. The *Vitis* karyotype (V) is consistent with that of *V. rotundifolia*.

**Supplementary Fig. 12 |** Construction of the ancestral karyotype of *Vitis* and Parthenocissus (VP). **a**. Genome-wide synteny comparison between the *Vitis* karyotype (V) and *P. tricuspidata*. Chromosomal rearrangements were detected involving chromosomes 4 and 20. **b**. Based on the shared karyotype structure among *V. rotundifolia, A. glandulosa* (Ag), and *P. tricuspidata* (Pt), chromosomes 4 and 20 of V were inferred to have formed later. Accordingly, chromosomes Pt4 and Pt20 were inferred as the ancestral VP chromosomes. **c**. Internal structural variations were also observed on chromosomes 6, 10, and 12 between V and pa. Pa10 and Pa12 are consistent with the karyotype of *V. rotundifolia* and were therefore inferred as ancestral VP chromosomes. d. Ancestral chromosome status of Vr6 was further validated using ACEK analysis.e. Origin and chromosomal composition of each inferred VP ancestral chromosome.

**Supplementary Fig. 13 |** Construction of ancestral VP and *A. glandulosa* karyotype (VPA).

**a.** Genome-wide synteny comparison between VP and *A. glandulosa*. Structural variations were detected on chromosomes 2, 7, 13, and 18 between them. **b.** Homologous chromosomes corresponding to VP chromosomes 2, 7, 13, and 18 were extracted from *A. glandulosa* and *C. rotundifolia*. Arrowheads indicate breakpoint positions between VP and *A. glandulosa*, whereas the chromosomal structures in *Cissus* are consistent with those of VP. **c**. Origin and chromosomal composition of each inferred VPA ancestral chromosome.

**Supplementary Fig. 14 |** Construction of ancestral Vitaceae karyotype (AVK). Genome-wide synteny comparison among ACEK(**a**), *T. hemsleyanum* (**b**), *C. rotundifolia* (**c**), and the VPA ancestral karyotype. Except for chromosome 14, all VPA chromosomes have karyotypically consistent counterparts in one of the other three genomes. **d**. Synteny comparison of chromosome 14 in the VPA karyotype with the homologous chromosomes from ACEK, *T. hemsleyanum*, and *C. rotundifolia*. *T. hemsleyanum* and ACEK share identical breakpoint positions, indicating that chromosome 14 of VPA was formed by an RCT fusion of two ancestral AVK chromosomes. AVK21 corresponds to Th21, whereas AVK14 is consistent with the distal segment of VPA chromosome 14. **e.** Origin and chromosomal composition of each inferred AVK ancestral chromosome. Vr14* represents genes Vr14_1 to Vr14_1011.

**Supplementary Fig. 15 |** Reconstruction of chromosome evolution in the ancestral AVK karyotype. a. Genome-wide synteny comparison between ACEK and the AVK. Red, green, and blue arrows indicate representative RTC-mediated rearrangements involving ACEK chromosome pairs (ACEK2/5, ACEK9/17, and ACEK3/21), which contributed to the formation of AVK chromosomes. b. Schematic illustration of the origin and chromosomal composition of AVK chromosomes inferred from synteny and breakpoint analyses.

**Supplementary Fig. 16 |** Evolutionary reconstruction of the transition from AVK to VPA. **a.** Genome-wide synteny comparison between the VPA and the inferred AVK. Strong collinearity is observed across most chromosomes. A clear end-to-end joining (EEJ) signal is detected between AVK14 and AVK21.

**b.** Schematic illustration of the formation of the VPA chromosome 14. VPA14 was generated by an EEJ-mediated fusion of AVK14 and AVK21.

**Supplementary Fig. 17 |** Evolutionary reconstruction of the transition from AVK to *C. rotundifolia*. **a.** Genome-wide synteny comparison between *Cissus rotundifolia* and the inferred AVK ancestral karyotype. Conserved syntenic blocks and breakpoint patterns reveal multiple chromosomal rearrangement events, including NCF and EEJ. **b.** Schematic reconstruction of the chromosomal rearrangements underlying the transition from AVK to *C. rotundifolia*.

**Supplementary Fig. 18 |** Breakpoint analysis of chromosomal fusion events from AVK to *T. hemsleyanum*.

**Supplementary Fig. 19 |** Chromosome-level synteny plots between *A. glandulosa* and VPA (a), and between *P. tricuspidata* and VP (b), indicate highly conserved karyotypes without detectable chromosomal rearrangements.

**Supplementary Fig. 20 |** Inference of phylogenetic relationships based on shared chromosomal breakpoints involving *A. glandulosa*. a. A chromosomal fusion event detected exclusively in Euvitis b. A chromosomal fusion event shared by Euvitis and *V. rotundifolia*. c. A chromosomal fusion event conserved in four species, being present in all analyzed samples except *T. hemsleyanum* and *C. rotundifolia*, suggesting that this rearrangement occurred after the divergence of *T. hemsleyanum* and *C. rotundifolia*.

**Supplementary Fig. 21 |** Phenotypes of selected *Vitis* accessions and TE annotations: **a**. Proportion of various intact LTRs; **b**. Relationship between Alesia, Angela, Blanca, CRM, and genome size.

**Supplementary Fig. 22 |** Relationship between Reina, Ogre, SIRE, Tekay, Tork, TAR, Galadriel, Ivana, Ikeros and genome size.

**Supplementary Fig. 23 |** The landscape of structural variants. **a.** Size and frequency of various types of structural variations. **b**. The relationship between genes and structural variation distance. **c**. Frequency of structural variations occurring in each gene interval.

**Supplementary Fig. 24 |**KEGG and GO functional enrichment of genes in Variable A/B (**a**) and Broad-Conserve B (Conserve B and Dispensable B) (**b**).

**Supplementary Fig. 25 |** KEGG and GO functional enrichment of genes in Broad-Conserve B (Conserve B and Dispensable B).

**Supplementary Fig. 26 |** Structural variants associated with TAD boundary variation (TBV). **a**, Distribution of structural variants (SVs) at TBVs and non-TBVs, partitioned into the boundary itself, 50 kb upstream, and 50 kb downstream, as well as combined regions (TBV + upstream 50 kb; TBV + downstream 50 kb). **b**, Statistical comparison of SV abundance between TBVs and non-TBVs, assessed by two-tailed t-tests. Significance levels are indicated by *P* values. **c**, Comparative analysis of TAD boundary–associated SVs among grapevine populations, with enrichment significance evaluated by odds ratio (OR) tests.

**Supplementary Fig. 27 |** A phylogenetic tree using NB-ARC domain sequences across the Vitaceae family(**a**). The relative proportions of different NLR clades are shown for each species (**b**).

**Supplementary Fig. 28 |** The number of NLR genes on ancestral chromosomal segments. **a.** The distribution of NLRs on ancestral chromosomal segments in 6 Vitaceae species.NLR genes identified in each genome were assigned to ancestral chromosomal segments based on conserved synteny relationships. **b.** The distribution of NLRs on ancestral chromosomal segments in *Citrullus lanatus, Solanum lycopersicum* and *Malus fusca,* shown for comparison with Vitaceae species. Ancestral chromosomal segments are labeled according to their origin from the ancestral eudicot karyotype (AEK), including subchromosomal segments generated by subsequent chromosomal rearrangements. Bar heights indicate the number of NLR genes per ancestral segment. AEK labels with numerical suffixes (for example, AEK1-2-3) denote distinct subchromosomal segments derived from the same ancestral core eudicot chromosome (ACEK1) following the core eudicot-whole-genome triplication (hexaploidization, γ event).

**Supplementary Fig. 29 |** The TE frequency within the 2 kb upstream and downstream regions of NLR and non-NLR genes in *Vitis romanetii, Vitis heyneana, Vitis yunnanensis, Vitis wuhanensis, Vitis amurensis* and *Vitis chunganensis*.

**Supplementary Fig. 30 |** The TE frequency within the 2 kb upstream and downstream regions of NLR and non-NLR genes in *Vitis betulifolia, Vitis sinocinerea, Vitis rotundifolia, Vitis berlandieri, Vitis champini,* and *Vitis wilsonae*.

**Supplementary Fig. 31 |** The TE frequency within the 2 kb upstream and downstream regions of NLR and non-NLR genes in *Vitis lanceolatifoilosa, Vitis flexuosa, Vitis davidii, Vitis piasezkii, Vitis hancockii,* and *Vitis bellula*.

**Supplementary Fig. 32 |** The TE frequency within the 2 kb upstream and downstream regions of NLR and non-NLR genes in *Vitis pseudoreticulata, Vitis adstricta, Vitis labrusca, Vitis canadicans x Vitis rupestris, Vitis rupestris and Vitis riparia*.

**Supplementary Fig. 33 |** The TE frequency within the 2 kb upstream and downstream regions of NLR and non-NLR genes in *C. rotundifolia*, *T. hemsleyanum*, *Vitis vinifera*,*P. tricuspidata*, and *A. glandulosa*.

**Supplementary Fig. 34 |** The positional relationship between NLR and Helitrons. NLR-Helitron: Helitrons surrounded by NLRs. NLR-Helitron family: Types of Helitrons surrounded by NLRs. Helitron-NLR: NLRs surrounded by Helitrons. Truncated-Helitron-NLR: NLRs surrounded by Helitron fragments. None-Helitron-NLR: NLRs with no identified Helitrons or Helitron fragments nearby.

**Supplementary Fig. 35 |** Helitrons carrying NLR coding sequences in six representative Vitaceae species: *V. vinifera*(a), *V. rotundifolia*(b), *P. tricuspidata*(c), *A. glandulosa*(d), *C. rotundifolia*(e), and *T. hemsleyanum*(f).

**Supplementary Fig. 36 | a.** The impact of TE frequency within the 2 kb upstream and downstream regions of NLR genes on the expression levels of different NLRs in 27 *Vitis* samples (Mock). The boxplot is generated using the expression levels of a single NLR gene from 25 different species. (n (High/Low/Non): for DNA, CC=2908/2612/1832, CCR=108/36/9, NBARC=1725/1732/1160, TIR=1066/793/494; for Gypsy, CC=577/1407/5368, CCR=17/46/90, NBARC=423/1219/2975, TIR=254/582/1467; for Copia, CC=1077/2110/4165, CCR=41/56/56, NBARC=806/1430/2381, TIR=385/670/1248; for LINE, CC=176/557/6619, CCR=1/1/151, NBARC=117/438/4062, TIR=69/177/2057; for SINE, CC=5/93/7254, CCR=NA/1/152, NBARC=2/47/4568, TIR=2/59/2242.). **b.** The impact of DNA, Gypsy, Copia, LINE and SINE frequency within the 2 kb upstream and downstream regions of NLR genes on the expression levels of different NLRs in 25*Vitis* samples (*Pv* infection). The boxplot is generated using the expression levels of a single NLR gene from 25 different species. (n (High/Low/Non): for DNA, CC=2908/2612/1832, CCR=108/36/9, NBARC=1725/1732/1160, TIR=1066/793/494; for Gypsy, CC=577/1407/5368, CCR=17/46/90, NBARC=423/1219/2975, TIR=254/582/1467; for Copia, CC=1077/2110/4165, CCR=41/56/56, NBARC=806/1430/2381, TIR=385/670/1248; for LINE, CC=176/557/6619, CCR=1/1/151, NBARC=117/438/4062, TIR=69/177/2057; for SINE, CC=5/93/7254, CCR=NA/1/152, NBARC=2/47/4568, TIR=2/59/2242.)

**Supplementary Fig. 37 |** The correlation between the number of Helitrons and the number of other five types of TEs within the 2 kb upstream and downstream regions of NLR genes.

**Supplementary Fig. 38 |** The TAD landscape of the *Rpv-Vbel1* gene in the *V. rotundifolia* and *V. vinifera* genomes. The figure is divided into two parts: the upper part represents the *V. rotundifolia* genome information, and the lower part represents the *V. vinifera* genome information. The connecting bands between the two regions indicate high variability areas. In the TAD track, the triangular areas formed by the broken lines represent TAD-like regions.

**Supplementary Fig. 39 |** The distribution of Helitrons on ancestral chromosomal segments in 6 Vitaceae species. The bar plot shows the number of Helitrons identified on ancestral chromosomal segments (ACEK-derived segments) across six Vitaceae species. Each bar represents the Helitrons count associated with a specific ancestral chromosome segment, and colors indicate different species as shown in the legend.

**Supplementary Table 1.** Statistics summary of genome sequencing reads (PacBio HiFi, NGS, ONT and Hi-C) generated for the 27 Vitaceae accessions.

**Supplementary Table 2.** The genomic features of 27 Vitaceae accessions. The table summarizes genome assembly statistics and quality metrics for each accession, including assembly size, contiguity (number of contigs, largest contig, contig N50, and gaps), base-level accuracy (QV), gene space completeness assessed by BUSCO, long terminal repeat (LTR) assembly quality (LAI), anchor rate, proportion of repetitive elements, GC content, and genome-wide heterozygosity.

**Supplementary Table 3.** Summary of genome annotation and BUSCO assessment for Vitaceae species.

**Supplementary Table 4.** The results of Dsuite for Vitaceae. D-statistics (ABBA–BABA tests) were calculated for different species trios to assess signatures of introgression among Vitaceae lineages. For each test, the table reports the samples assigned as P1, P2, and P3, the D statistic, Z-score, p-value, and f4-ratio, as well as the counts of BBAA, ABBA, and BABA site patterns.

**Supplementary Table 5.** The results of QuIBL analysis for different triplet tree topologies. QuIBL was applied to evaluate support for incomplete lineage sorting (ILS) versus introgression among different species triplets. For each triplet, analyses were performed using each taxon in turn as the outgroup. Mixprop1 and Mixprop2 represent the estimated mixture proportions of the two models, while Lambda1dist and Lambda2dist denote the corresponding exponential rate parameters. Bic1dist and Bic2dist indicate Bayesian Information Criterion (BIC) scores for the single- and two-distribution models, respectively. Count represents the number of gene trees supporting each triplet configuration.

**Supplementary Table 6.** Detailed locations of fusion breakpoints for shared major chromosomal rearrangements across sample species.

**Supplementary Table 7.** Lengths of various TEs across 29 Vitaceae species

**Supplementary Table 8.** The number of different types of NLR genes in 27 species.

**Supplementary Table 9.** Expression Levels (TPM) of *VbRpv35* under Mock and *Pv* infection conditions.

**Supplementary Table 10**. Information on chromosomal breakpoints identified based on ACEK homology. Homologous chromosomal segments corresponding to the ancestral core eudicot karyotype (ACEK) were identified and used to partition chromosomes of extant species. For each breakpoint, the table reports its genomic coordinates and the corresponding ACEK-derived chromosomal segments, providing a framework for reconstructing chromosomal rearrangement events across lineages.

**Supplementary Table 11.** The oligo and CDS sequences of *VbRpv35* and the oligo sequences of *VvACT1*.

