## Supplementary_figure for "Origin and evolution of grapevine genomes"

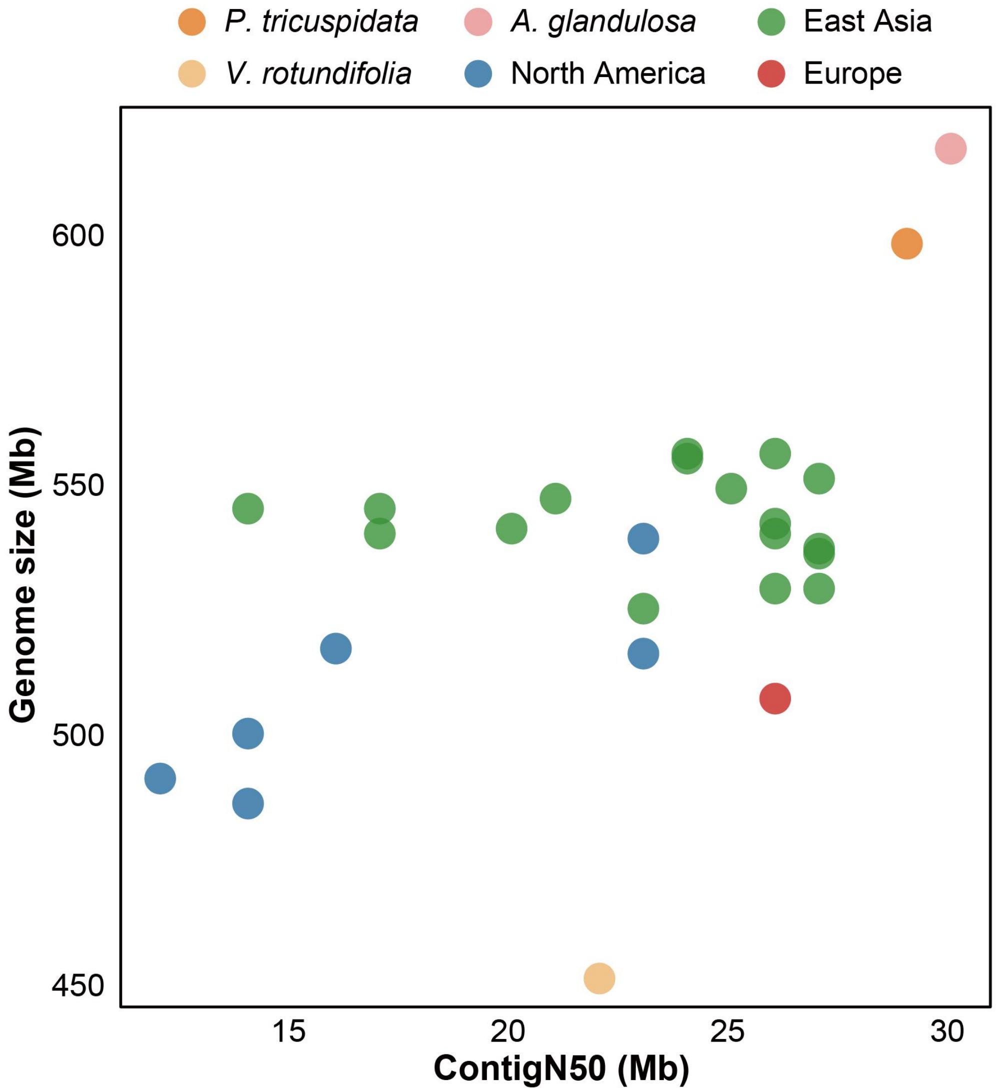


**Supplementary Fig. 1 |** The evaluation of genome assembly quality. The genome size and contig N50 of 27 Vitaceae species. The colors represent Viticeae species with *Vitis* accessions were categorized by different geographical origins.

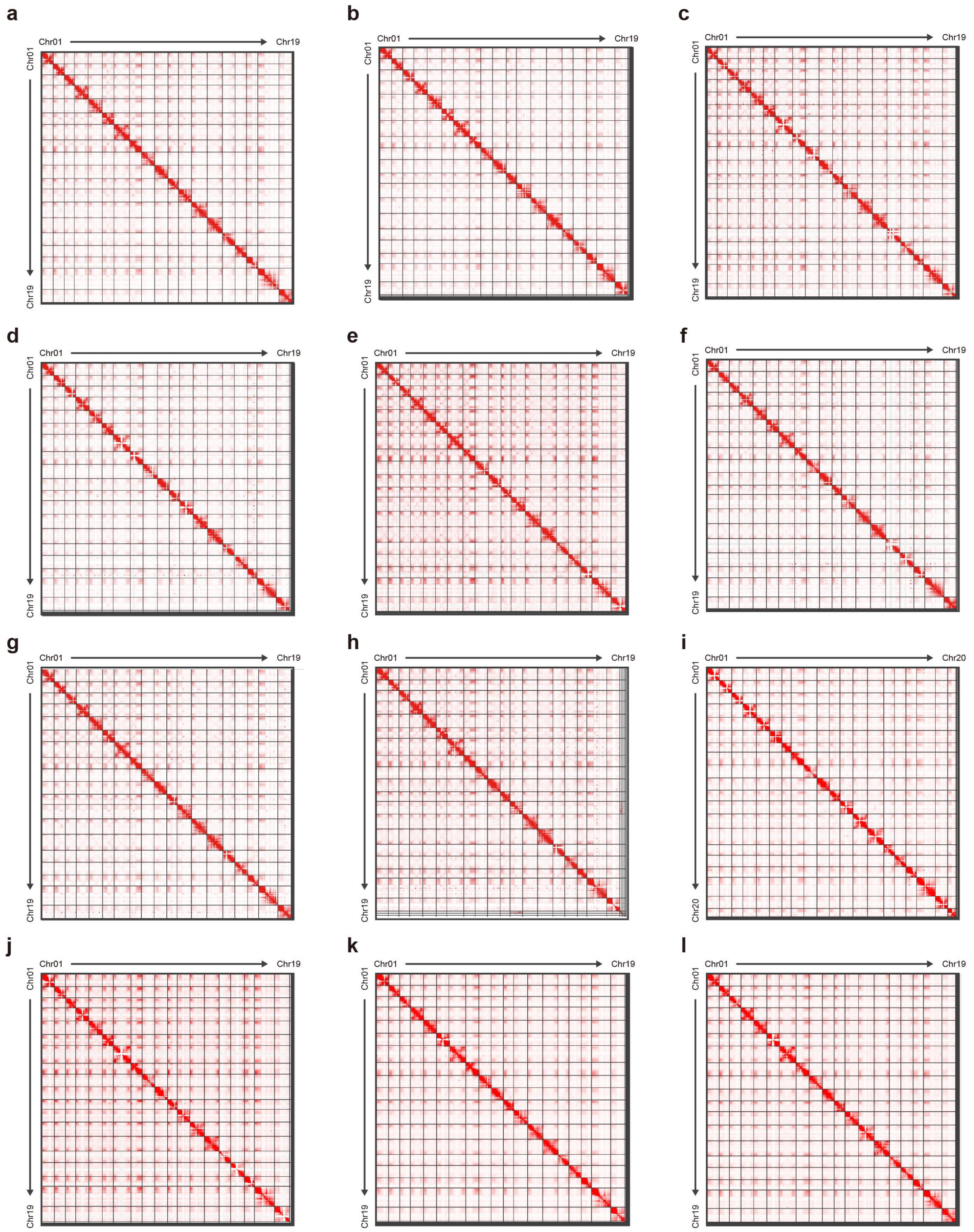


**Supplementary Fig. 2 |** Hi-C heatmap for 12 samples. a-l: *Vitis romanetii, Vitis heyneana, Vitis yunnanensis, Vitis wuhanensis, Vitis amurensis, Vitis chunganensis, Vitis betulifolia, Vitis sinocinerea , Vitis rotundifolia, Vitis adstricta, Vitis labrusca and Vitis canadicans x Vitis rupestris*

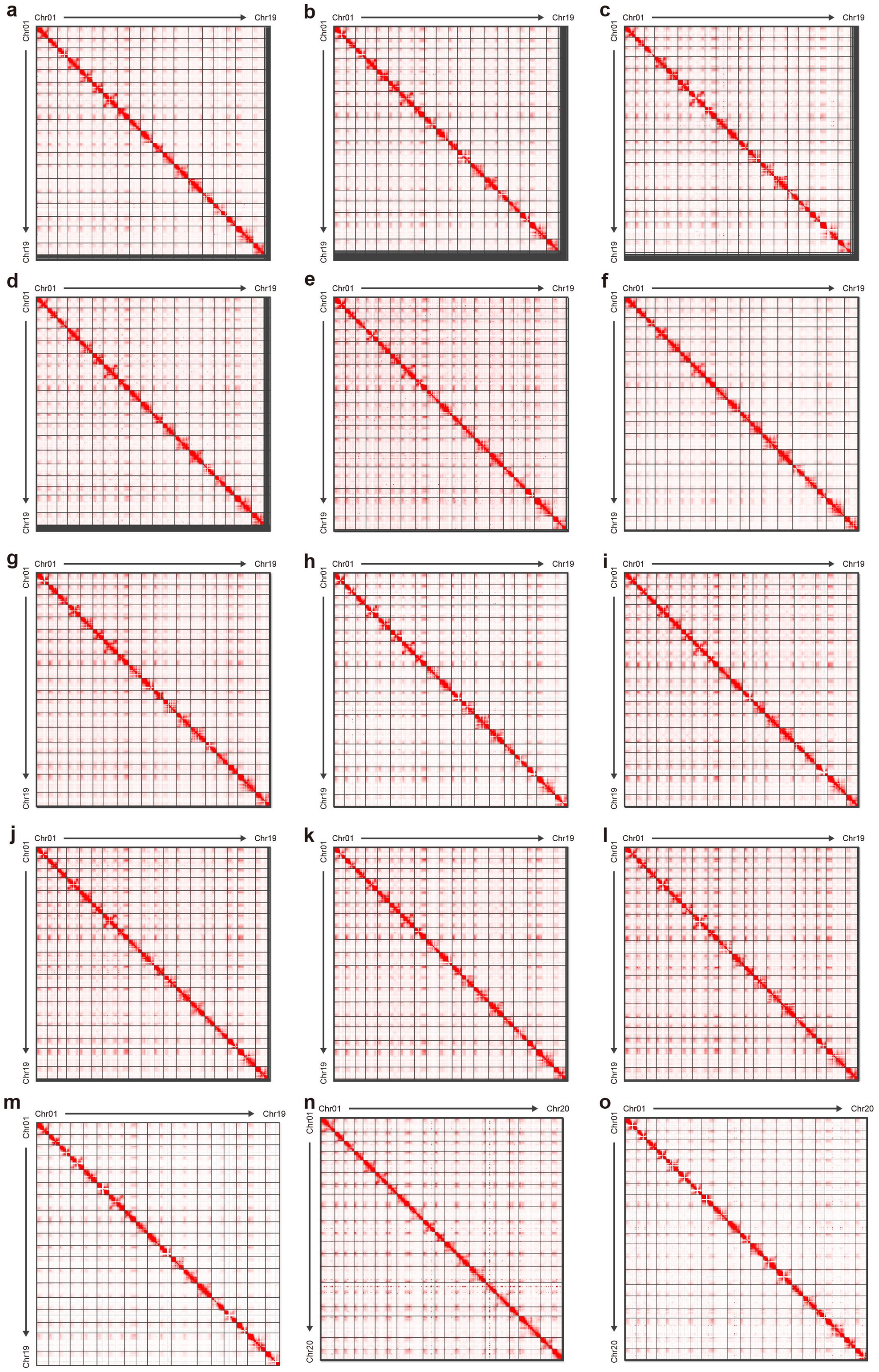


**Supplementary Fig. 3 |** Hi-C heatmap for 15 samples. a-o: *Vitis rupestris, Vitis riparia , Vitis berlandieri, Vitis champini , Vitis wilsonae, Vitis lanceolatifoilosa, Vitis flexuosa, Vitis davidii , Vitis piasezkii, Vitis hancockii, Vitis bellula, Vitis pseudoreticulata, Vitis vinifera, Parthenocissus tricuspidata and Ampelopsis glandulosa*

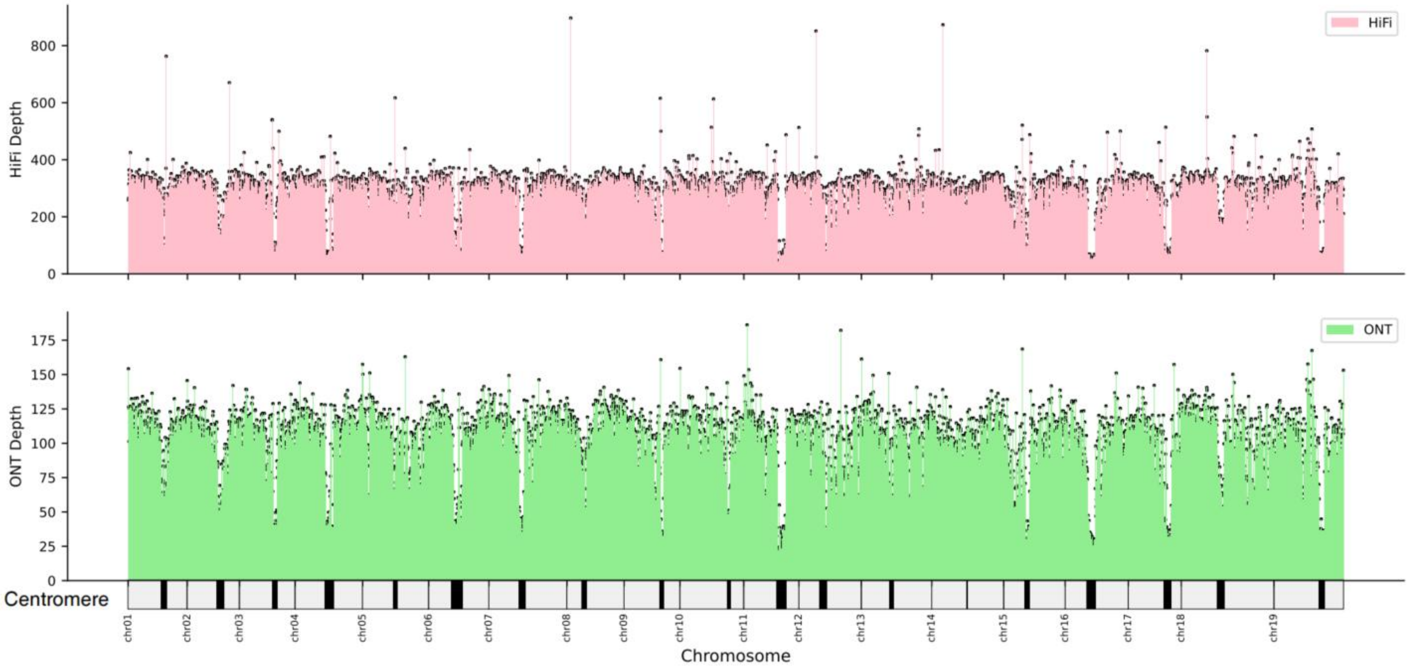


**Supplementary Fig. 4 |** Coverage Profiles of HiFi and ONT Reads Mapped to *Vitis vinifera*.

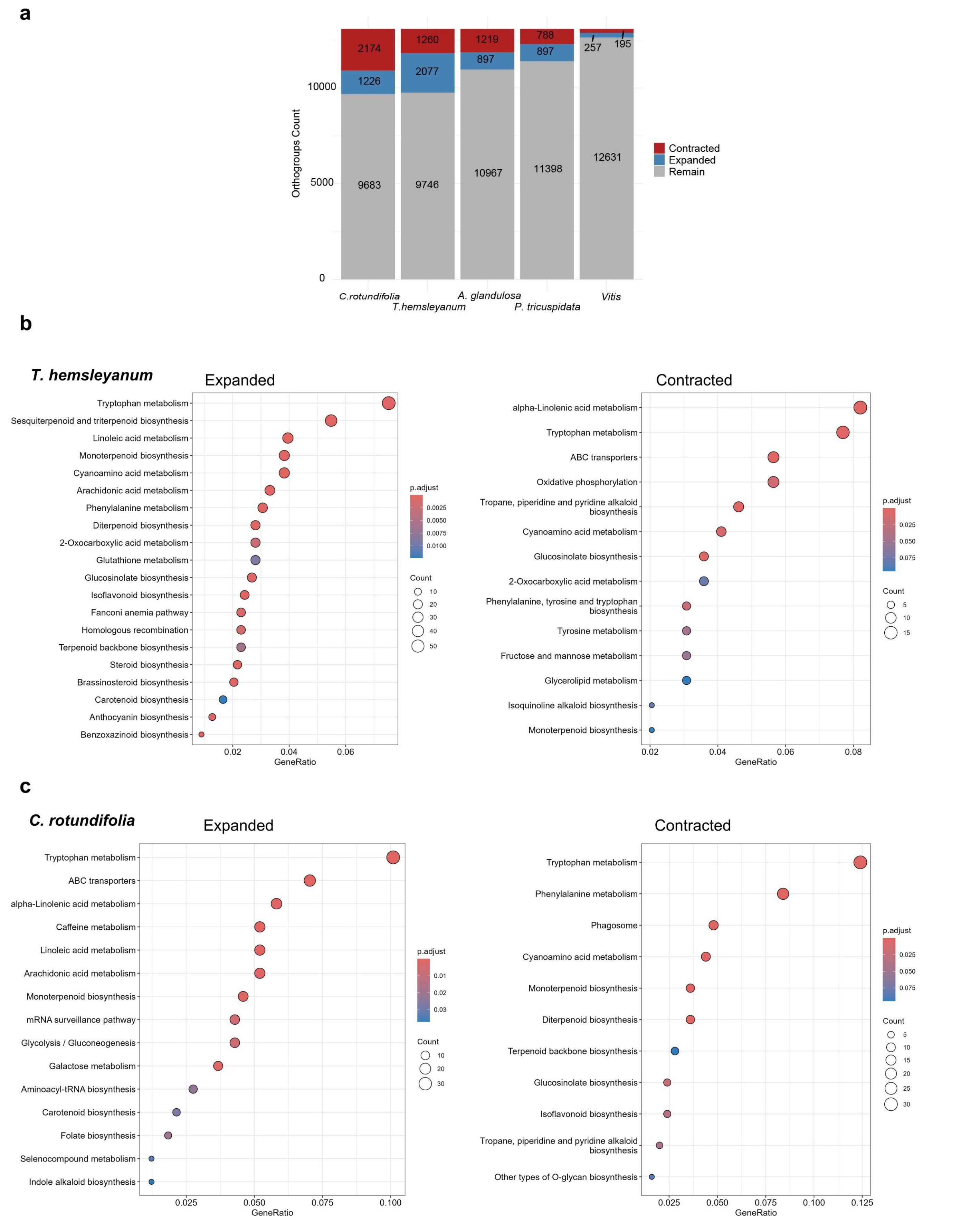


**Supplementary Fig. 5 | a.** Expansion and contraction of gene families. The bar chart shows the number of gene families that have undergone expansion or contraction in each species. Significantly enriched KEGG pathways for expanded and contracted gene families in *T. hemaleyanum* (**b**) and *C. rotundifolia* (**c**) .

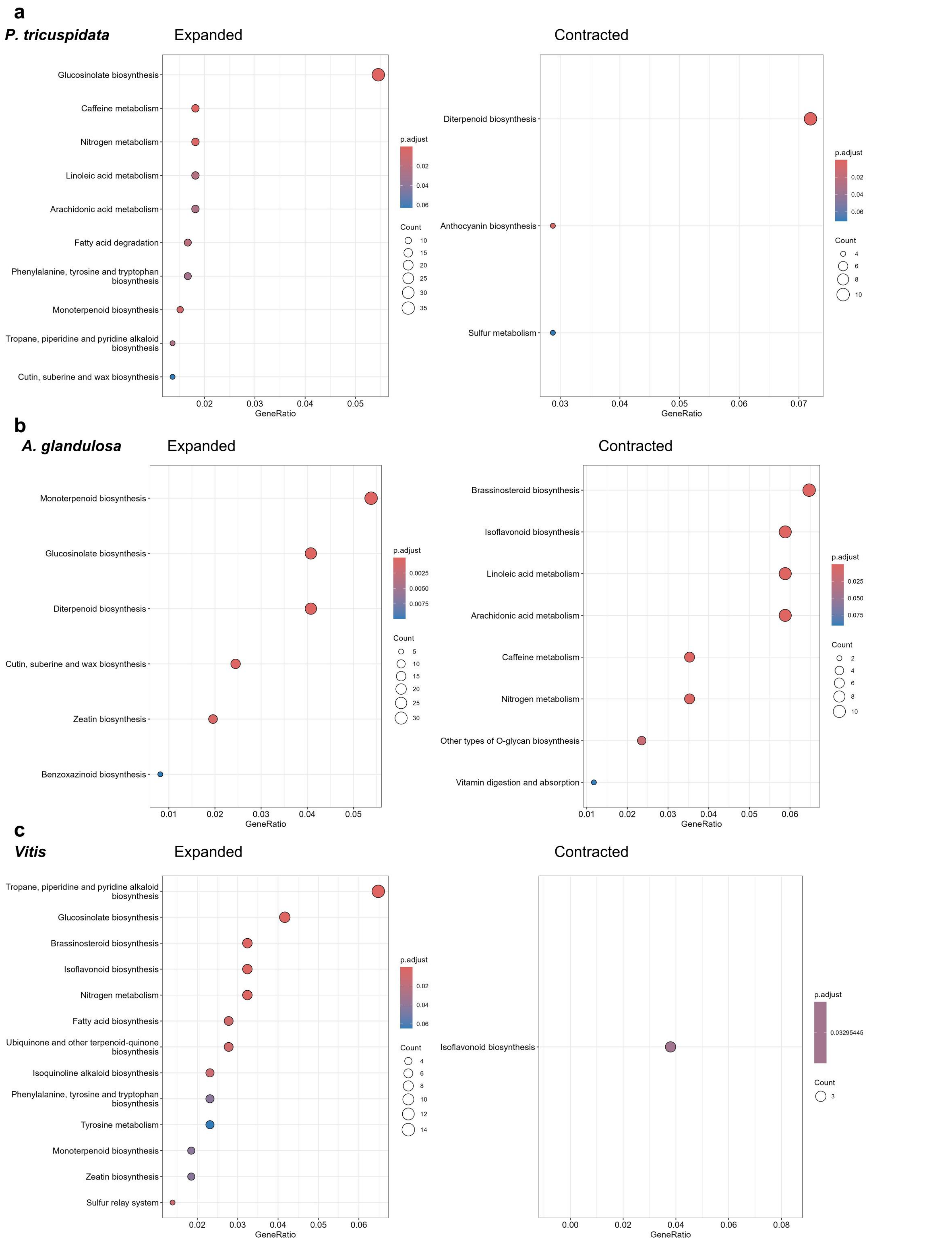


**Supplementary Fig. 6** | Significantly enriched KEGG pathways for expanded (top) and contracted (bottom) gene families in (**a**)*P. tricuspidata,* (**b**) *A. glandulosa* and (**c**) *Vitis*.

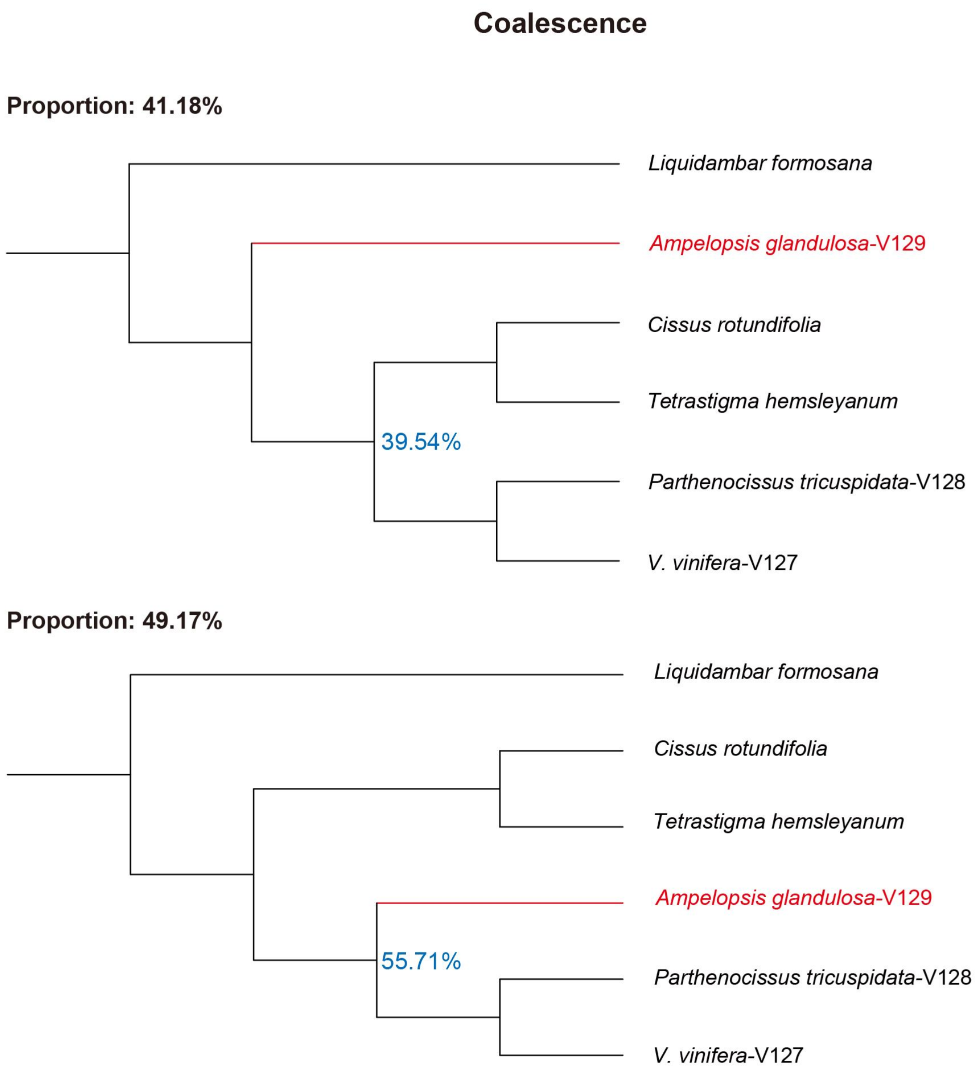


**Supplementary Fig. 7 |** Coalescence analyses of 2,365 concatenated single-copy genes. Summary of the proportion of gene tree topologies using single-copy genes.

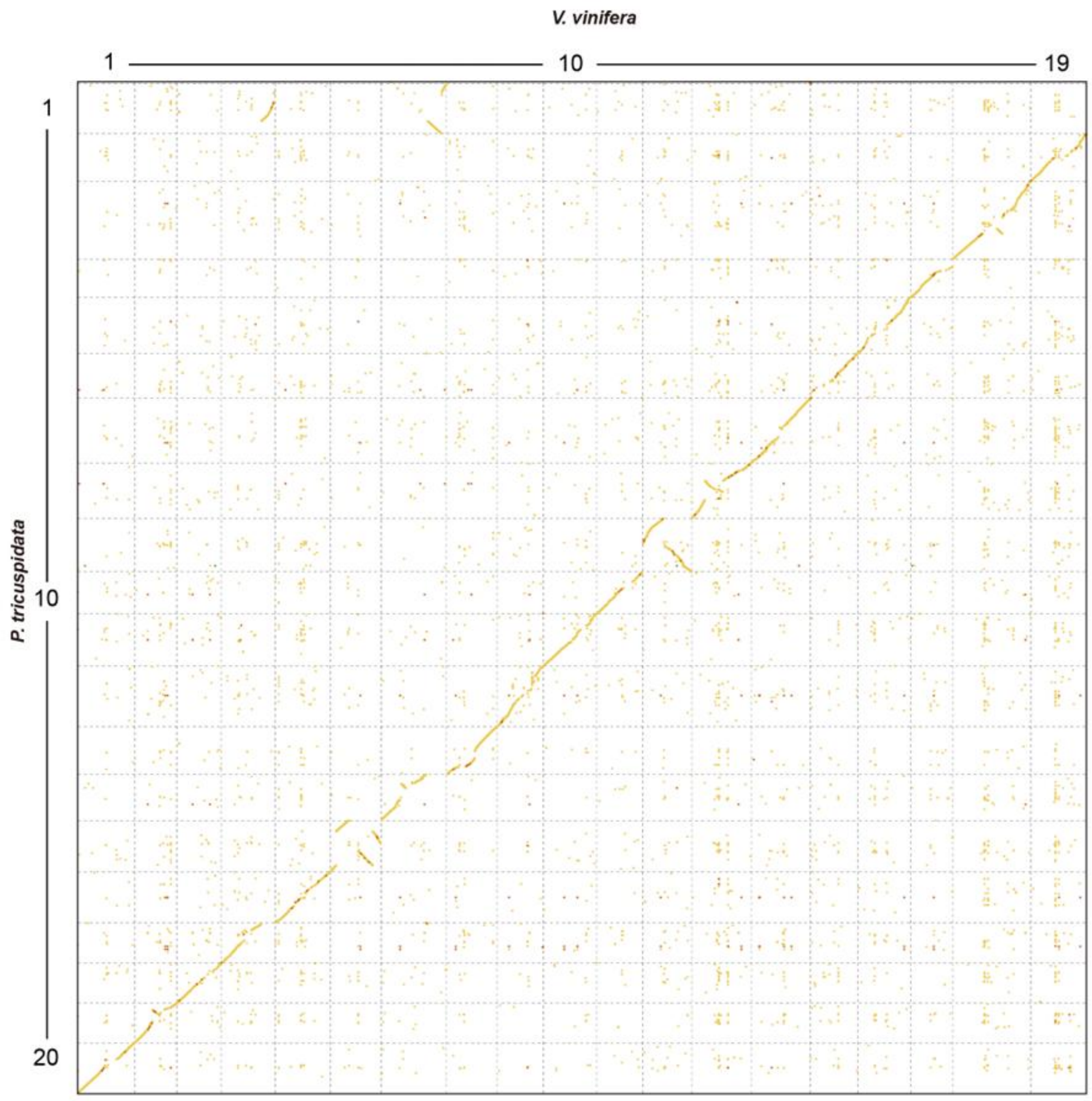


**Supplementary Fig. 8 |** Dot plot of sequence alignment between *P. tricuspidata* and *V. vinifera*.

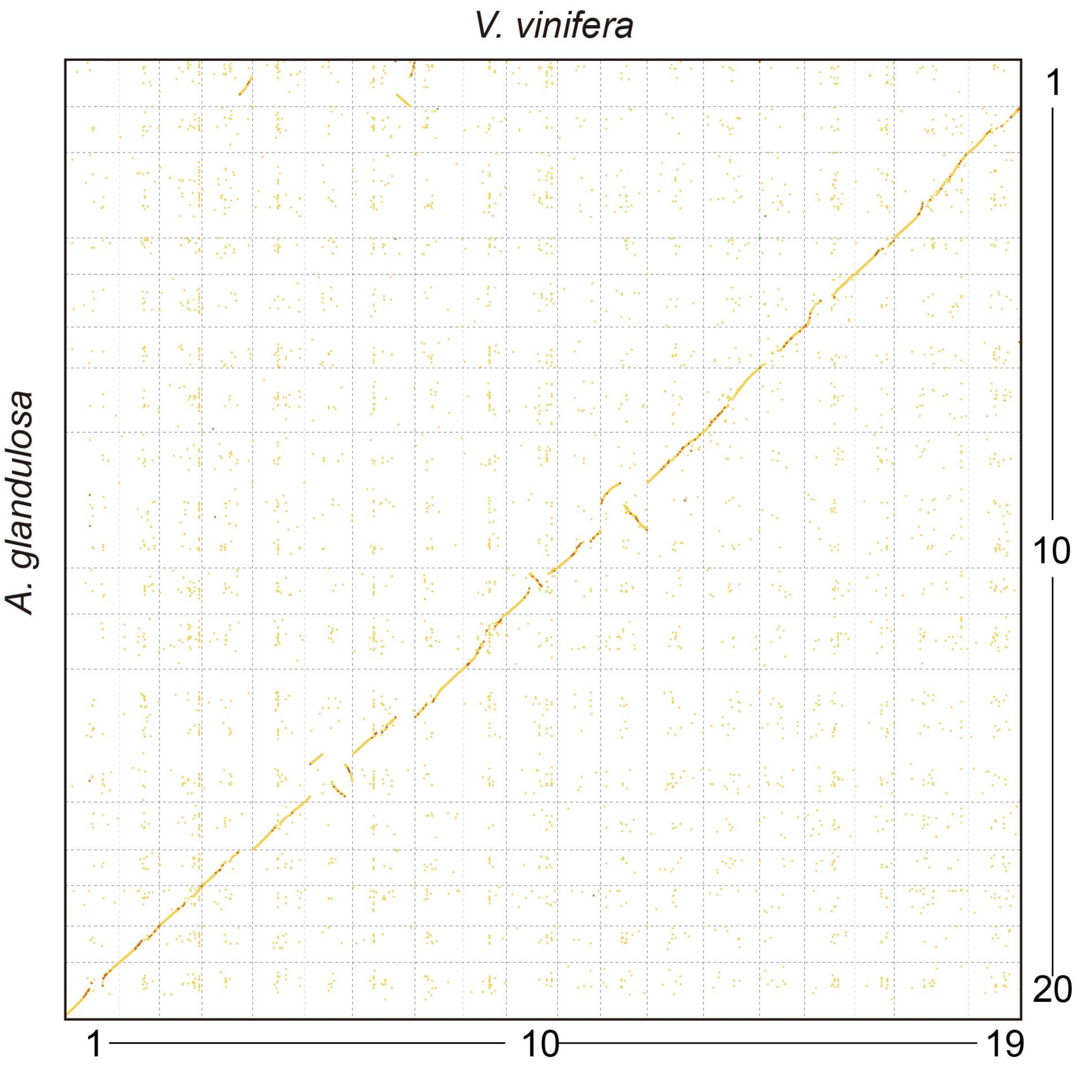


**Supplementary Fig.9 |** Dot plot of sequence alignment between *A. glandulosa* and *V. vinifera*.

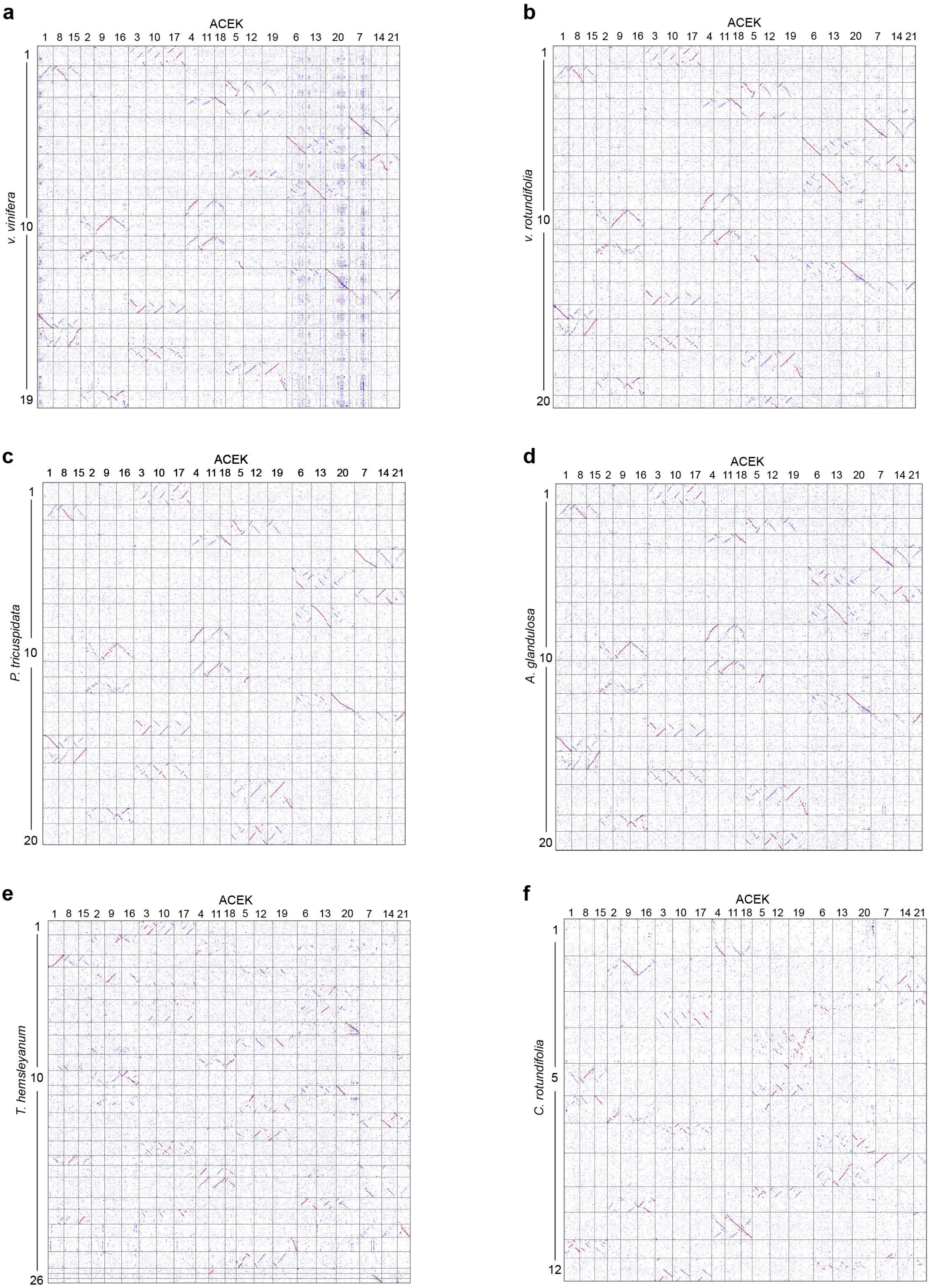


**Supplementary Fig. 10 | Chromosomal Synteny Analysis**: Dotplots of *V. vinifera* (a), *V. rotundifolia* (b), *P. tricuspidata* (c), *A. glandulosa* (d), *T. hemsleyanum* (e) and *C. rotundifolia* (f) compared to ACEK.

**a**

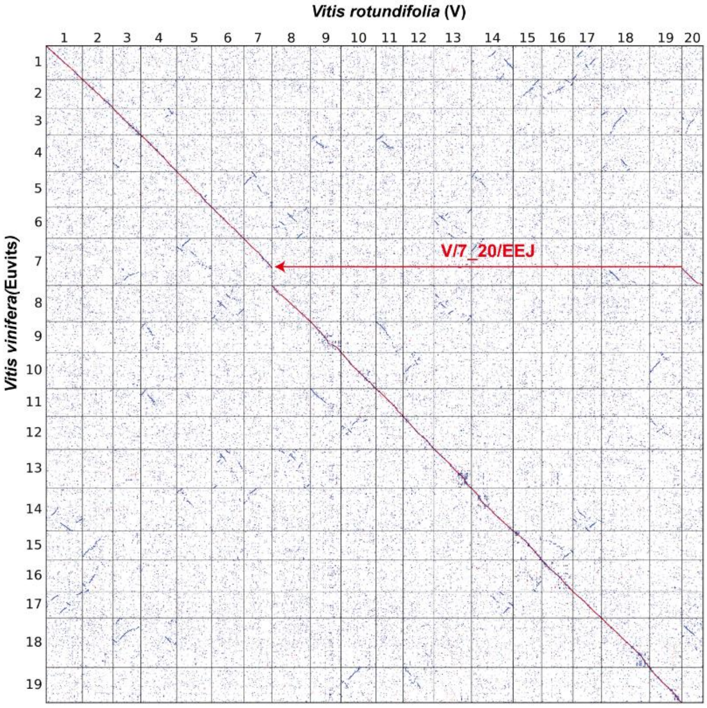


**b**

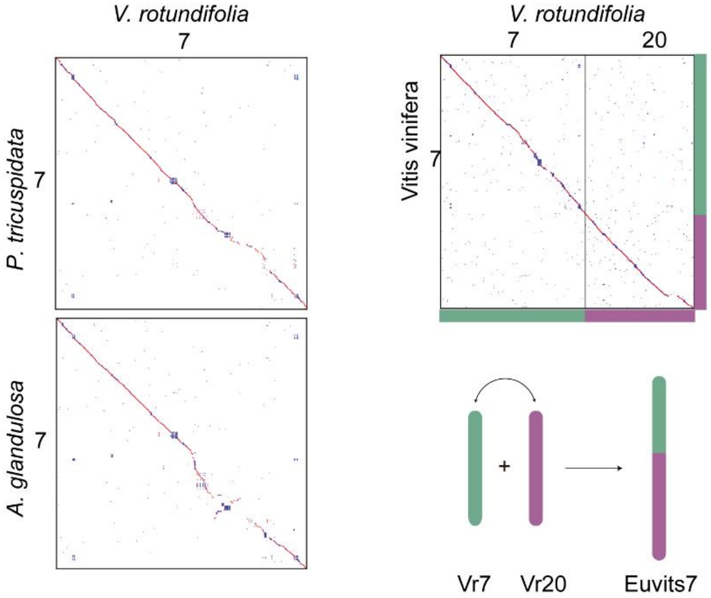


**c**

| V1 | V2 | V3 | V4 | V5 | V6 | V7 | V8 | V9 | V10 | V11 | V12 | V13 | V14 | V15 | V16 | V17 | V18 | V19 | V20 |
|---|---|---|---|---|---|---|---|---|---|---|---|---|---|---|---|---|---|---|---|
| Vr1 | Vr2 | Vr3 | Vr4 | Vr5 | Vr6 | Vr7 | Vr8 | Vr9 | Vr10 | Vr11 | Vr12 | Vr13 | Vr14 | Vr15 | Vr16 | Vr17 | Vr18 | Vr19 | Vr20 |

**Supplementary Fig. 11 Construction of ancestral *Vitis* karyotype(V).**

**a**. Genome-wide synteny comparison between *V. rotundifolia* (Vr) and *V. vinifera* (Euvitis). Chromosome 7 of *V. vinifera* (Euvitis) was formed by the EEJ fusion (V/7_20/EEJ) of Vr7 and Vr20.

**b.** Chromosome 7 is intact in *P. tricuspidata and A. glandulosa*, indicating that the fusion of Vr7 and Vr20 occurred after their divergence.

**c**. The *Vitis* karyotype (V) is consistent with that of *V. rotundifolia*.

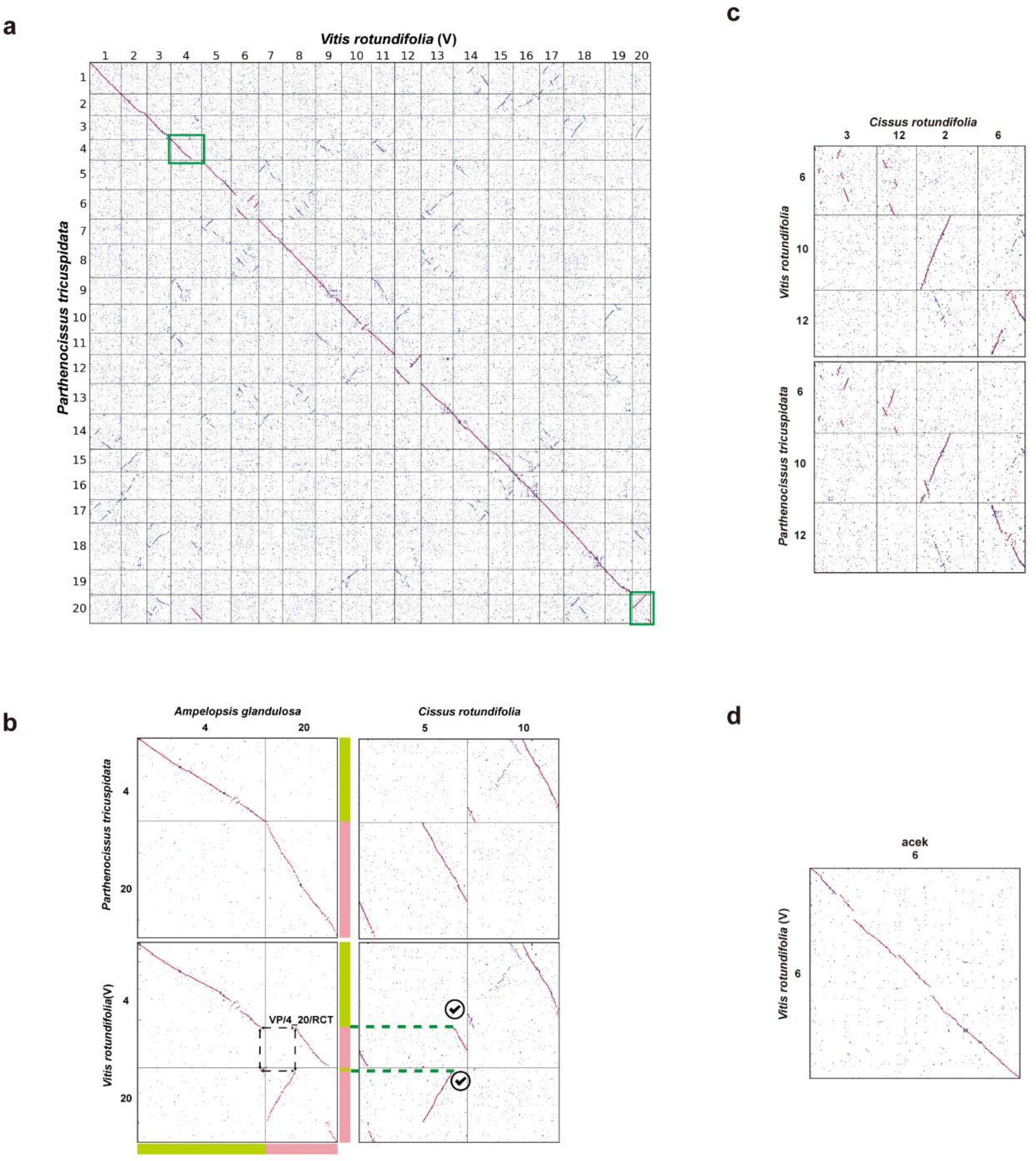


e

| | | | | | | | | | | | | | | | | | | | |
|---|---|---|---|---|---|---|---|---|---|---|---|---|---|---|---|---|---|---|---|
| VP1 | VP2 | VP3 | VP4 | VP5 | VP6 | VP7 | VP8 | VP9 | VP10 | VP11 | VP12 | VP13 | VP14 | VP15 | VP16 | VP17 | VP18 | VP19 | VP20 |
| Vr1 | Vr2 | Vr3 | Pa4 | Vr5 | Vr6 | Vr7 | Vr8 | Vr9 | Vr10 | Vr11 | Pa12 | Vr13 | Vr14 | Vr15 | Vr16 | Vr17 | Vr18 | Vr19 | Pa20 |

**Supplementary Fig. 12 Construction of the ancestral karyotype of *Vitis* and *Parthenocissus* (VP).**

**a**. Genome-wide synteny comparison between the *Vitis* karyotype (V) and *P. tricuspidata*. Chromosomal rearrangements were detected involving chromosomes 4 and 20.

**b.** Based on the shared karyotype structure among *V. rotundifolia*, *A. glandulosa* (Ag), and *P. tricuspidata* (Pa), chromosomes 4 and 20 of V were inferred to have formed later. Accordingly, chromosomes Pa4 and Pa20 were inferred as the ancestral VP chromosomes.

**c.** Internal structural variations were also observed on chromosomes 6, 10, and 12 between V and pa. Pa10 and Pa12 are consistent with the karyotype of *V. rotundifolia* and were therefore inferred as ancestral VP chromosomes.

**d.** Ancestral chromosome status of Vr6 was further validated using ACEK analysis.

**e.** Origin and chromosomal composition of each inferred VP ancestral chromosome.

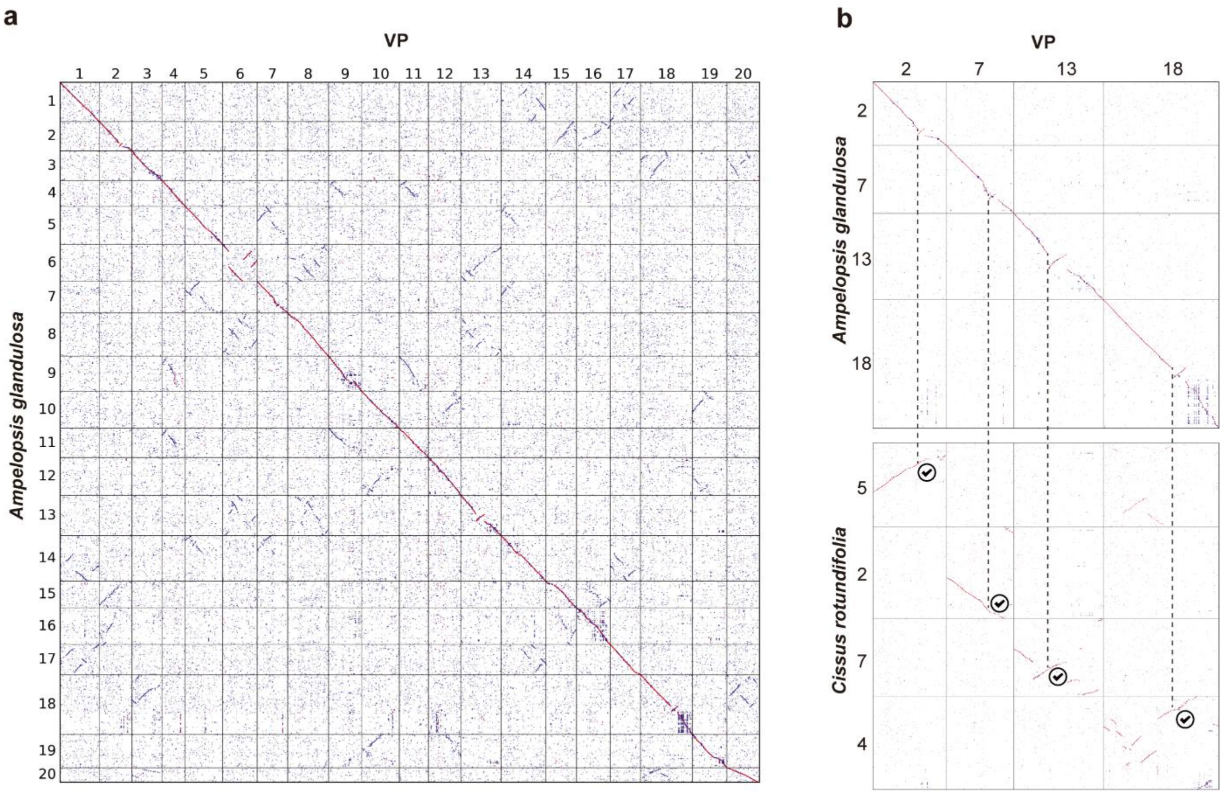


c

| | | | | | | | | | | | | | | | | | | | |
|---|---|---|---|---|---|---|---|---|---|---|---|---|---|---|---|---|---|---|---|
| VPA1 | VPA2 | VPA3 | VPA4 | VPA5 | VPA6 | VPA7 | VPA8 | VPA9 | VPA10 | VPA11 | VPA12 | VPA13 | VPA14 | VPA15 | VPA16 | VPA17 | VPA18 | VPA19 | VPA20 |
| Vr1 | Vr2 | Vr3 | Pt4 | Vr5 | Vr6 | Vr7 | Vr8 | Vr9 | Vr10 | Vr11 | Pt12 | Vr13 | Vr14 | Vr15 | Vr16 | Vr17 | Vr18 | Vr19 | Pt20 |

**Supplementary Fig. 13 Construction of ancestral VP and *A. glandulosa* karyotype (VPA).**

**a**. Genome-wide synteny comparison between VP and *A. glandulosa*. Structural variations were detected on chromosomes 2, 7, 13, and 18 between them.

**b.** Homologous chromosomes corresponding to VP chromosomes 2, 7, 13, and 18 were extracted from *A. glandulosa* and *C. rotundifolia*. Arrowheads indicate breakpoint positions between VP and *A. glandulosa*, whereas the chromosomal structures in *Cissus* are consistent with those of VP.

**c**. Origin and chromosomal composition of each inferred VPA ancestral chromosome.

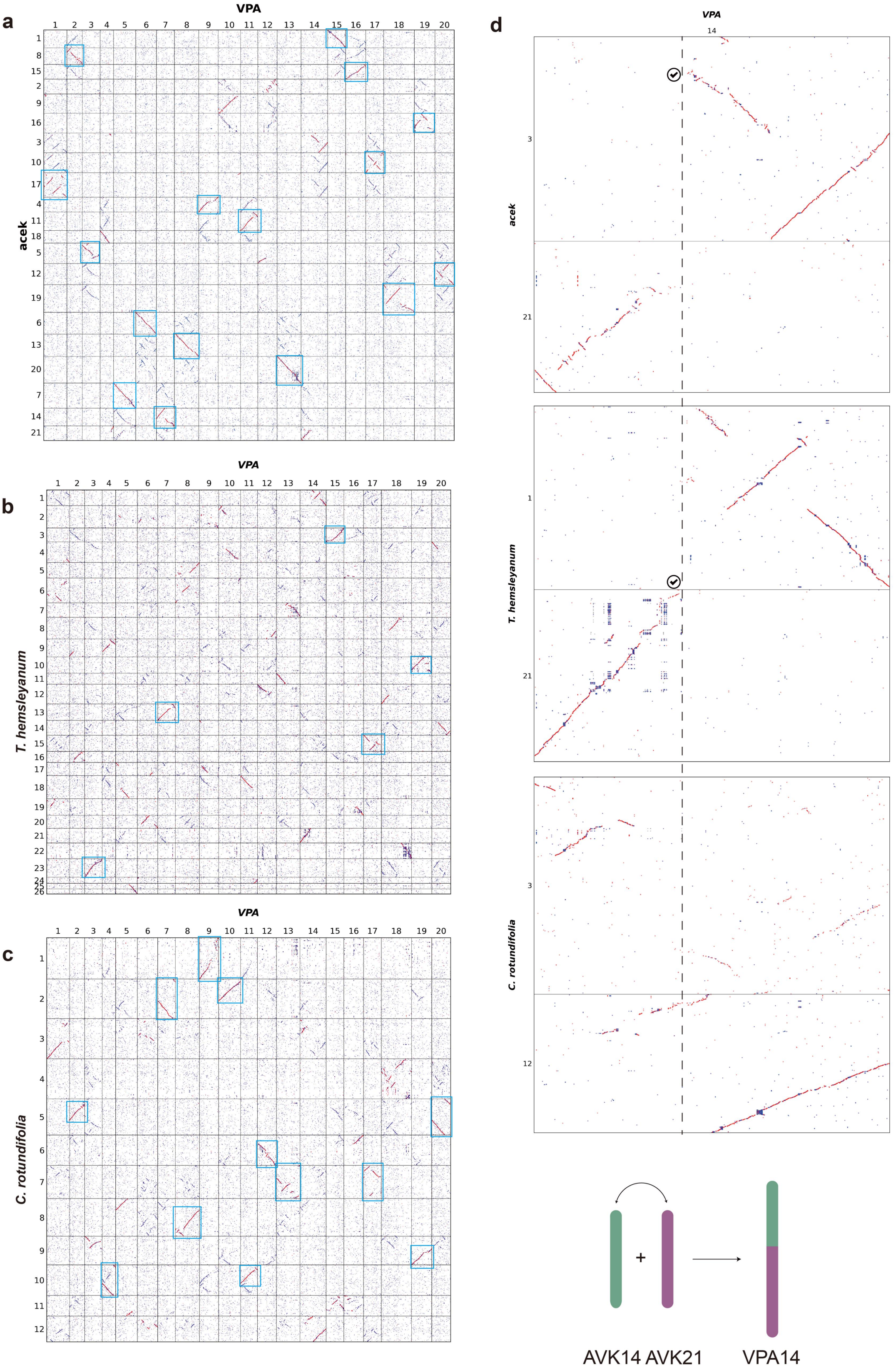


e

| | | | | | | | | | | | | | | | | | | | | |
|---|---|---|---|---|---|---|---|---|---|---|---|---|---|---|---|---|---|---|---|---|
| AVK1 | AVK2 | AVK3 | AVK4 | AVK5 | AVK6 | AVK7 | AVK8 | AVK9 | AVK10 | AVK11 | AVK12 | AVK13 | AVK14 | AVK15 | AVK16 | AVK17 | AVK18 | AVK19 | AVK20 | AVK21 |
| Vr1 | Vr2 | Vr3 | Pt4 | Vr5 | Vr6 | Vr7 | Vr8 | Vr9 | Vr10 | Vr11 | Pt12 | Vr13 | Vr14* | Vr15 | Vr16 | Vr17 | Vr18 | Vr19 | Pt20 | Th21 |

**Supplementary Fig. 14 Construction of ancestral Vitaceae karyotype (AVK).** Genome-wide synteny comparison among ACEK(**a**), *T. hemsleyanum* (**b**), *C. rotundifolia* (**c**), and the VPA ancestral karyotype. Except for chromosome 14, all VPA chromosomes have karyotypically consistent counterparts in one of the other three genomes. **d.** Synteny comparison of chromosome 14 in the VPA karyotype with the homologous chromosomes from ACEK, *T. hemsleyanum*, and *C. rotundifolia*. *T. hemsleyanum* and ACEK share identical breakpoint positions, indicating that chromosome 14 of VPA was formed by an RCT fusion of two ancestral AVK chromosomes. AVK21 corresponds to Th21, whereas AVK14 is consistent with the distal segment of VPA chromosome 14.**e.** Origin and chromosomal composition of each inferred AVK ancestral chromosome. Vr14* represents genes Vr14_1 to Vr14_1011.

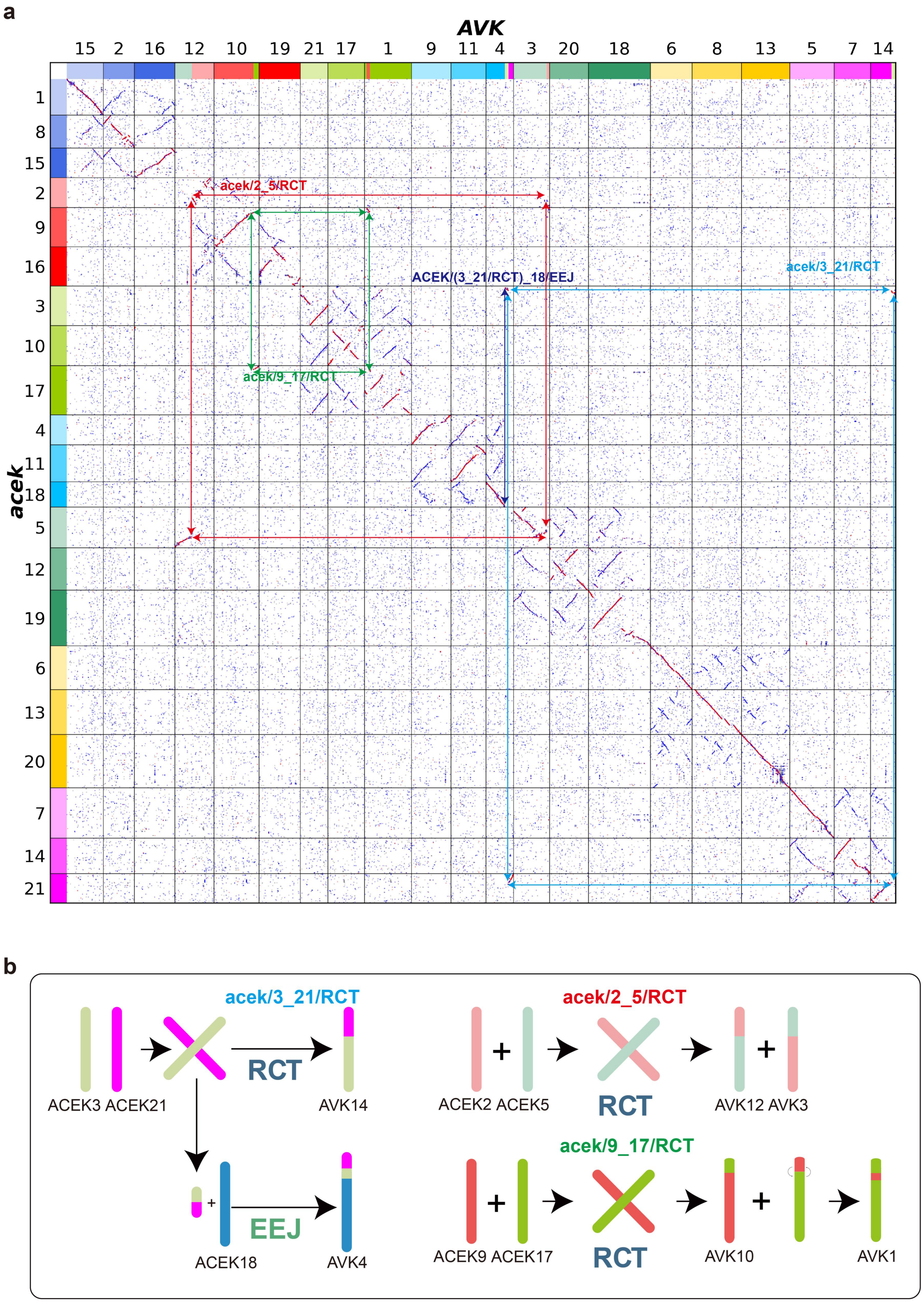


**Supplementary Fig. 15 Reconstruction of chromosome evolution in the ancestral AVK karyotype.**

a. Genome-wide synteny comparison between ACEK and the AVK. Red, green, and blue arrows indicate representative RTC-mediated rearrangements involving ACEK chromosome pairs (ACEK2/5, ACEK9/17, and ACEK3/21) and EEJ(ACEK3_21/18), which contributed to the formation of AVK chromosomes.

b. Schematic illustration of the origin and chromosomal composition of AVK chromosomes inferred from synteny and breakpoint analyses.

a

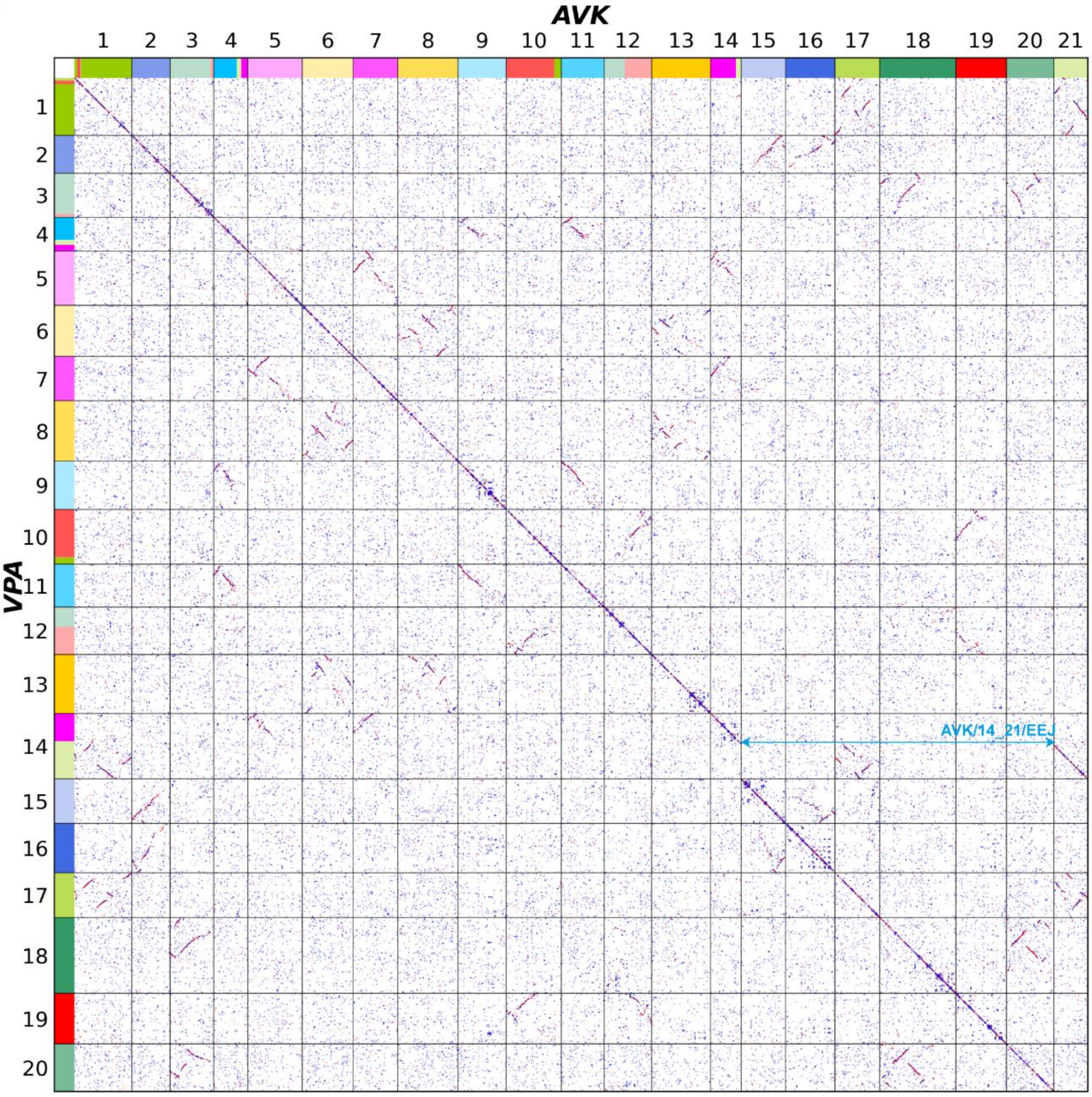


b

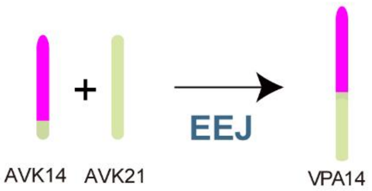


**Supplementary Fig. 16 Evolutionary reconstruction of the transition from AVK to VPA.**

**a.** Genome-wide synteny comparison between the VPA and the inferred AVK. Strong collinearity is observed across most chromosomes. A clear end-to-end joining (EEJ) signal is detected between AVK14 and AVK21.

**b.** Schematic illustration of the formation of the VPA chromosome 14. VPA14 was generated by an EEJ-mediated fusion of AVK14 and AVK21.

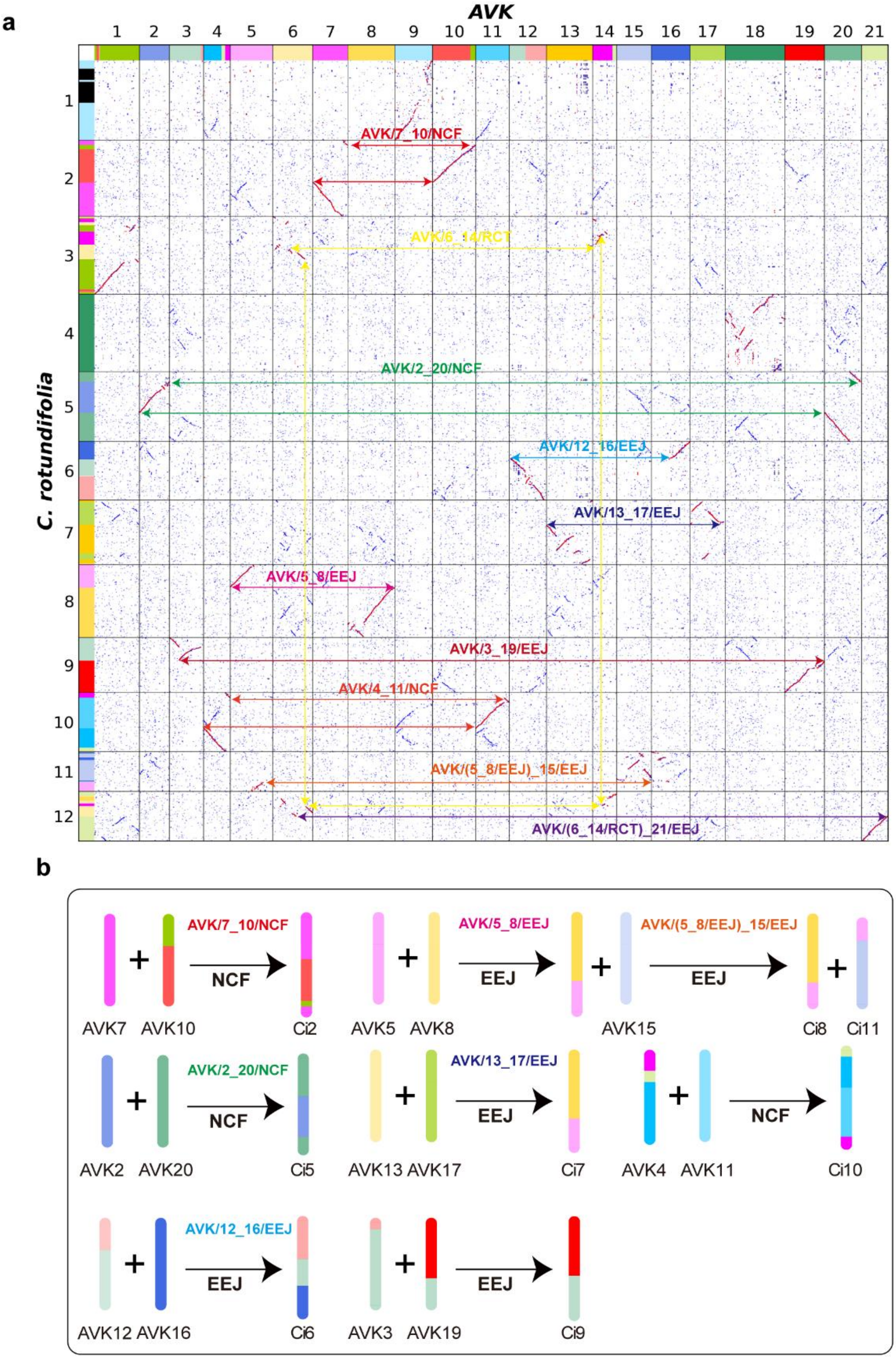


**Supplementary Fig. 17 Evolutionary reconstruction of the transition from AVK to *C. rotundifolia*.**

**a.** Genome-wide synteny comparison between *Cissus rotundifolia* and the inferred AVK ancestral karyotype. Conserved syntenic blocks and breakpoint patterns reveal multiple chromosomal rearrangement events, including NCF and EEJ.

**b.** Schematic reconstruction of the chromosomal rearrangements underlying the transition from AVK to *C. rotundifolia*.

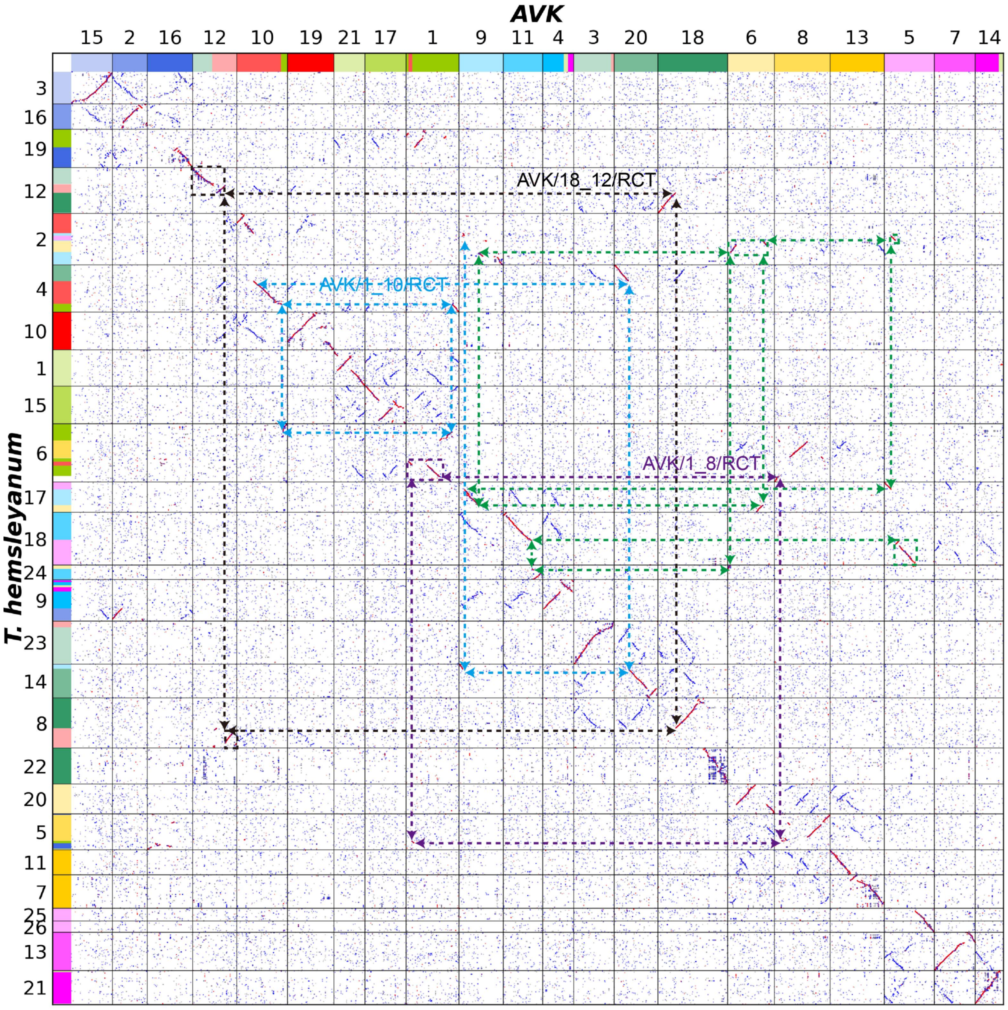


**Supplementary Fig. 18 | Breakpoint analysis of chromosomal fusion events from AVK to *T. hemsleyanum*.**

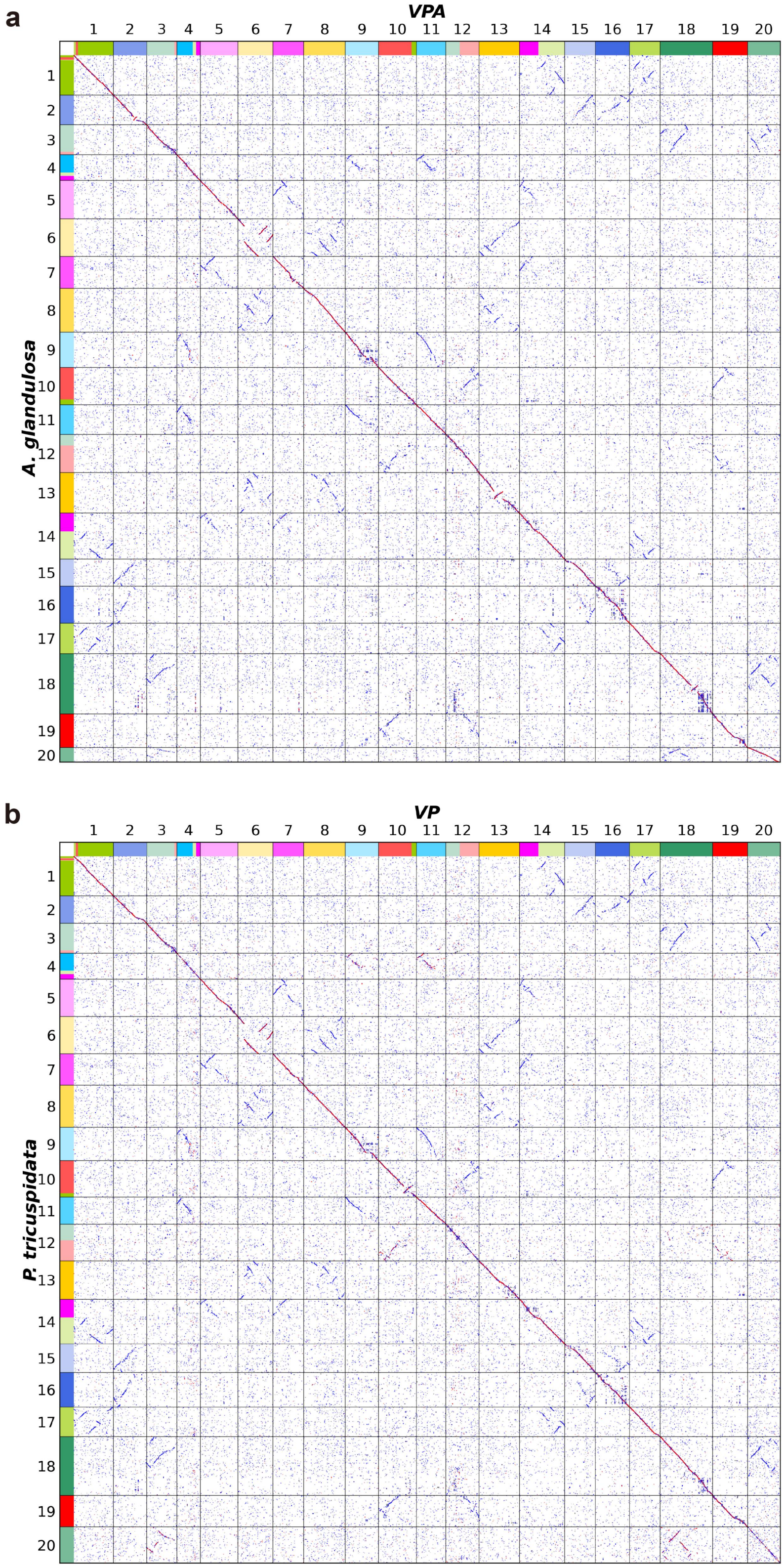


**Supplementary Fig. 19 | Chromosome-level synteny plots between *A. glandulosa* and VPA (a),** and between *P. tricuspidata* and VP **(b),** indicate highly conserved karyotypes without detectable chromosomal rearrangements.

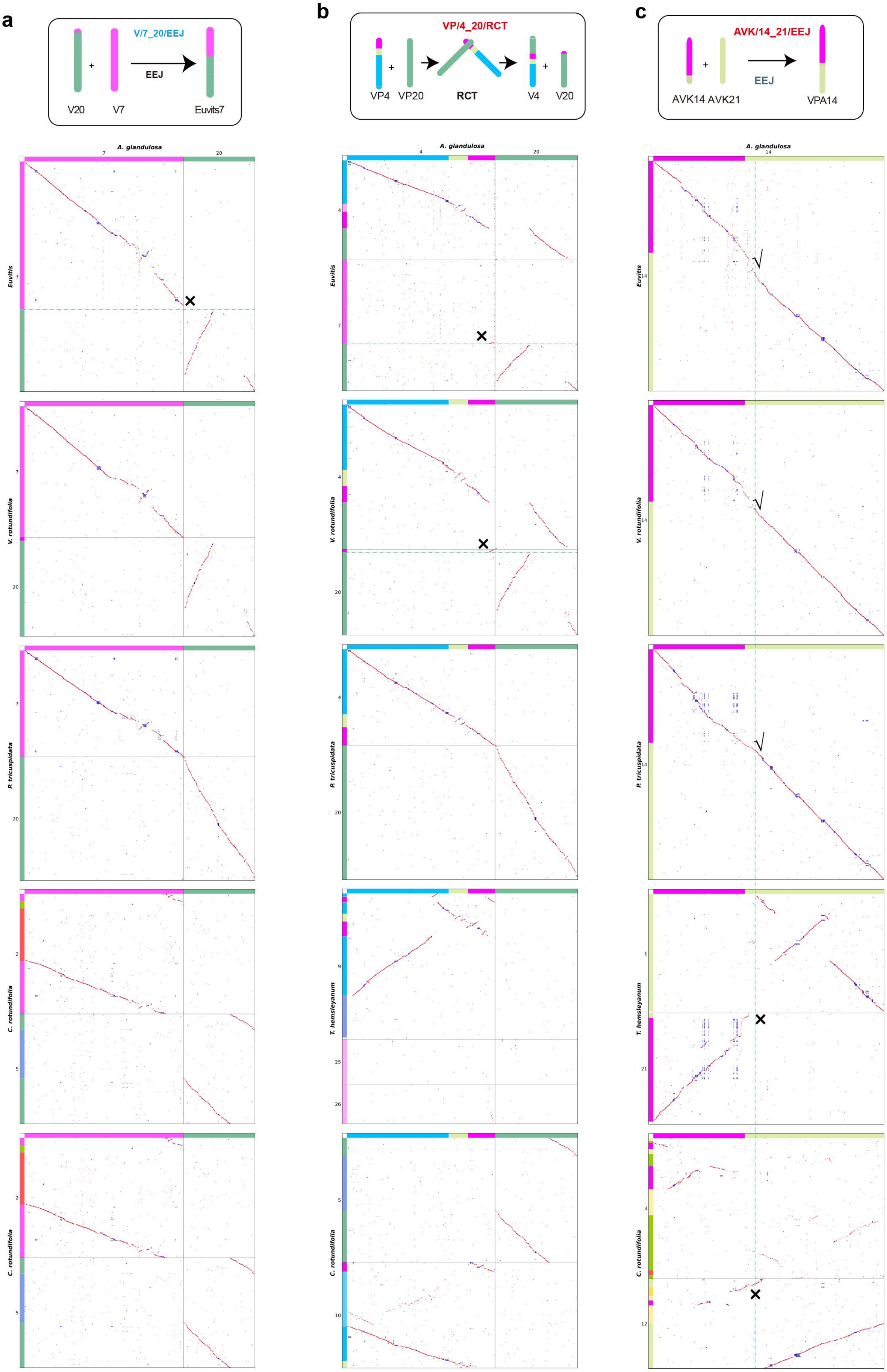


**Supplementary Fig. 20 | Inference of phylogenetic relationships based on shared chromosomal breakpoints involving *A. glandulosa*. a.** A chromosomal fusion event detected exclusively in Euvitis **b**. A chromosomal fusion event shared by Euvitis and *V. rotundifolia*. **c**. A chromosomal fusion event conserved in four species, being present in all analyzed samples except *T. hemsleyanum* and *C. rotundifolia*, suggesting that this rearrangement occurred after the divergence of *T. hemsleyanum* and *C. rotundifolia*.

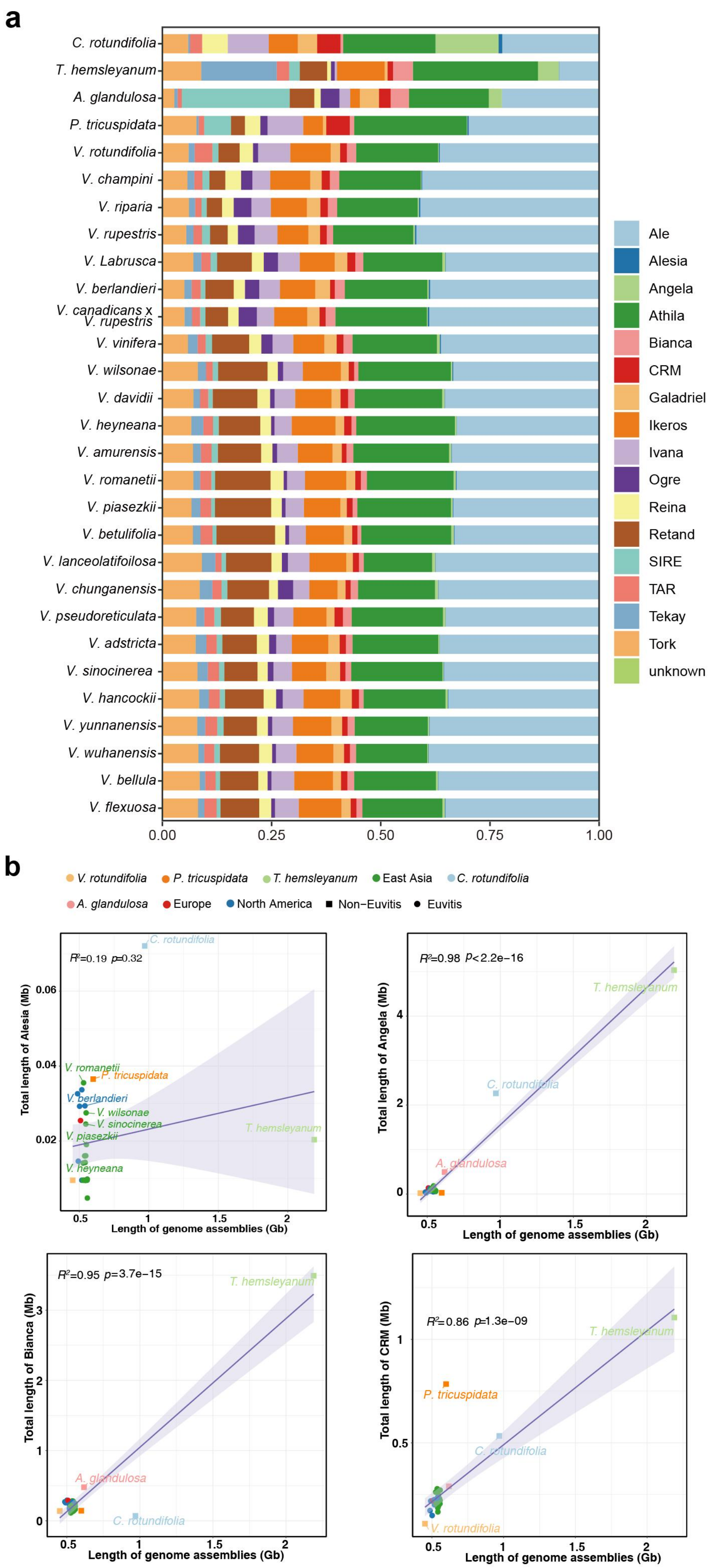


**Supplementary Fig. 21 |** Phenotypes of selected *Vitis* accessions and TE annotations: **a**. Proportion of various intact LTRs; **b**. Relationship between Alesia, Angela, Blanca, CRM, and genome size.

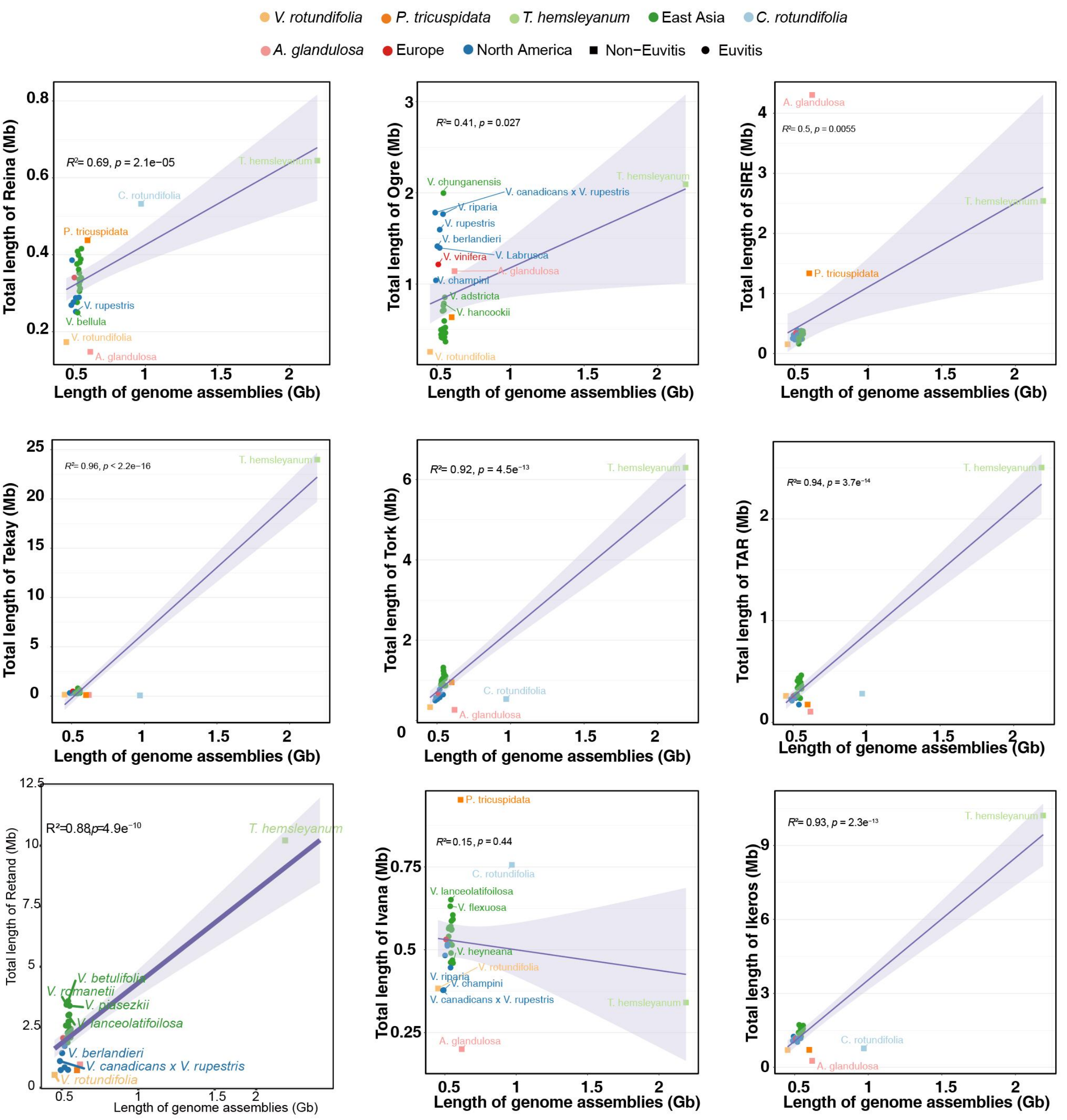


**Supplementary Fig.22 |** Relationship between Reina, Ogre, SIRE, Tekay, Tork, TAR, Galadriel, Ivana, Ikeros and genome size.

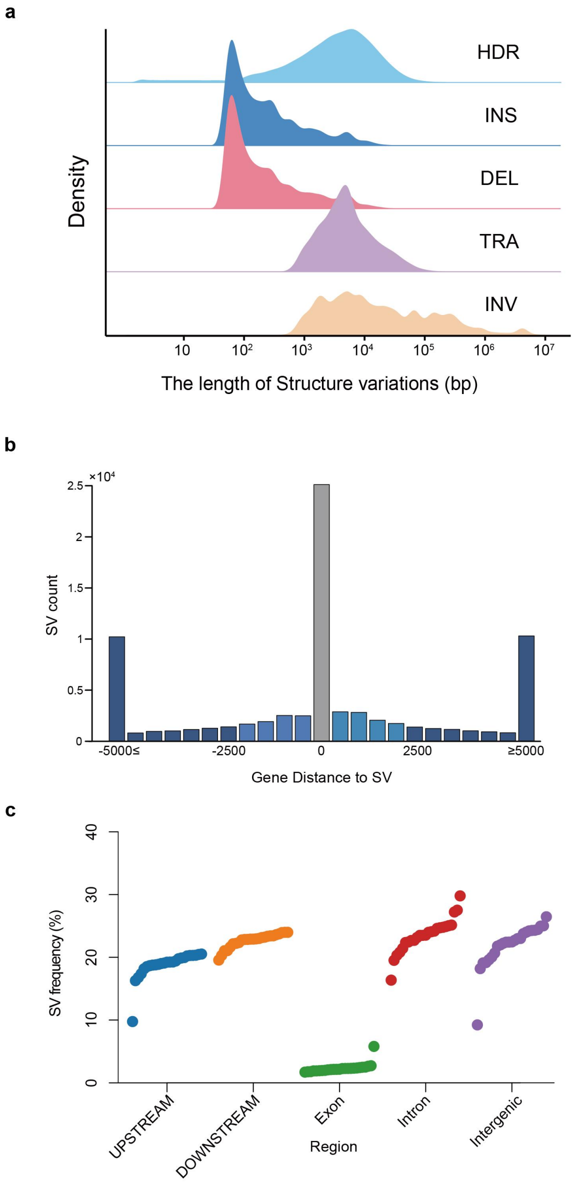


**Supplementary Fig. 23 |** The landscape of structural variants. **a.** Size and frequency of various types of structural variations. **b**. The relationship between genes and structural variation distance. **c**. Frequency of structural variations occurring in each gene interval.

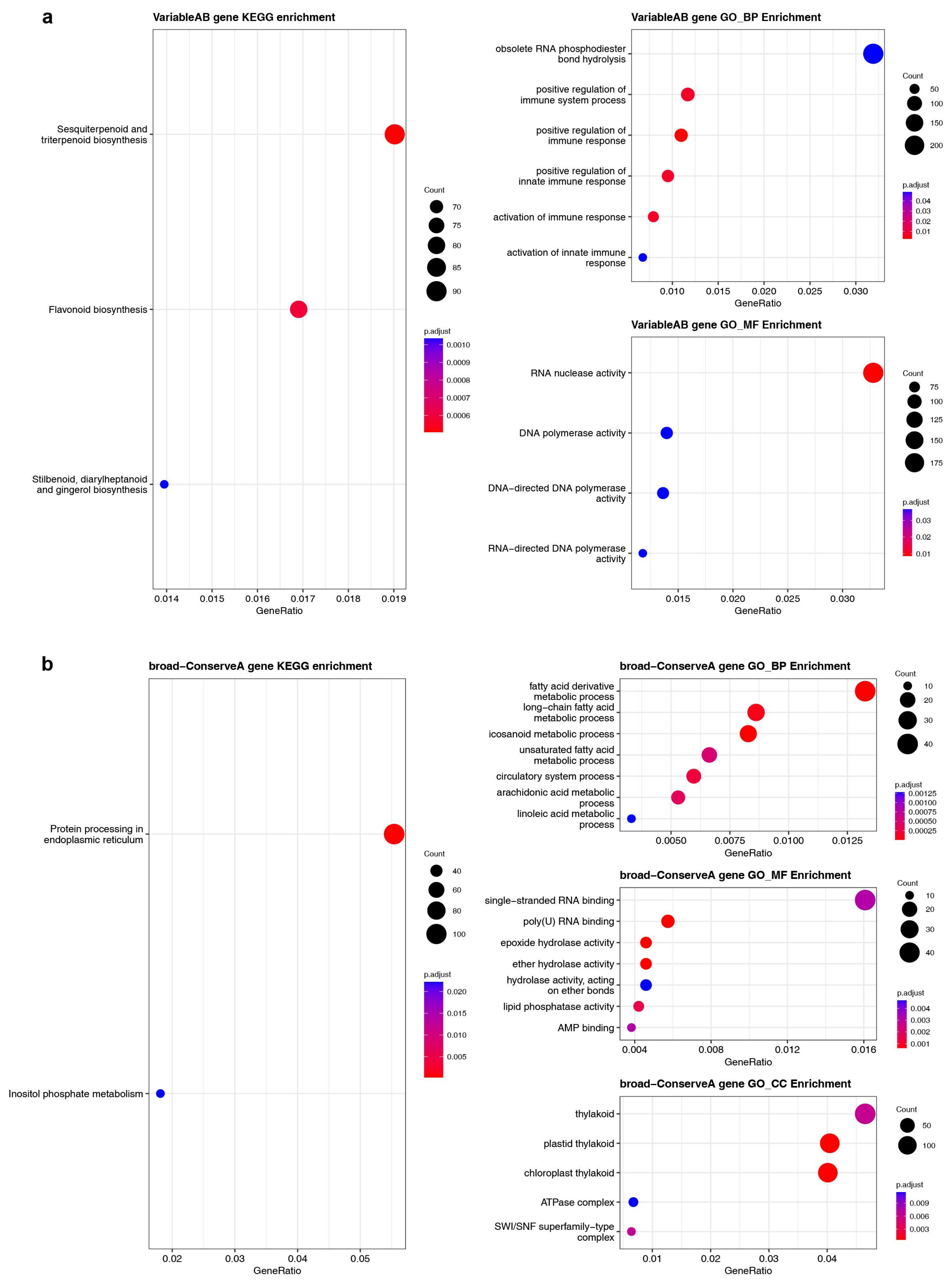


**Supplementary Fig. 24 |** KEGG and GO functional enrichment of genes in Variable A/B (**a**) and Broad-Conserve B (Conserve B and Dispensable B) (**b**).

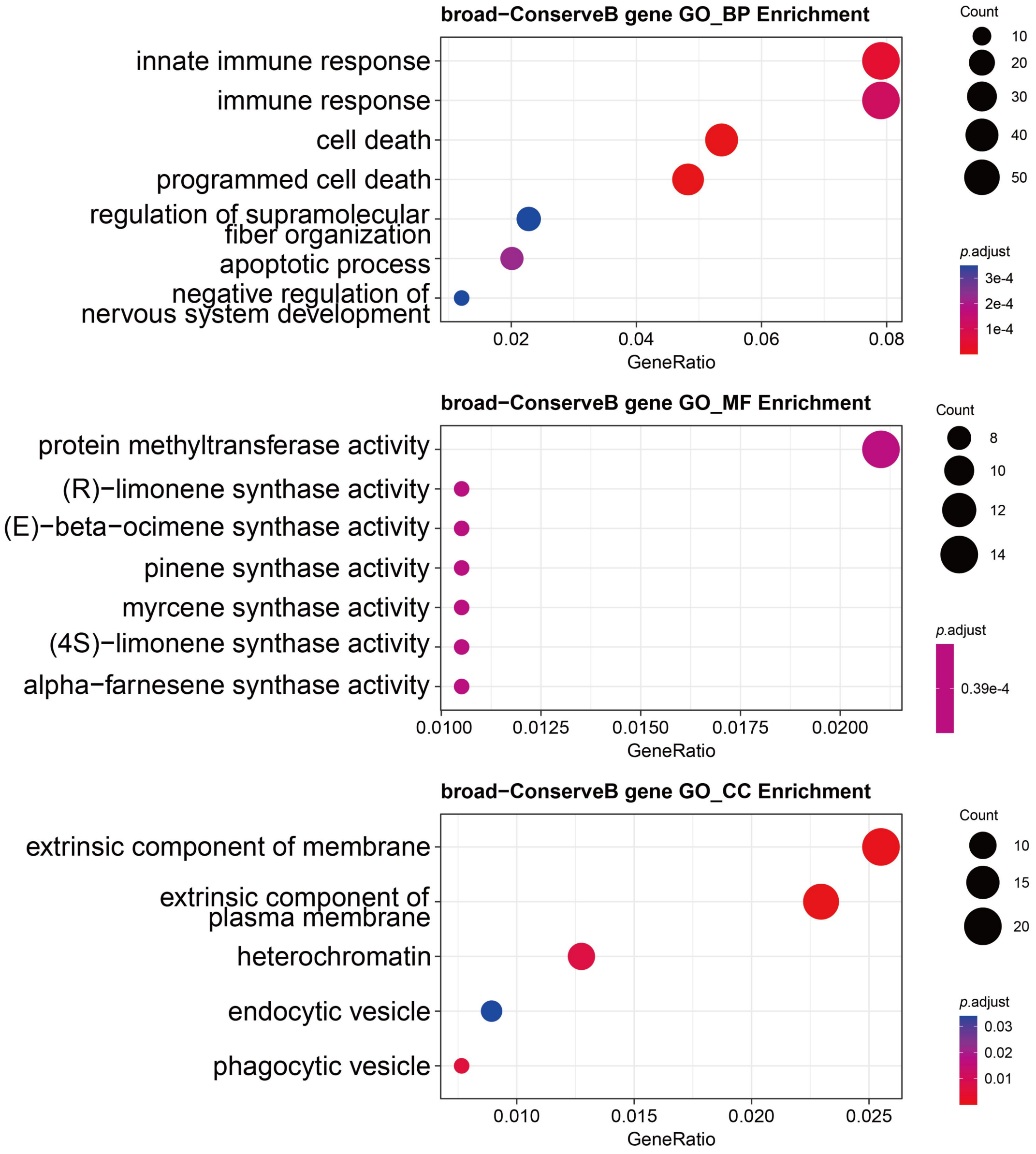


**Supplementary Fig. 25** |KEGG and GO functional enrichment of genes in Broad-Conserve B (Conserve B and Dispensable B).

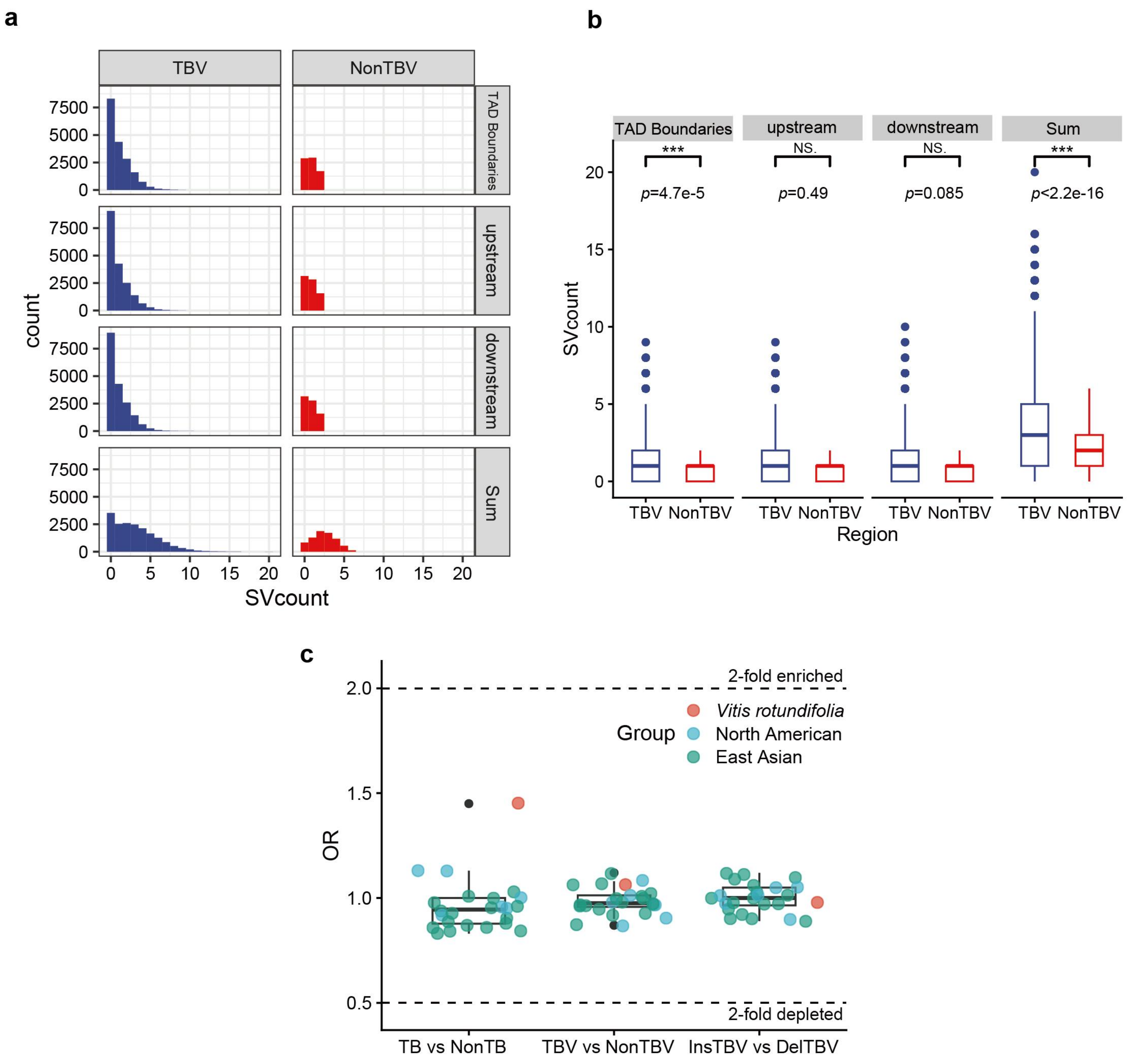


**Supplementary Fig. 26** | Structural variants associated with TAD boundary variation (TBV). **a**, Distribution of structural variants (SVs) at TBVs and non-TBVs, partitioned into the boundary itself, 50 kb upstream, and 50 kb downstream, as well as combined regions (TBV + upstream 50 kb; TBV + downstream 50 kb). **b**, Statistical comparison of SV abundance between TBVs and non-TBVs, assessed by two-tailed t-tests. Significance levels are indicated by *P* values. **c**, Comparative analysis of TAD boundary–associated SVs among grapevine populations, with enrichment significance evaluated by odds ratio (OR) tests.

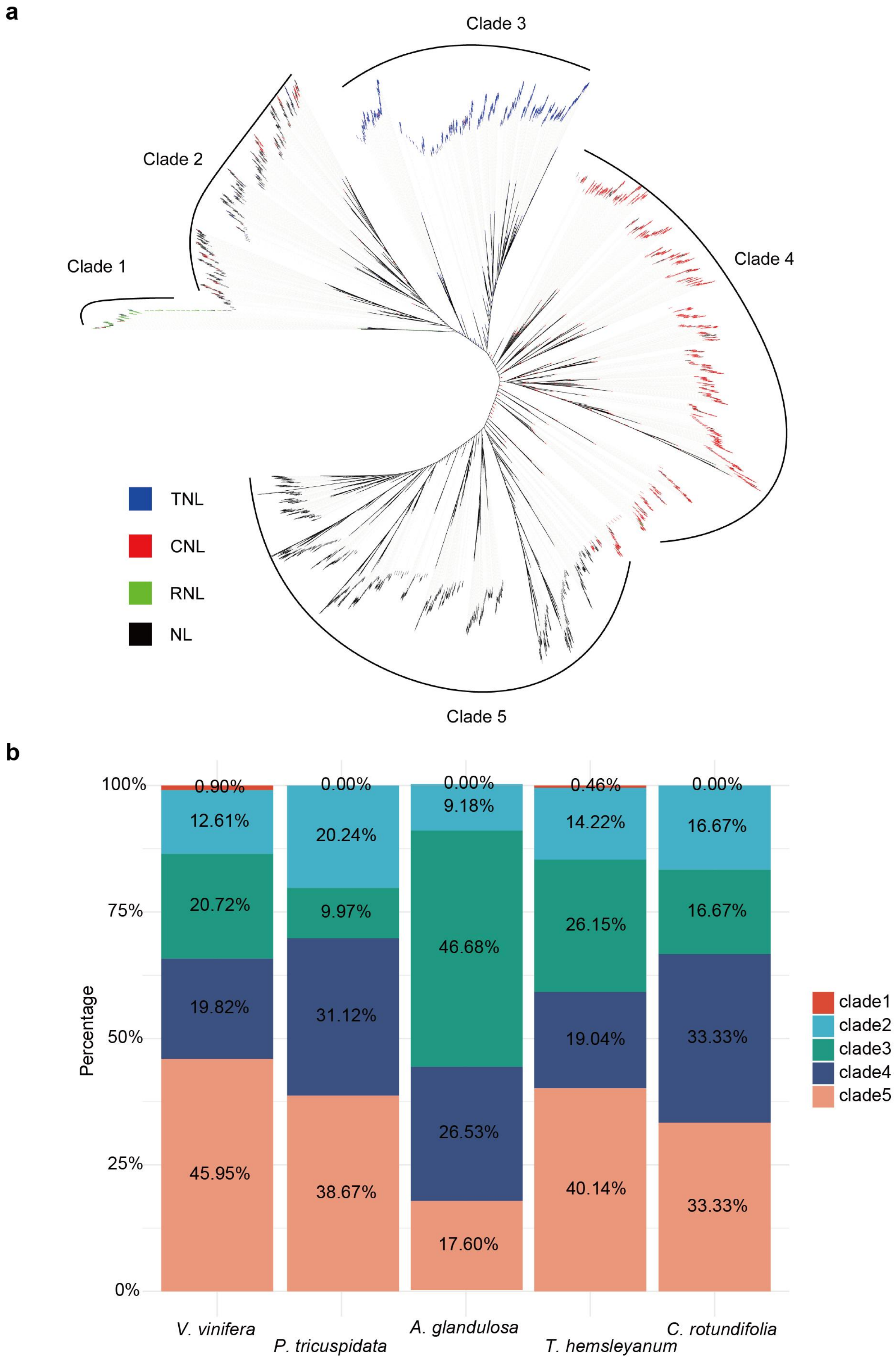


**Supplementary Fig. 27 |** A phylogenetic tree using NB-ARC domain sequences across the Vitaceae family(**a**). The relative proportions of different NLR clades are shown for each species (**b**).

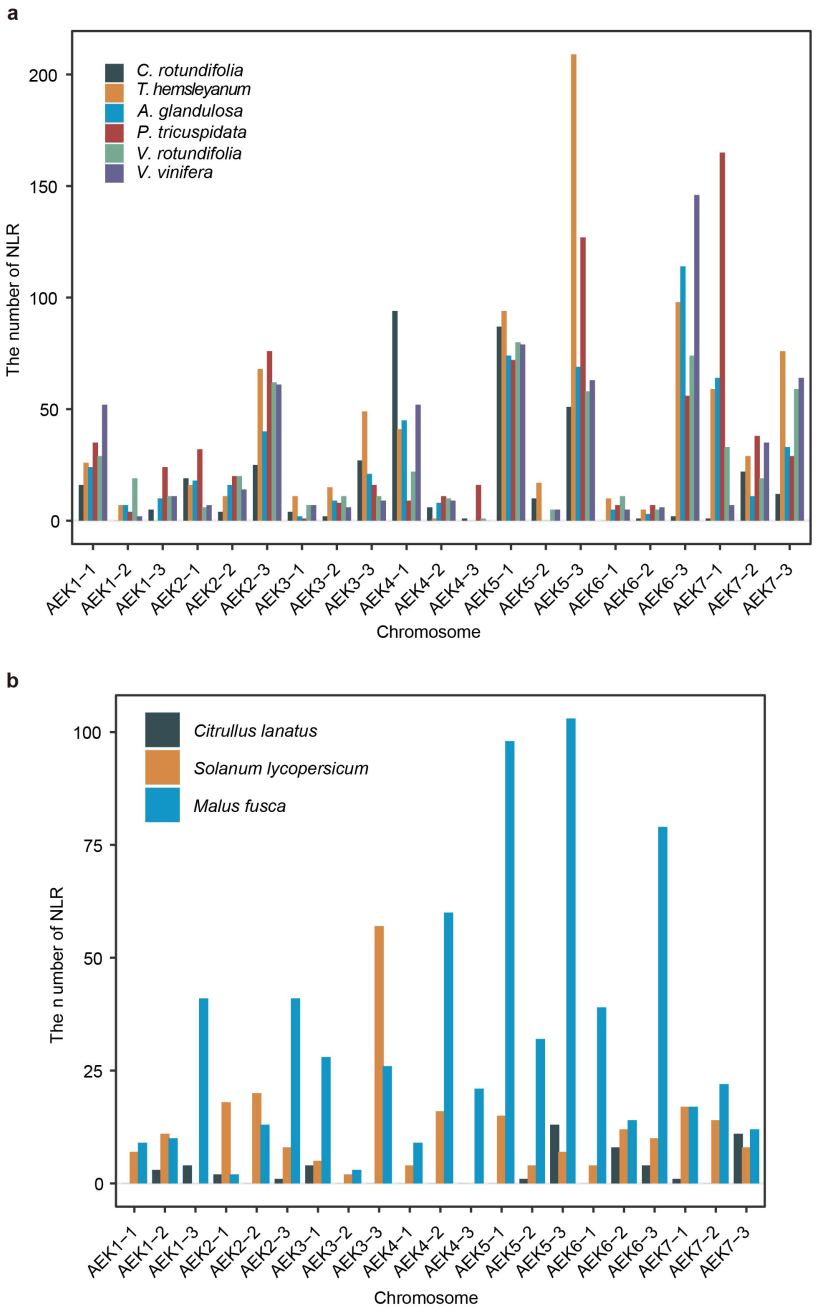


**Supplementary Fig. 28 |** The number of NLR genes on ancestral chromosomal segments. The number of NLR genes on ancestral chromosomal segments. **a.** The distribution of NLRs on ancestral chromosomal segments in 6 Vitaceae species.NLR genes identified in each genome were assigned to ancestral chromosomal segments based on conserved synteny relationships. **b.** The distribution of NLRs on ancestral chromosomal segments in *Citrullus lanatus, Solanum lycopersicum* and *Malus fusca,* shown for comparison with Vitaceae species. Ancestral chromosomal segments are labeled according to their origin from the ancestral eudicot karyotype (AEK), including subchromosomal segments generated by subsequent chromosomal rearrangements. Bar heights indicate the number of NLR genes per ancestral segment. AEK labels with numerical suffixes (for example, AEK1-2-3) denote distinct subchromosomal segments derived from the same ancestral core eudicot chromosome (ACEK1) following the core eudicot-whole-genome triplication (hexaploidization, $\gamma$ event).

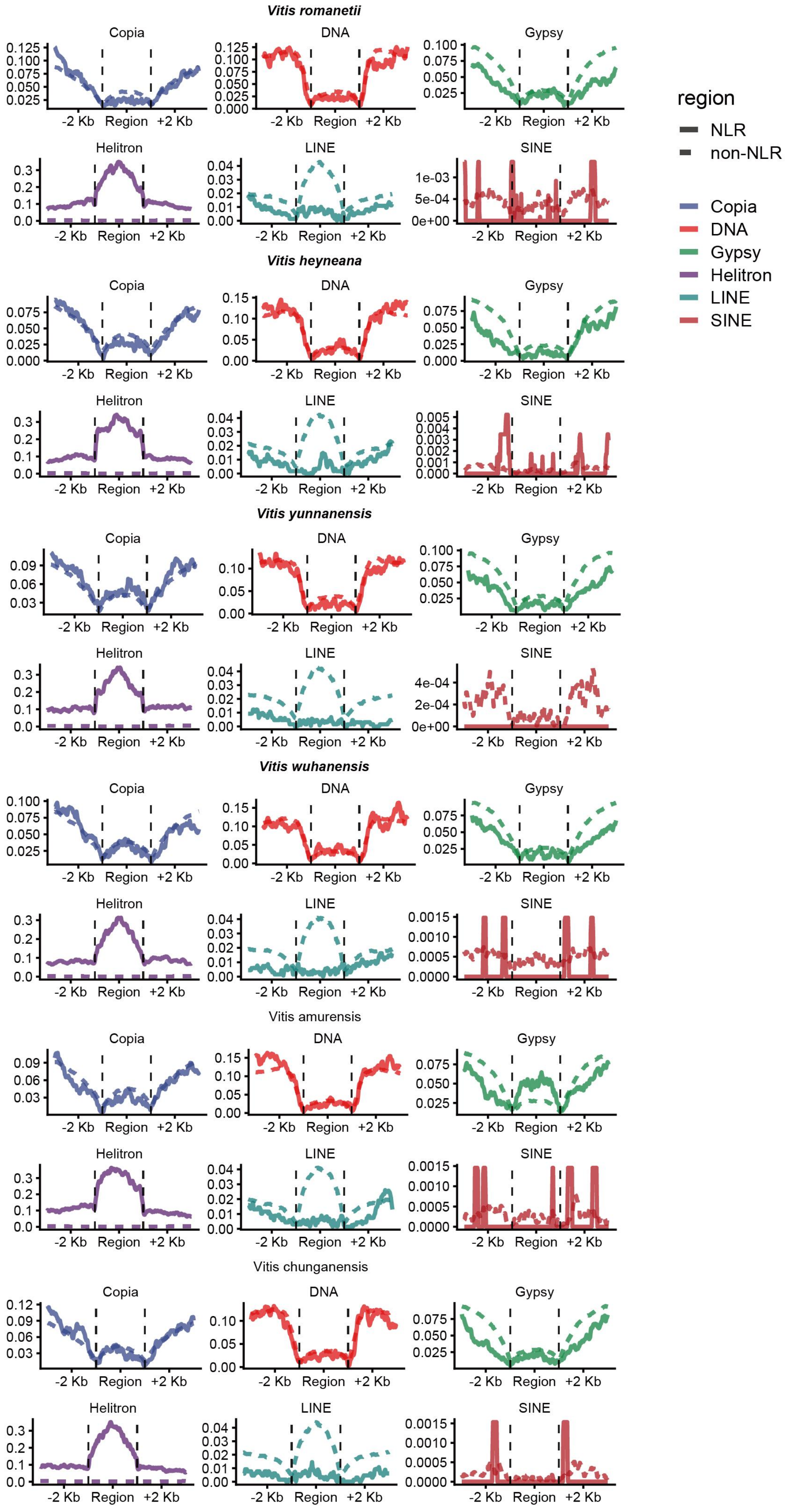


**Supplementary Fig. 29 |** The TE frequency within the 2 kb upstream and downstream regions of NLR and non-NLR genes in *Vitis romanetii, Vitis heyneana, Vitis yunnanensis, Vitis wuhanensis, Vitis amurensis* and *Vitis chunganensis*.

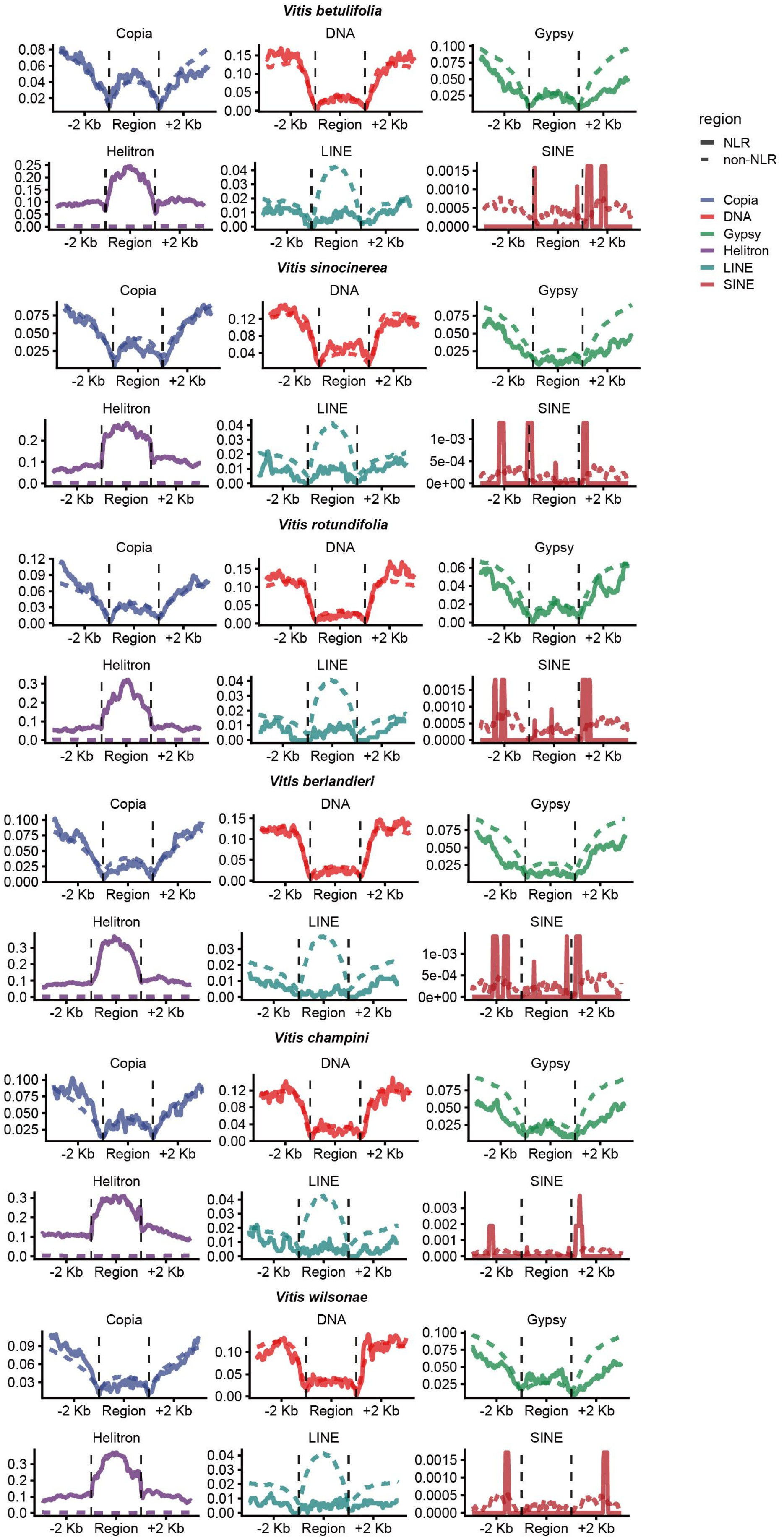


**Supplementary Fig. 30 |** The TE frequency within the 2 kb upstream and downstream regions of NLR and non-NLR genes in *Vitis betulifolia, Vitis sinocinerea, Vitis rotundifolia, Vitis berlandieri, Vitis champini,* and *Vitis wilsonae.*

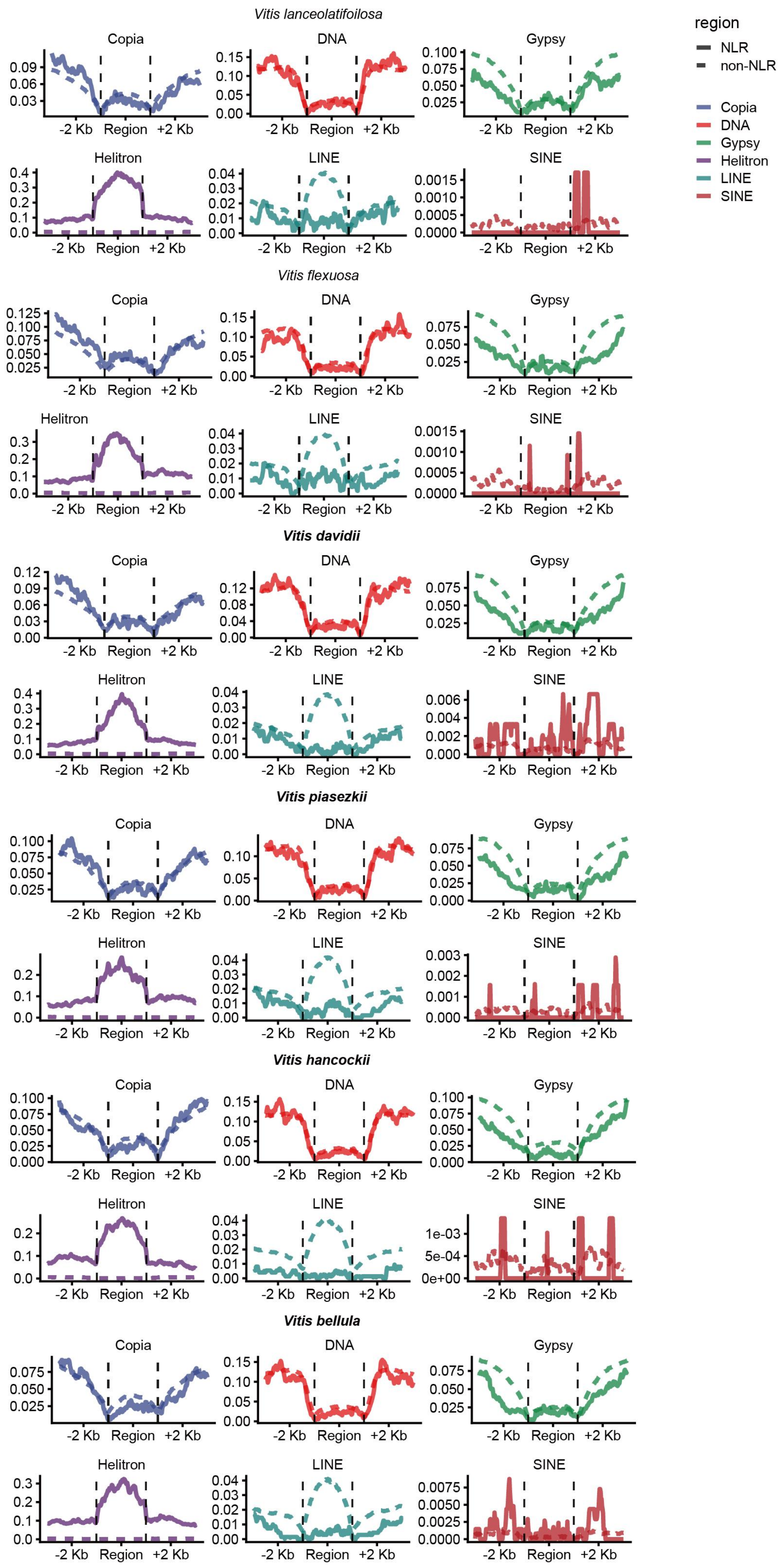


**Supplementary Fig. 31 |** The TE frequency within the 2 kb upstream and downstream regions of NLR and non-NLR genes in *Vitis lanceolatifoilosa, Vitis flexuosa, Vitis davidii, Vitis piasezkii, Vitis hancockii,* and *Vitis bellula.*

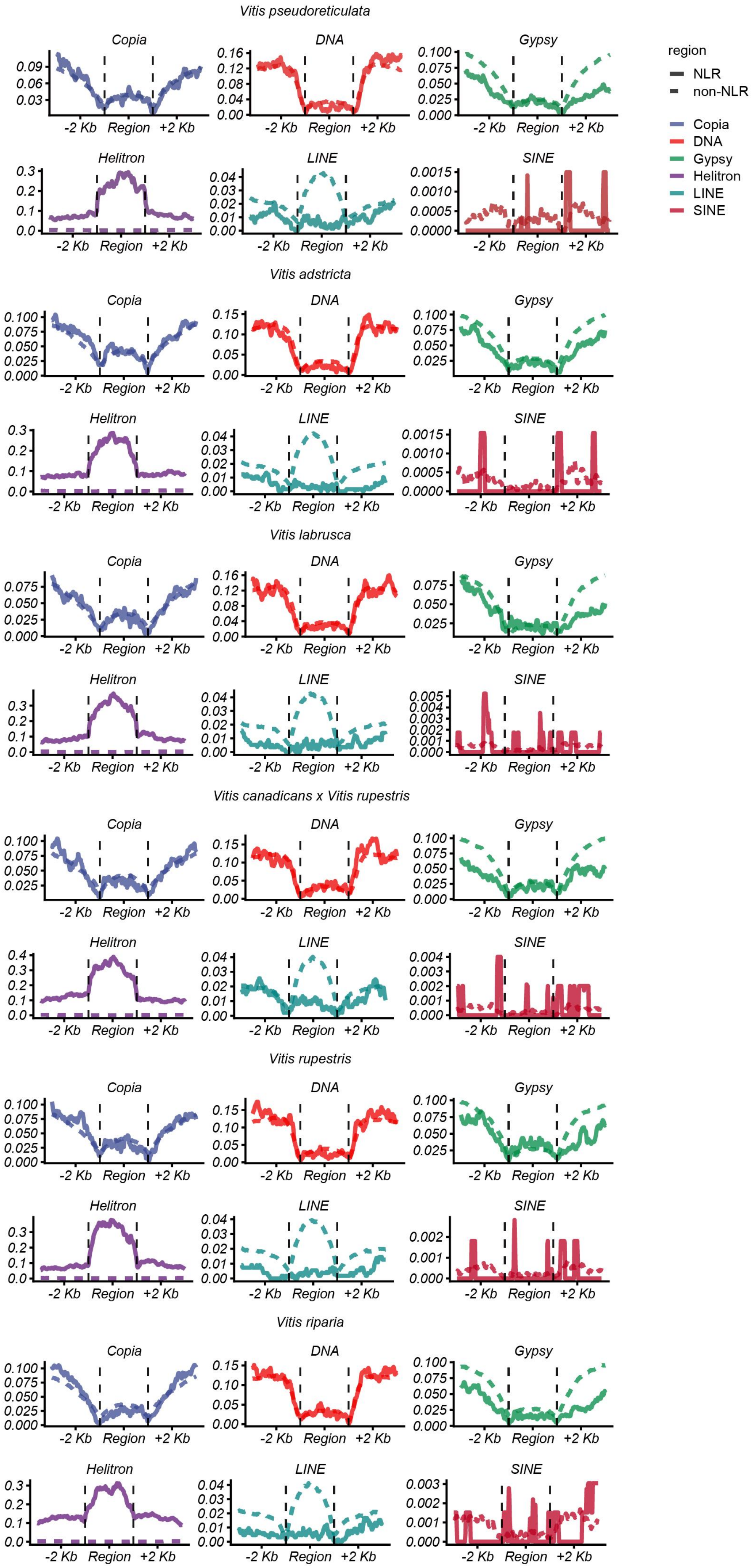


**Supplementary Fig. 32 |** The TE frequency within the 2 kb upstream and downstream regions of NLR and non-NLR genes in *Vitis pseudoreticulata, Vitis adstricta, Vitis labrusca, Vitis canadicans x Vitis rupestris, Vitis rupestris and Vitis riparia.*

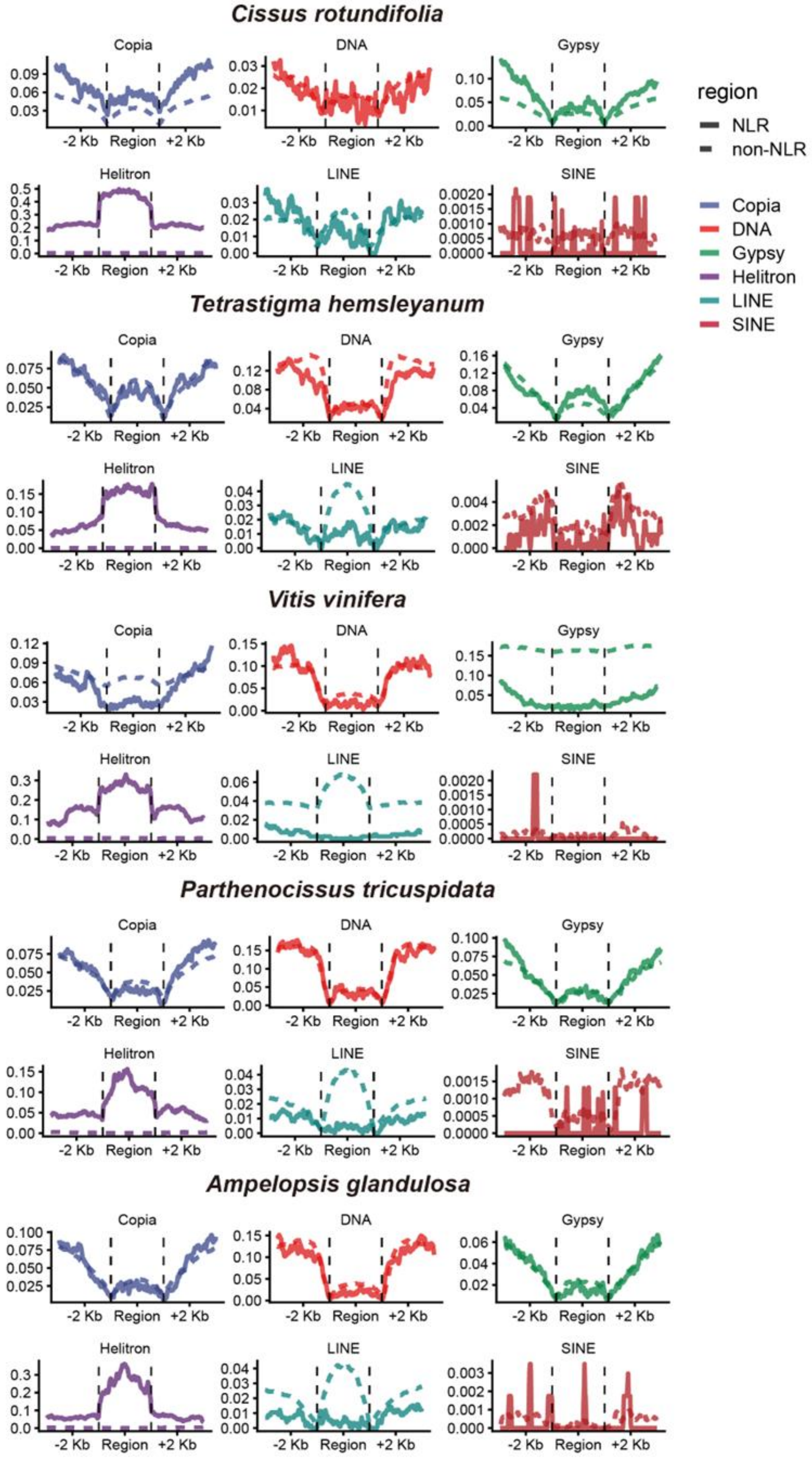


**Supplementary Fig. 33 |** The TE frequency within the 2 kb upstream and downstream regions of NLR and non-NLR genes in *C. rotundifolia*, *T. hemsleyanum*, *Vitis vinifera*, *P. tricuspidata*, and *A. glandulosa*.

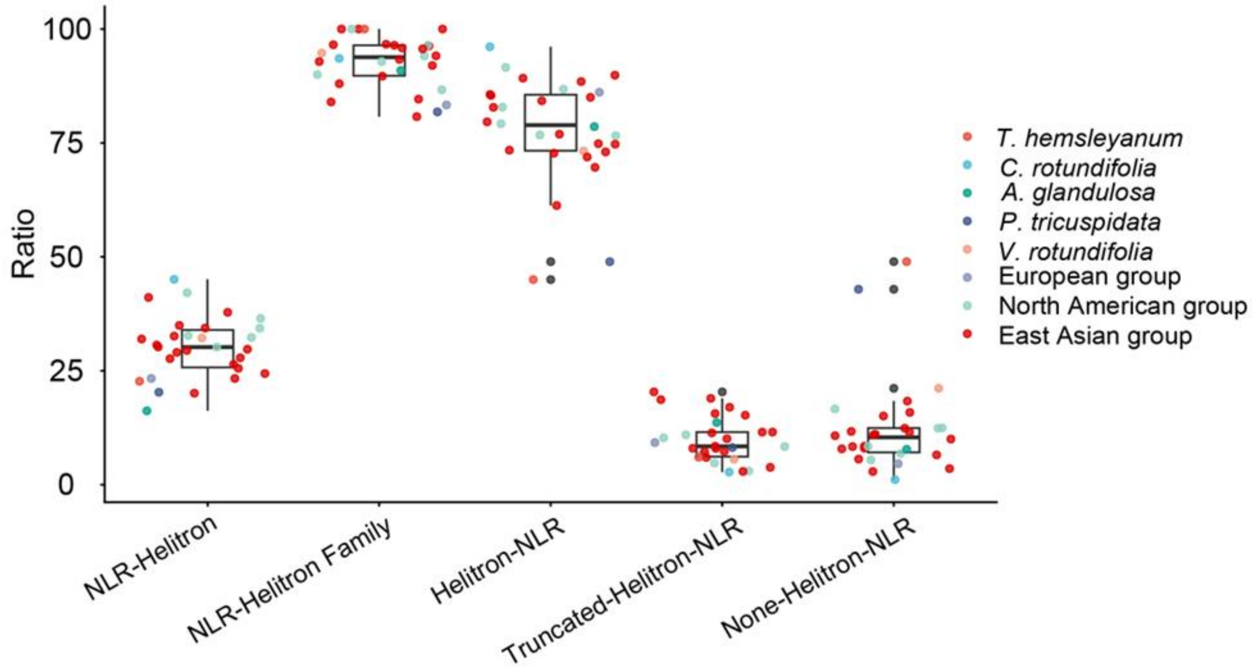


**Supplementary Fig. 34 |** The positional relationship between NLR and Helitron. NLR-Helitron: Helitrons surrounded by NLRs. NLR-Helitron family: Types of Helitrons surrounded by NLRs. Helitron-NLR: NLRs surrounded by Helitrons. Truncated-Helitron-NLR: NLRs surrounded by Helitron fragments. None-Helitron-NLR: NLRs with no identified Helitrons or Helitron fragments nearby.

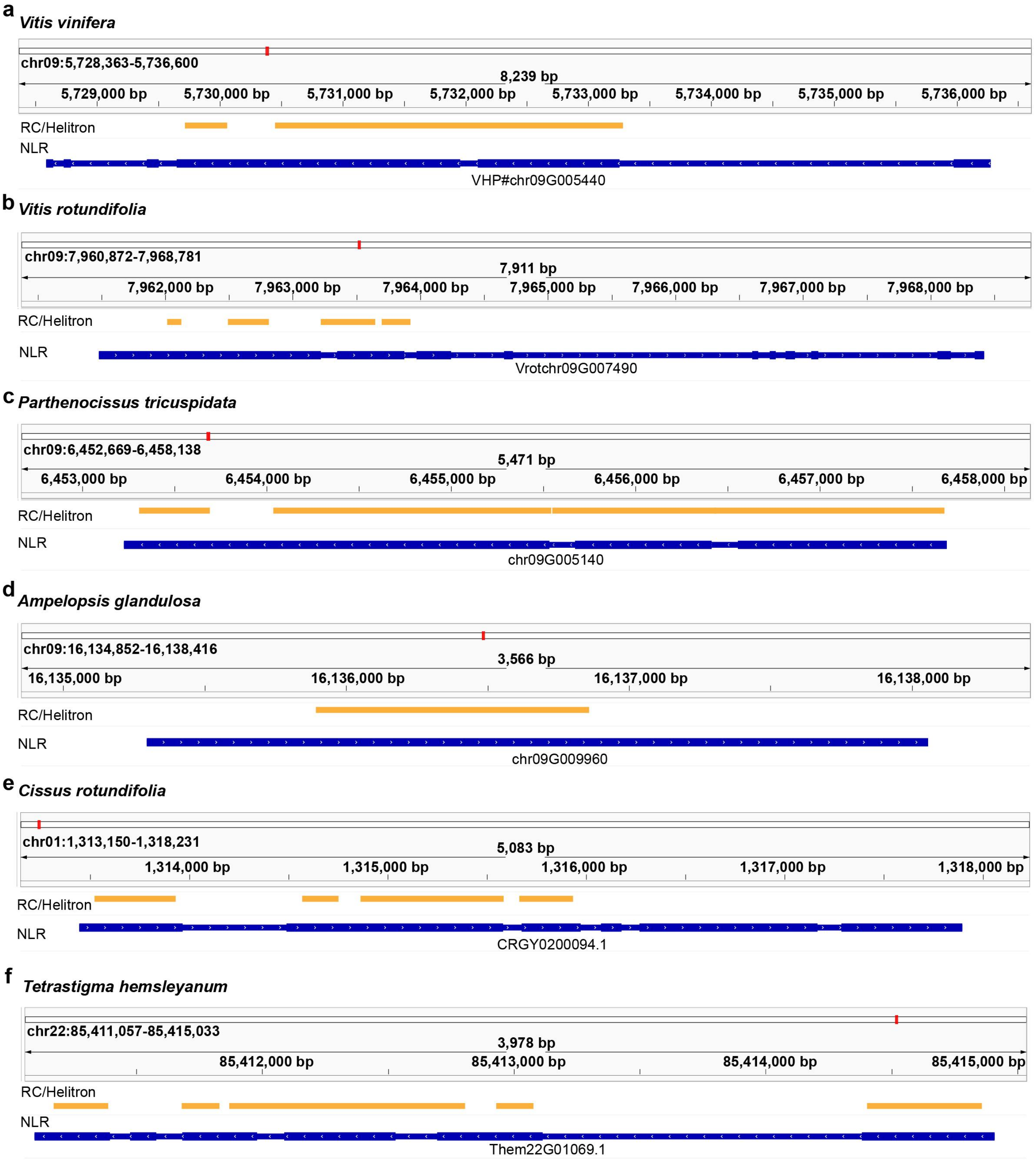


**Supplementary Fig. 35 |** Helitron elements carrying NLR coding sequences in six representative Vitaceae species: *V. vinifera*(a), *V. rotundifolia*(b), *P. tricuspidata*(c), *A. glandulosa*(d), *C. rotundifolia*(e), and *T. hemsleyanum*(f)

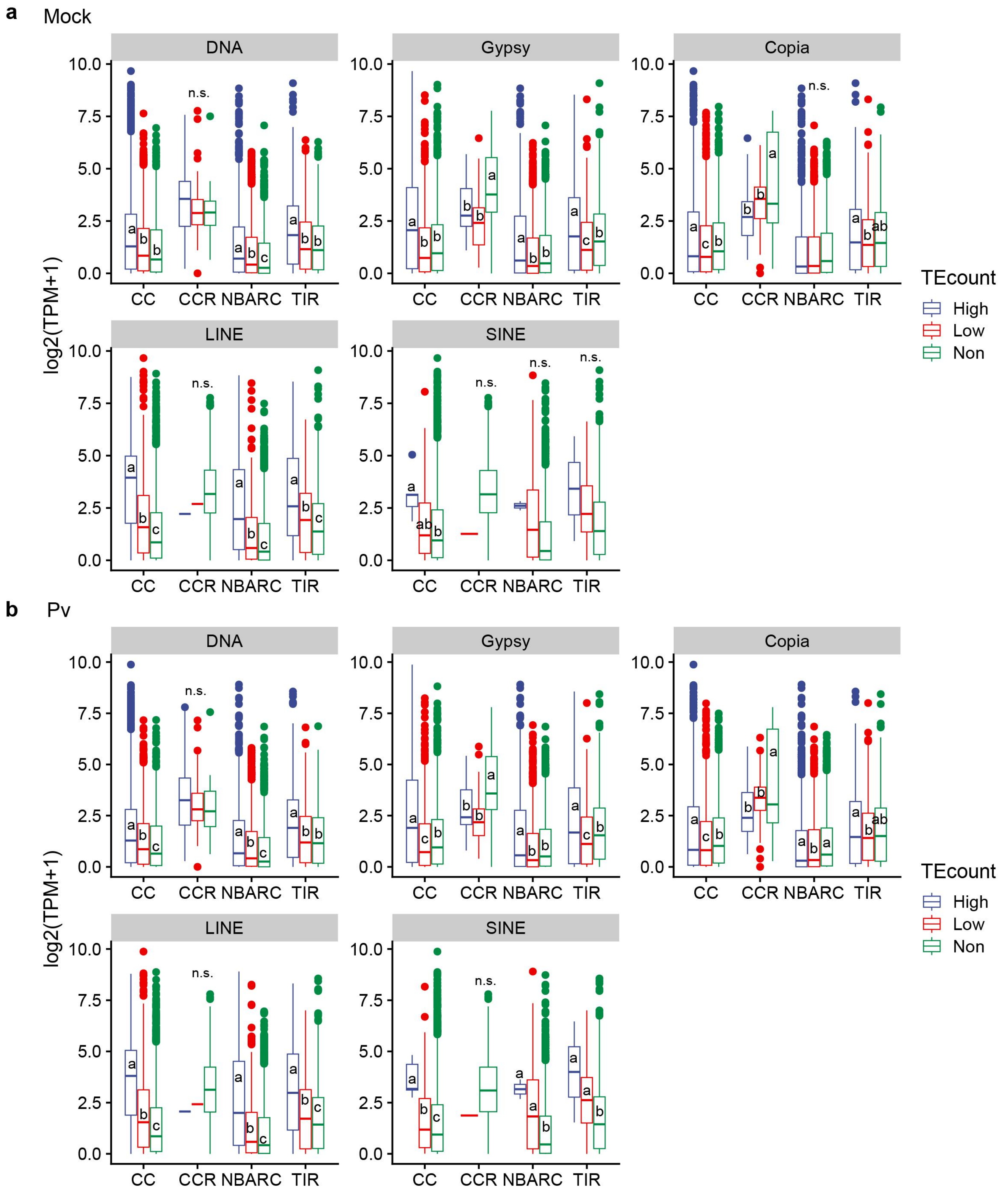


**Supplementary Fig. 36 | a.** The impact of TE frequency within the 2 kb upstream and downstream regions of NLR genes on the expression levels of different NLRs in 27 *Vitis* samples (Mock). The boxplot is generated using the expression levels of a single NLR gene from 25 different species. (n (High/Low/Non): for DNA, CC=2908/2612/1832, CCR=108/36/9, NBARC=1725/1732/1160, TIR=1066/793/494; for Gypsy, CC=577/1407/5368, CCR=17/46/90, NBARC=423/1219/2975, TIR=254/582/1467; for Copia, CC=1077/2110/4165, CCR=41/56/56, NBARC=806/1430/2381, TIR=385/670/1248; for LINE, CC=176/557/6619, CCR=1/1/151, NBARC=117/438/4062, TIR=69/177/2057; for SINE, CC=5/93/7254, CCR=NA/1/152, NBARC=2/47/4568, TIR=2/59/2242.). **b.** The impact of DNA, Gypsy, Copia, LINE and SINE frequency within the 2 kb upstream and downstream regions of NLR genes on the expression levels of different NLRs in 25*Vitis* samples (*Pv* infection). The boxplot is generated using the expression levels of a single NLR gene from 25 different species. (n (High/Low/Non): for DNA, CC=2908/2612/1832, CCR=108/36/9, NBARC=1725/1732/1160, TIR=1066/793/494; for Gypsy, CC=577/1407/5368, CCR=17/46/90, NBARC=423/1219/2975, TIR=254/582/1467; for Copia, CC=1077/2110/4165,

CCR=41/56/56, NBARC=806/1430/2381, TIR=385/670/1248; for LINE, CC=176/557/6619, CCR=1/1/151, NBARC=117/438/4062, TIR=69/177/2057; for SINE, CC=5/93/7254, CCR=NA/1/152, NBARC=2/47/4568, TIR=2/59/2242.)

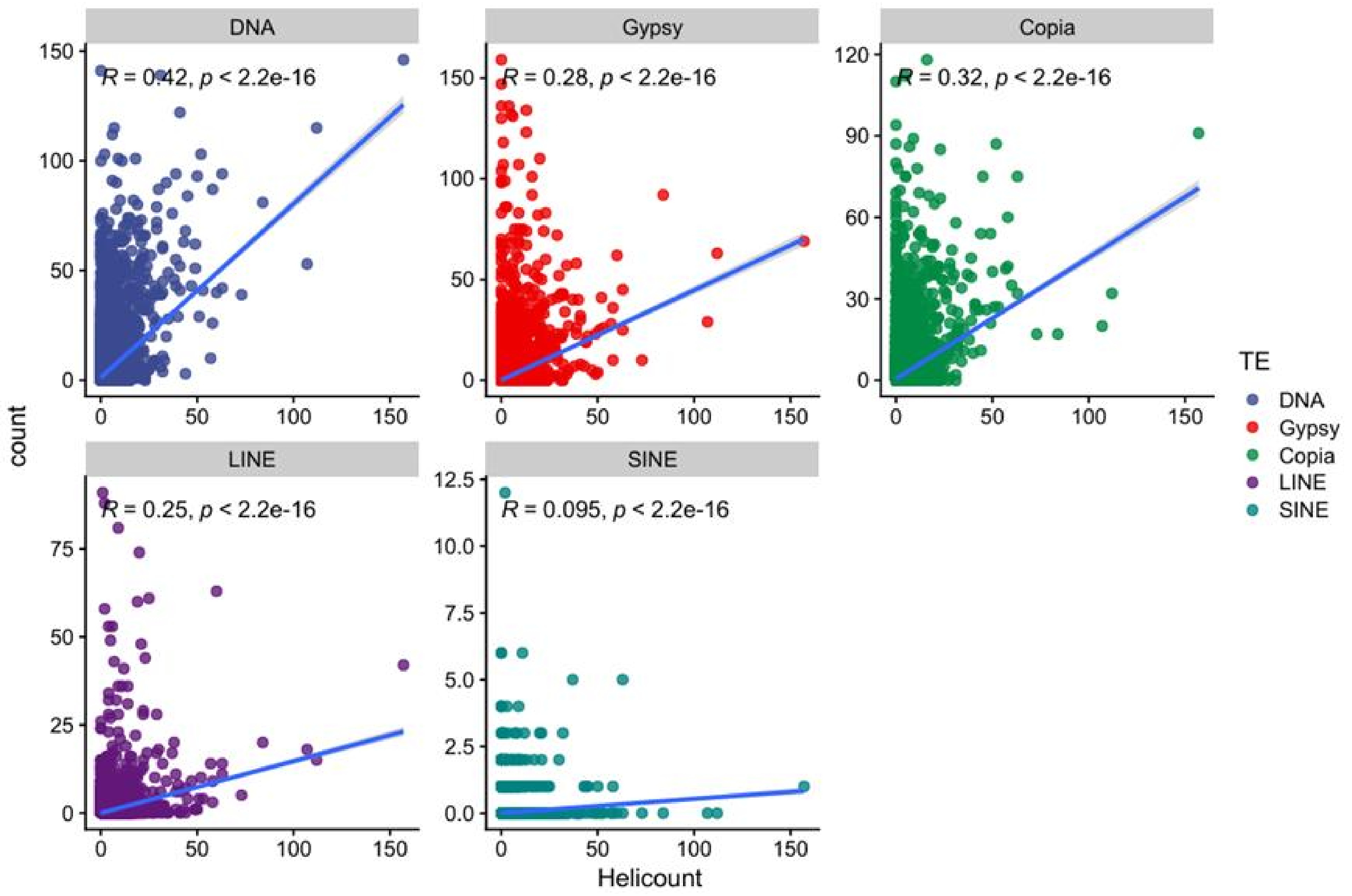


**Supplementary Fig. 37 |** The correlation between the number of helitrons and the number of other five types of TEs within the 2 kb upstream and downstream regions of NLR genes.

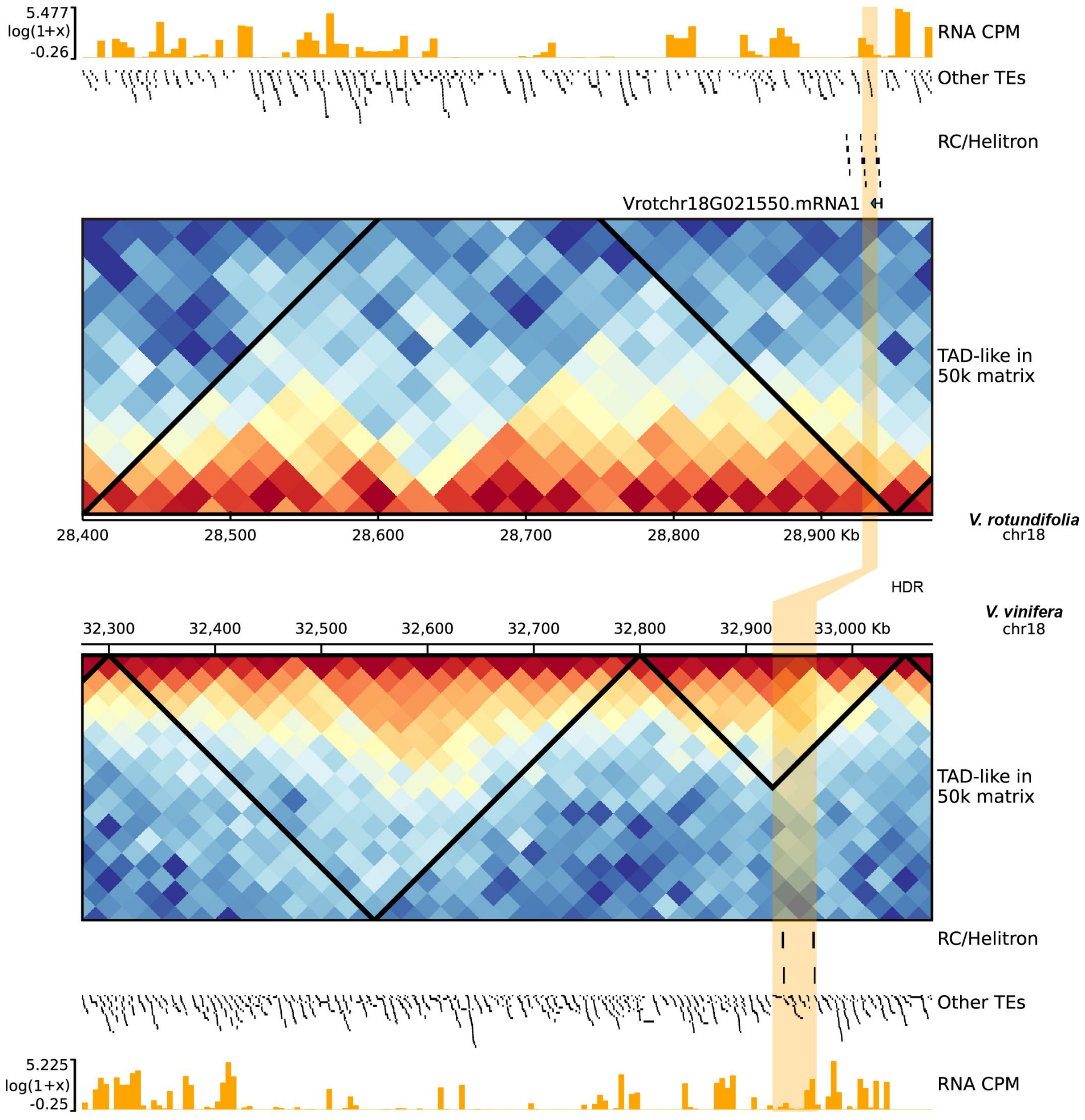


**Supplementary Fig. 38 |** The TAD landscape of the *Rpv-Vbel1* gene in the *V. rotundifolia* and *V. vinifera* genomes. The figure is divided into two parts: the upper part represents the *V. rotundifolia* genome information, and the lower part represents the *V. vinifera* genome information. The connecting bands between the two regions indicate high variability areas. In the TAD track, the triangular areas formed by the broken lines represent TAD-like regions.

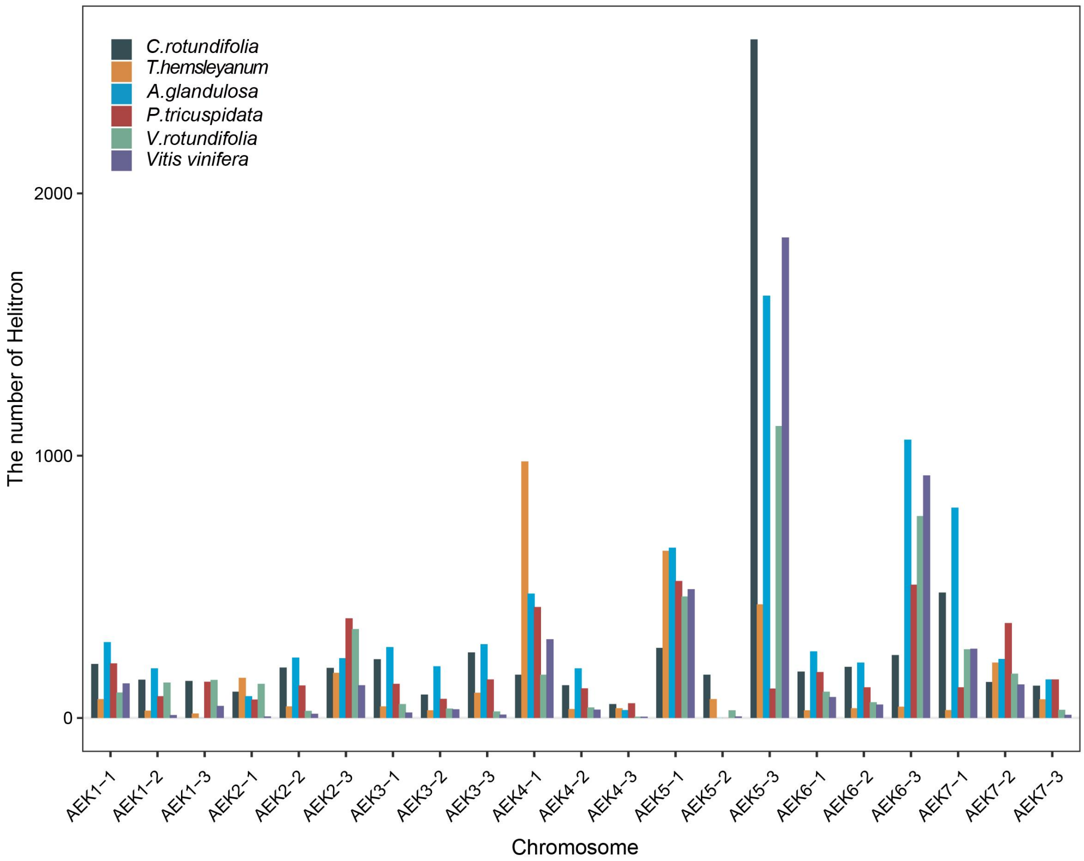


**Supplementary Fig. 39 |** The distribution of Helitron on ancestral chromosomal segments in 6 Vitaceae species. The bar plot shows the number of Helitron transposable elements identified on ancestral chromosomal segments (ACEK-derived segments) across six Vitaceae species. Each bar represents the Helitron count associated with a specific ancestral chromosome segment, and colors indicate different species as shown in the legend.
